# Structural Validation of the Intermediate Leptomeningeal Layer in the Human Central Nervous System

**DOI:** 10.1101/2025.07.14.664673

**Authors:** Ashutosh Kumar, Sanjeev Kumar Paikra, Rajesh Kumar, Ravi K. Narayan, Rakesh K. Jha, Shreyas Santosh Yadav, Pankaj Kumar, Arthi Ganapathy, Monika Anant, Shreekant Bharti, Nitesh Kumar, Vikas Pareek, S.N. Pandey, Shashank Shekhar, Mahboobul Haque, Tapas Chandra Nag, Tara Sankar Roy, Ashok Kumar Datusalia, Ashok Kumar Rastogi, Chiman Kumari, Adil Asghar

## Abstract

Traditionally, the human central nervous system (CNS) is described as having three meningeal layers, from outer to inner: dura mater, arachnoid mater, and pia mater. The arachnoid and pia mater are called the leptomeninges, and the space between them is filled with cerebrospinal fluid (CSF). Using gross dissection, light microscopy, and ultrastructural analysis of fresh postmortem and cadaveric CNS specimens spanning fetal to adult ages (N=61), we demonstrate a fibrocellular intermediate leptomeningeal layer (ILL) from the cortex to the caudal end of the spinal cord. The ILL divides the subarachnoid space (SAS) into two distinct structural compartments, through which vessels and nerves pass. The ILL shows unique structural features, such as dips into the brain’s sulci and fissures, as a double-fold membrane that bears intra-layer trabeculae, carries vessels, and forms the perivascular sheath. Moreover, throughout the CNS, it appears to be a non-sieved barrier, characterized by the presence of tight and adherens junctions. ILL, predominantly in the spinal cord, contains macrophage-like cells, indicating its layer-specific immune properties. The ILL warrants recognition as a distinct human meningeal layer with potential barrier and immune functions.

**Significance:** *An Intermediate Leptomeningeal Layer encloses the Central Nervous System in Humans:* The integrated analysis of our macroscopic, microscopic, and ultrastructural study provides robust support for an intermediate leptomeningeal layer (ILL) in the subarachnoid space (SAS) of the human central nervous system (CNS) along the entire neural axis. The ILL is a fibrocellular macroscopic structure, with restricted permeability, that divides the cerebrospinal fluid (CSF)-filled SAS into two distinct structural compartments. Uniquely, ILL revealed the presence of cells with macrophage-like properties, suggesting a possible role in immune surveillance. The ILL may redefine the established concept of protective coverings of CNS, CSF circulation dynamics, and the role of leptomeninges in health and disease, including drug delivery.

## Introduction

The traditional description of the meningeal arrangement around the neural axis in the human central nervous system (CNS) identifies three layers, from outer to inner—dura, arachnoid, and pia mater, with the latter two referred to as the leptomeninges (*1*). A few recent studies have reported a new leptomeningeal layer—the ‘subarachnoid lymphatic-like membrane (SLYM)’—between the arachnoid and pia of mouse brains, dividing the subarachnoid space into two compartments (*2*, *3*). Further, Rasmussen et al., 2024, indicated the presence of this layer in the meningeal coverings of the trigeminal ganglion in mice (*4*).

This layer has been described as a mesothelial layer that prevents the passage of particles larger than 1 μm in diameter and 3 kilodaltons in mass (*2*). This layer also hosts immune cells, dendritic cells and macrophages, whose numbers increase under inflammation-promoting conditions (aging, toxin exposure, etc), suggesting an immunogenic barrier function (*2*).

No prior study has detailed the macroscopic anatomy or established a nomenclature for an additional meningeal layer within the existing leptomeningeal framework. A rare report by Nicholas and Weller (1988) (*5*), first described a sieved “intermediate leptomeningeal layer (ILL)” in the human spinal cord (*5*)(*6*). As the term ‘ILL’ reflects the anatomical position of this layer between the arachnoid and pia, we consider it equivalent to the ‘SLYM’ described in recent mouse studies (2, 3) and adopt this terminology in the present study.

In our recent systematic review of historical literature, we demonstrated that previous studies in both animals and humans have also described a structure resembling the ILL in either the brain or spinal cord (*6*). Furthermore, recent radiological imaging and advanced microscopic studies have suggested the presence of a perivascular sheath enclosing cerebrospinal fluid (CSF) around cerebral vessels, which opens into the cisterns (*7–9*). Some authors have proposed that this CSF-filled perivascular sheath, located within the SAS and communicating with the cisterns, may have been created by the ILL (*7–10*).

Mollgård et al. (2023) and Pla et al. (2024) demonstrated a Podoplanin⁺ Prox-1⁺ Lyve-1⁻ Vegr-3⁻ SLYM layer in mouse brains but did not show its presence below the brainstem (*2*, *3*). Mollgård et al. (2023) also presented limited human data showing immunohistochemical expression of SLYM-specific markers in the frontal cortex (*2*). Notably, they showed matching marker expression in both mice and humans. We believe that the presented human data requires further validation in a larger sample size to prove the presence of such a layer, especially to describe its extent along the entire CNS.

Here, we examined the ILL in fresh postmortem and cadaveric human CNS specimens along the entire neural axis using gross dissection, brightfield and fluorescence light microscopy with histological and immunophenotypic characterization, and ultrastructural methods.

## Results

**Macroscopic observations:** (Fig.1A-D), Supplementary movies 1-8: https://doi.org/10.6084/m9.figshare.29375426

**Figure 1.**
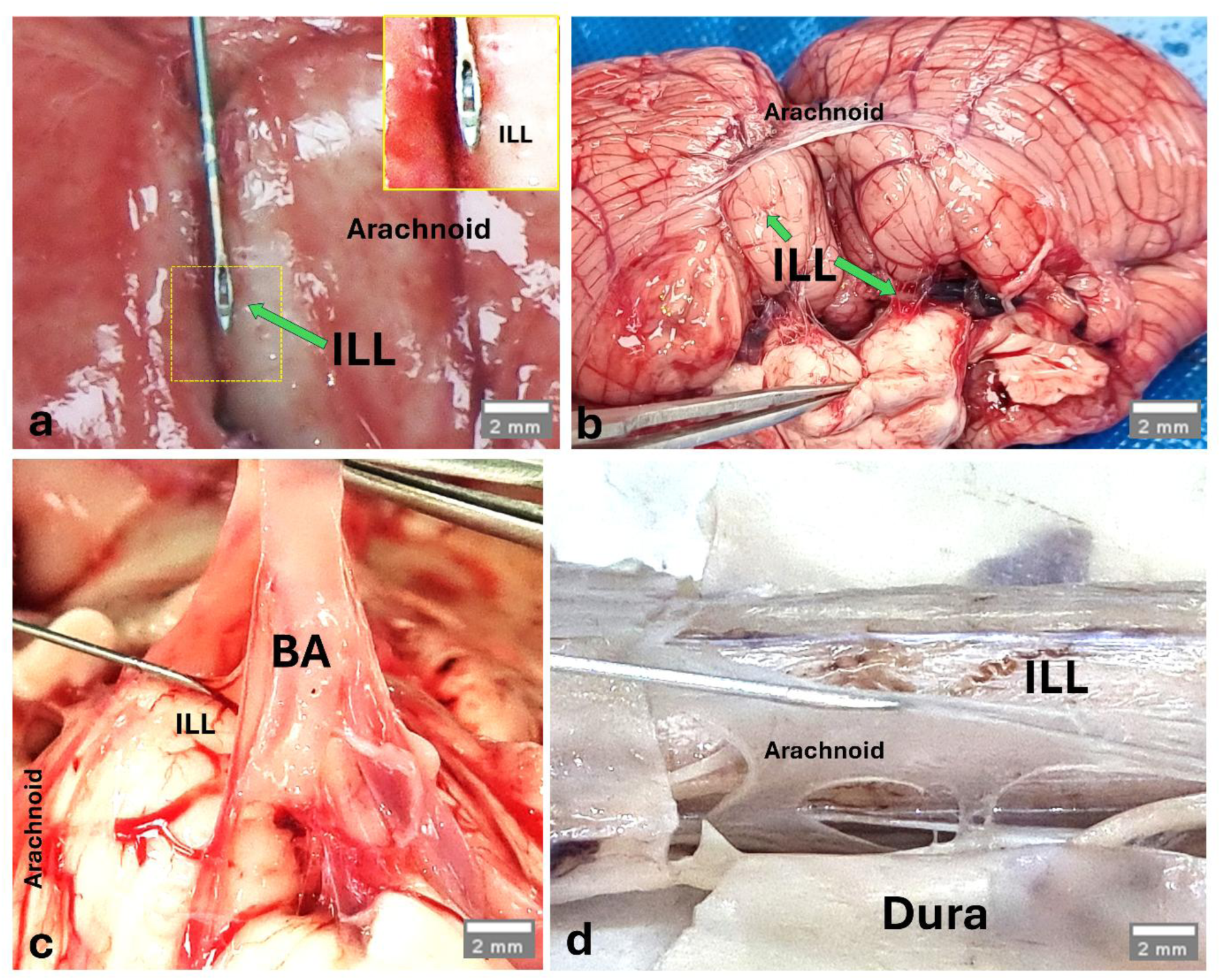
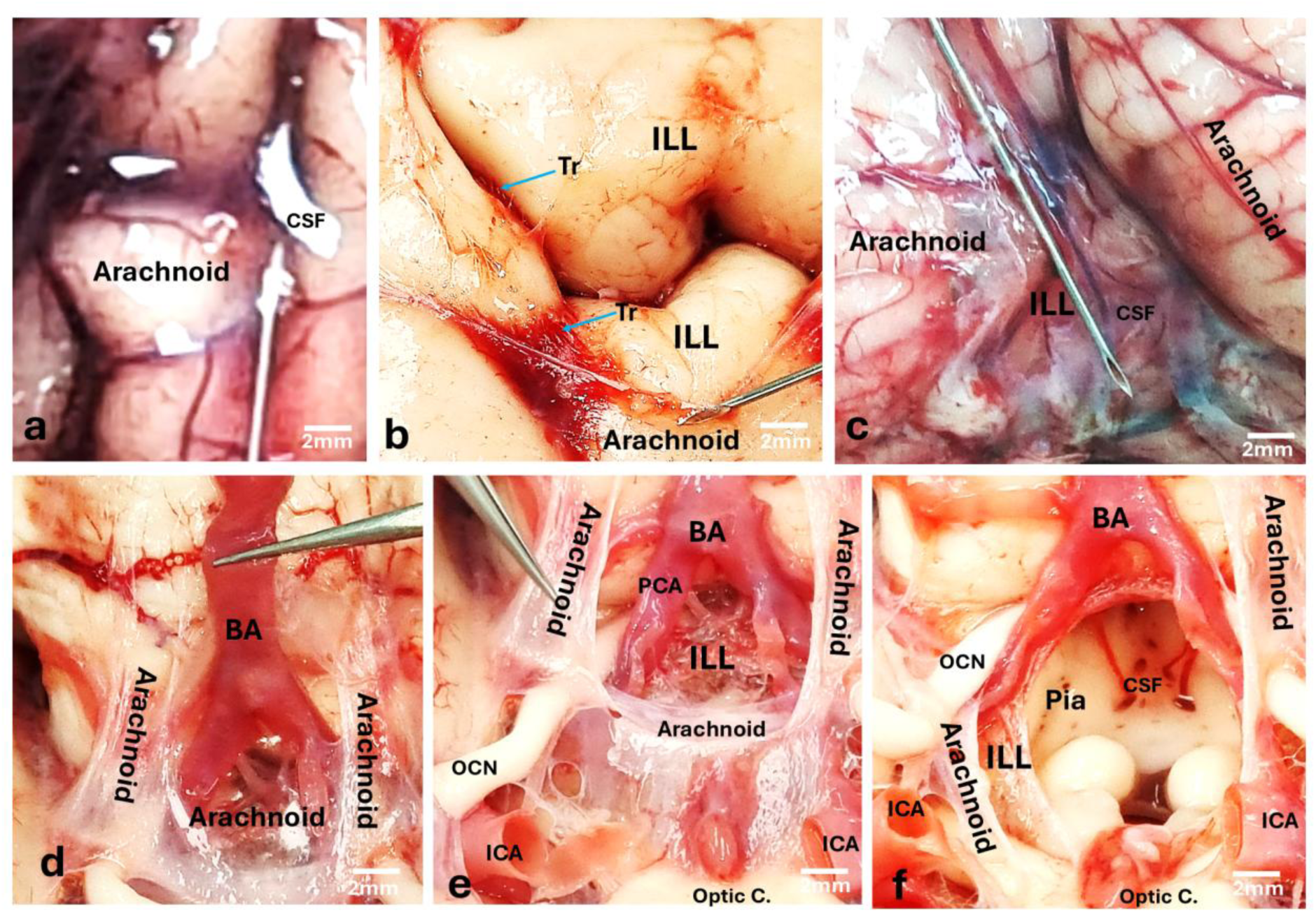
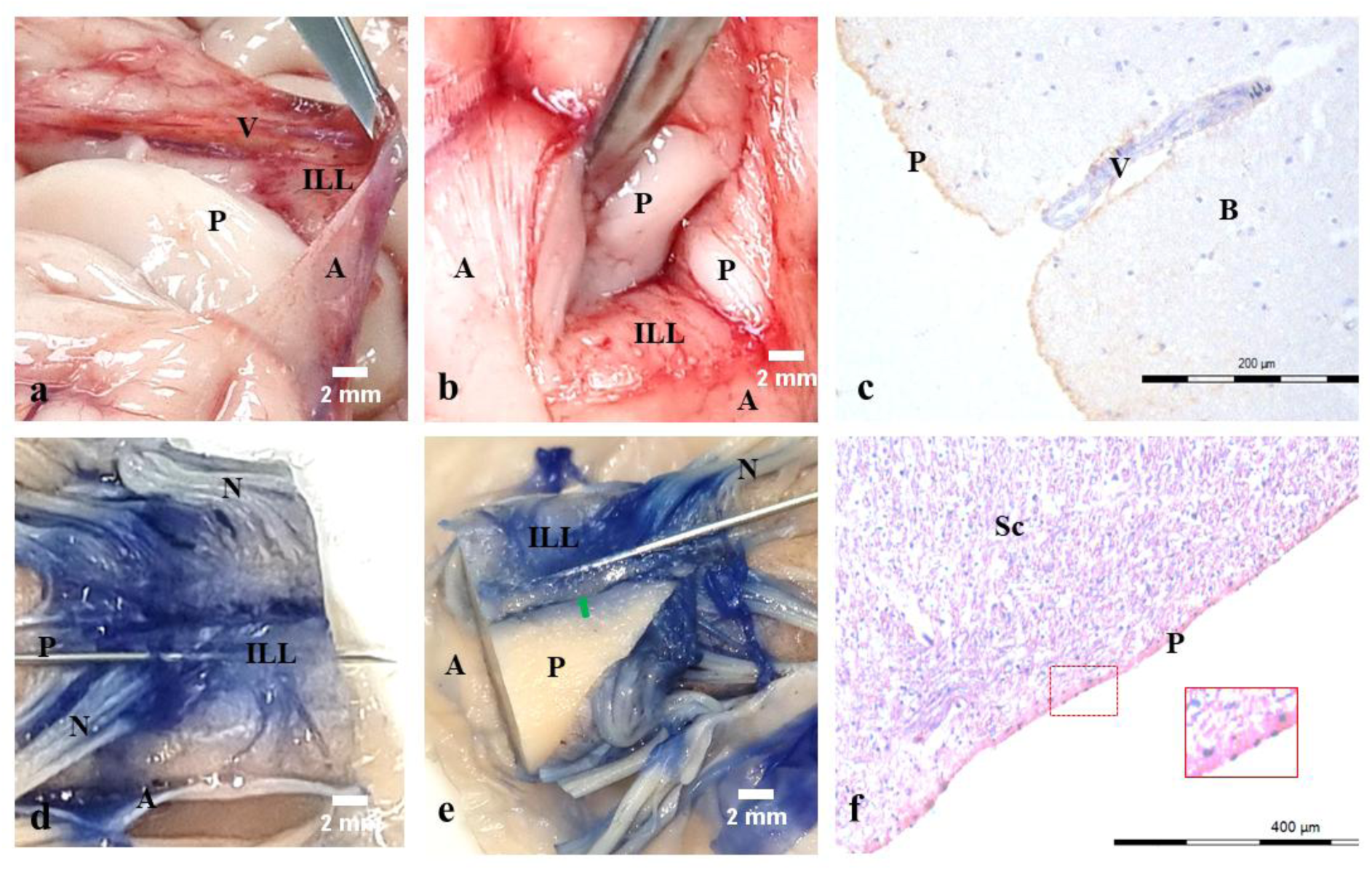
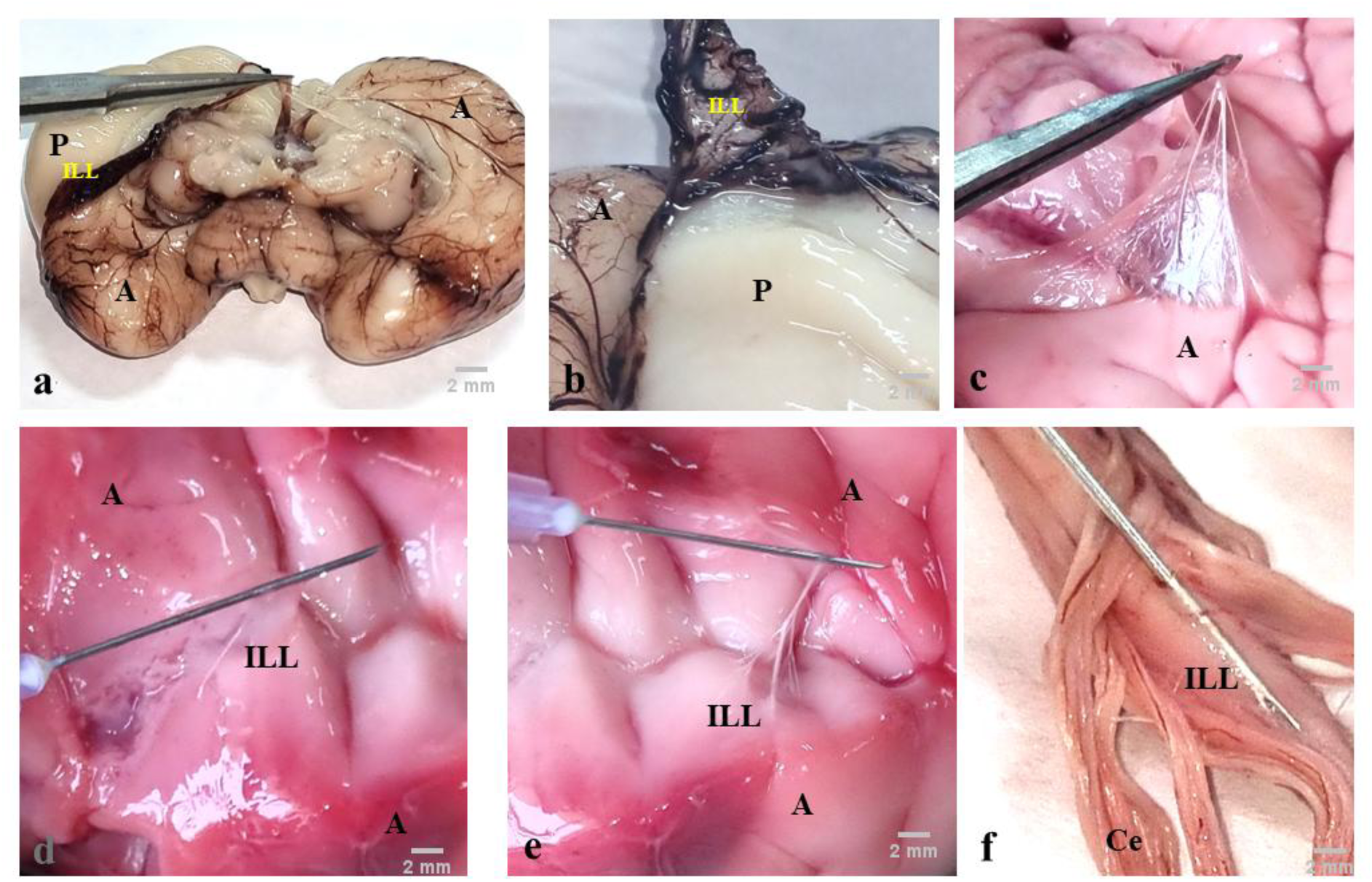
Disposition of Meningeal Layers in the Human Brain and Spinal Cord. (Still images from dissection videography of the subarachnoid space around the human brain and spinal cord in adult (A-C) and fetal (D) postmortem and cadaveric specimens.) **A. a. Cerebral cortex**: The dura mater was removed during autopsy. Leptomeningeal layers are shown near the cortical sulcus. The outermost leptomeningeal layer—the arachnoid mater—was incised and reflected. Beneath the arachnoid, another leptomeningeal layer, the ILL is visible (marked by a green arrow and lifted over the needle, also shown in the *inset*), entering the sulcus. Compared to the innermost leptomeningeal layer, the pia mater (not shown here), the ILL was loosely adherent to the cortical surface and easily separable (as demonstrated by lifting with the needle). **b. Cerebellum**: The arachnoid mater was incised and reflected from the cisterna magna region. A similar layer was visible covering the floor of the cisterna magna (small green arrow); this is the ILL. As the ILL extended from the cerebellar cortex, it formed the inferior medullary velum at the lower borders of the fourth ventricle (larger green arrow) and continued over the spinal cord. **c. Brainstem**: The arachnoid mater was incised over the pons and reflected laterally, revealing the basilar artery (BA) in the midline sulcus. A shiny layer, identified as the ILL, was visible covering the brainstem and passing beneath the basilar artery. Compared with the pia mater (not shown here), the ILL was loosely adherent to the surface and easily separable (shown lifted by the needle). Vessels can be seen passing beneath the ILL. **d. Spinal cord**: The dura mater and arachnoid mater were incised and reflected laterally. Beneath the arachnoid, a transparent and loose layer, the ILL was visible. Tortuous vessels could be seen passing beneath the ILL. **B. a–b. Cerebral cortex**: The dura mater was removed during autopsy as per standard protocol. Leptomeningeal layers are shown near a cortical sulcus. The outermost leptomeningeal layer, the arachnoid mater, was punctured with a needle to access the SAS, which is filled with CSF. The arachnoid mater was then incised and reflected laterally. Beneath it lies the ILL, which is seen entering the sulcus. Compared to the innermost leptomeningeal layer, the pia mater (not shown here), the ILL is loosely adherent to the cortical surface and easily separable. The ILL divides the CSF-filled SAS into two compartments: the outer SAS, which chiefly contains CSF, and the inner SAS, which forms a thin layer over the smooth cortical surface. ILL layers entering the sulcus are connected by dense ‘intralayer trabeculae’ (Tr, blue arrows). **c. Cerebellum**: The arachnoid mater was incised and reflected from the cisterna magna region. A shiny, transparent layer, identified as the ILL, is observed covering the floor of the cisterna magna (lifted with the needle). A CSF-filled space containing vessels is visible beneath the ILL. **d–f. Interpeduncular fossa**: The arachnoid mater completely covers the interpeduncular fossa and extends over the optic chiasma. The basilar artery is visible passing beneath the arachnoid mater to form the Circle of Willis (d). The lateral and anterior margins of the arachnoid mater are tough and fibrous. The arachnoid mater was partially incised (with margins left intact) and reflected to expose the Circle of Willis and its major branches lying over another meningeal layer—the ILL (e). During autopsy, the CSF was drained from the outer SAS. The ILL was then incised and reflected to reveal the pia-covered neural surface (f). In the cisternal regions, the inner SAS is slightly deeper than over the cortical surface. A scant amount of CSF is visible in this inner compartment, and fine vascular branches from the Circle of Willis can be seen piercing the pia-covered surface. **C. a–c. Cerebral Cortex:** The dura mater was removed during autopsy as per standard protocol. (a) The arachnoid mater (A) and ILL are separated *en bloc* to reveal the underlying pia mater (P), which is adherent to the cortical surface. Blood vessels (V) can be seen sandwiched between the arachnoid and the ILL. Leptomeningeal layers are shown near a cortical sulcus. (b) The outermost leptomeningeal layer, the arachnoid mater (A), was incised at a sulcus to reveal the ILL. A double fold of the ILL enters the depth of the sulcus. Stretching the sulcus separates the loosely adhered ILL to one side, revealing the cortical wall covered by pia mater (P). The pia mater is inseparable from the cortical surface in gross dissection. (c) The cortical tissue sample was later immunostained with Podoplanin, a mesothelial marker, confirming the presence of a single cell-layer-thick pia mater. **d–f. Spinal Cord** The dura mater was incised and removed (not shown here). Trypan blue dye was injected beneath the arachnoid mater, which was then incised and reflected to reveal the loosely adhered ILL (lifted over the needle). Removing the arachnoid also revealed spinal nerve rootlets between it and the ILL. The ILL showed a unique arrangement at the anterior median fissure, where it splits into superficial and deep layers at the sulcus margins. (d) The superficial layer of the ILL passes over the sulcus, bridging it, and forms a bed for the anterior spinal vessels that descend between the arachnoid and the ILL. (e) The ILL bridge over the fissure was incised and peeled off from the opposite side to reveal the deep layer—which enters into the depth of the fissure (green arrow), and the underlying pia-covered neural surface (P). (f) Hematoxylin and Eosin staining of the histological section confirmed the presence of the pia mater (P) (pial cells highlighted in the *inset*) over the neural surface after peeling off the ILL. **D. a–e. Cerebral Cortex**: Dissections were performed on a second-trimester (24-week) fetus (a–b) and a third-trimester, fully matured (40-week) fetus (c). The dura mater was removed during dissection as per standard protocol. In the earlier specimen, the meningeal layers were well developed; however, sulci had not yet formed. In the latter specimen, both fully developed meningeal layers and sulci were observed. (a–c): Based on a vascular pedicle, the arachnoid mater (A) and ILL were separated *en bloc*, revealing major subarachnoid vessels sandwiched between the two layers. In the 24-week fetus, since sulci had not yet developed, the arachnoid–ILL complex could be easily peeled off the cortical surface, exposing the underlying pia mater (P) adherent to the cortex (a–b). In the fully matured third-trimester fetus, the leptomeningeal arrangement resembled that of the adult brain (c-e). The arachnoid layer passed over the sulci, while the ILL dipped into them. The arachnoid was incised to expose the ILL within the sulcus, which extended to the floor (lifted over the needle, d). Subarachnoid vessels entered the sulcus between the folds of the ILL, and their fine branches pierced this layer to reach the space between it and the pia mater (a vascular pedicle of the ILL is being lifted over the needle to show vascular branches piercing it, e). **f. Spinal Cord**: A third-trimester, fully matured (40-week) fetus was dissected following standard protocol. The dura and arachnoid mater were incised and removed. In the lower sacral segments, spinal nerve rootlets forming the cauda equina (Ce) were observed lying between the arachnoid and the ILL. The ILL formed a thick tunic around the conus medullaris (lifted over the needle), which continued over the filum terminale. Abbreviations: D-dura, A-arachnoid, P-pia, ILL-intermediate leptomeningeal layer, Tr-trabeculae, N-nerve, V-vessel, BA-basilar artery, ICA-internal carotid artery, PCA-posterior cerebral artery, OCN-Oculomotor nerve, Optic c.-Optic chiasma, SAS-subarachnoid space, i-inner, o-outer. B-brain, Sc-spinal cord.

### ILL divides the SAS into two distinct CSF compartments

We observed a distinct ILL in the gross dissection of fresh postmortem and cadaveric brain and spinal cord specimens (Fig. 1A, Supplementary movie 1–4). Macroscopically, this layer appeared as a glistening membrane with intact epithelial surfaces. It was present along the entire neural axis from the cortex to the caudal end of the spinal cord. We also noted the presence of coarse and fine trabeculae separating the ILL from the arachnoid and pia, respectively (Fig. 1B, Supplementary movie 1-4).

In addition, the vascular networks and the cranial and spinal nerves between the leptomeningeal layers, as they traverse through the SAS, further distinguish the ILL from the other two layers (Fig. 1B-D, Supplementary movie 2–5). The ILL formed the perineural sheath as the cranial nerve or spinal nerve rootlets passed through this layer (Supplementary movie 2–5).

The ILL divided the SAS into a wide outer and a shallow inner CSF compartment. The CSF was primarily contained in the outer SAS (Fig. 1B, Supplementary movie 6). In the inner SAS, the CSF volume was scanty, except in the larger cisterns of the brain, such as the cisterna magna, where a noticeable volume of CSF was observed in some specimens (Supplementary movie 7). Notably, the inner CSF compartment was markedly reduced at the gyral surfaces (Fig. 1B). This observation was further substantiated in the quantitative estimation of CSF spaces in histological images (See result section: Morphometric and quantitative estimations: Outer vs. Inner SAS compartments, Fig. 7B).

### ILL thickness varies along the neural axis

The ILL appeared to be thicker than the pia but thinner than the arachnoid in the cortex. However, in the spinal cord, particularly in the lumbosacral region, the ILL was comparable in thickness to the arachnoid (Fig. 1A: d, C:d-f, Supplementary movie 4). This was confirmed by quantitative ILL thickness measurements in SEM images (see Morphometric Results: Comparative Thickness of ILL, Fig. 7A).

### ILL sinks into sulci and fissures

Unlike the arachnoid, which bridges over the sulci and fissures, the ILL descends into their depths. However, unlike the pia—which also descends but closely adheres to the walls—the ILL appears as a loosely attached, double-folded membrane that carries subarachnoid vessels into the sulci and fissures (Fig. 1C-D, Supplementary movie 1). In the spinal cord, the ILL dips into the major sulci/fissures. It thus can be easily differentiated from the arachnoid layer that bridges over them (Fig 1A: d. C: d-e, Supplementary movie 4, 8).

### ILL bears unique intra-layer trabeculae

Uniquely, fine intra-layer trabeculae were found between the opposing surfaces of the ILL, entering the sulci and fissures in the adult CNS. These fine trabeculae were densest at the entry of sulci and fissures and seemed to create a seal or diaphragm (Fig. 1B: b, Supplementary movie 1).

### Distinct compartmental organization of SAS vessels

We observed a distinct compartmental organization of vessels in the SAS. The outer SAS contained a coarse network of higher caliber vessels, while the inner SAS compartment contained a finer network of small caliber vessels in the brain (Fig. 1B-d-f, Supplementary movie 1-2, also see result section: Morphometric and quantitative estimations: Outer vs. Inner SAS compartments, Fig. 7B). A slightly varying vascular pattern was observed in the spinal cord, where, after having a brief course in the outer SAS near their point of origin, the anterior spinal and radicular spinal arteries entered the inner SAS. In the inner SAS, the spinal arteries ran longitudinally in the anterior median fissure. From there, they sent fine branches to create a network in the inner compartment (Supplementary movie 8).

### ILL development parallels other leptomeningeal layers

In the fetuses, dissections were performed from the second trimester (24 weeks) onwards (Fig. 1D: a–f). In second-trimester specimens, where sulci had not yet formed, the arachnoid–ILL complex peeled easily from the cortical surface, exposing the underlying pia mater adherent to the cortex (Fig. 1D: a–b).

In fully matured third-trimester fetuses (40 weeks), the leptomeningeal arrangement resembled that of the adult brain (Fig. 1D: c-e). The arachnoid layer passed over the sulci, while the ILL dipped into them and extended to the floor. Subarachnoid vessels entered the sulcus between the folds of the ILL, and their fine branches pierced this layer to reach the space between it and the pia mater.

In the spinal cord, the arachnoid–ILL complex was continued up to its terminal end. In the lower sacral segments, spinal nerve rootlets forming the cauda equina (CE) were observed lying between the arachnoid and the ILL. The ILL formed a thick tunic around the conus medullaris, which continued over the filum terminale (Fig. 1D: f).

### Brightfield and fluorescence light microscopic observations

Histological analyses, including unstained, routine, and specialized staining, corroborated the macroscopic findings (Fig. S1, 2A–E). In unstained thick spinal cord sections, transmitted light revealed the ILL delineating two CSF compartments (Fig. S1). Routine hematoxylin and eosin staining confirmed separation of leptomeningeal layers by CSF spaces with intervening vascular profiles (Fig. 2A: a–c, 2B: a–b), while in traumatic brain and spinal injury, extravasated blood and inflammatory cells remained compartmentalized by the ILL (Fig. 2A: d, 2B: c–d).

**Figure 2.**
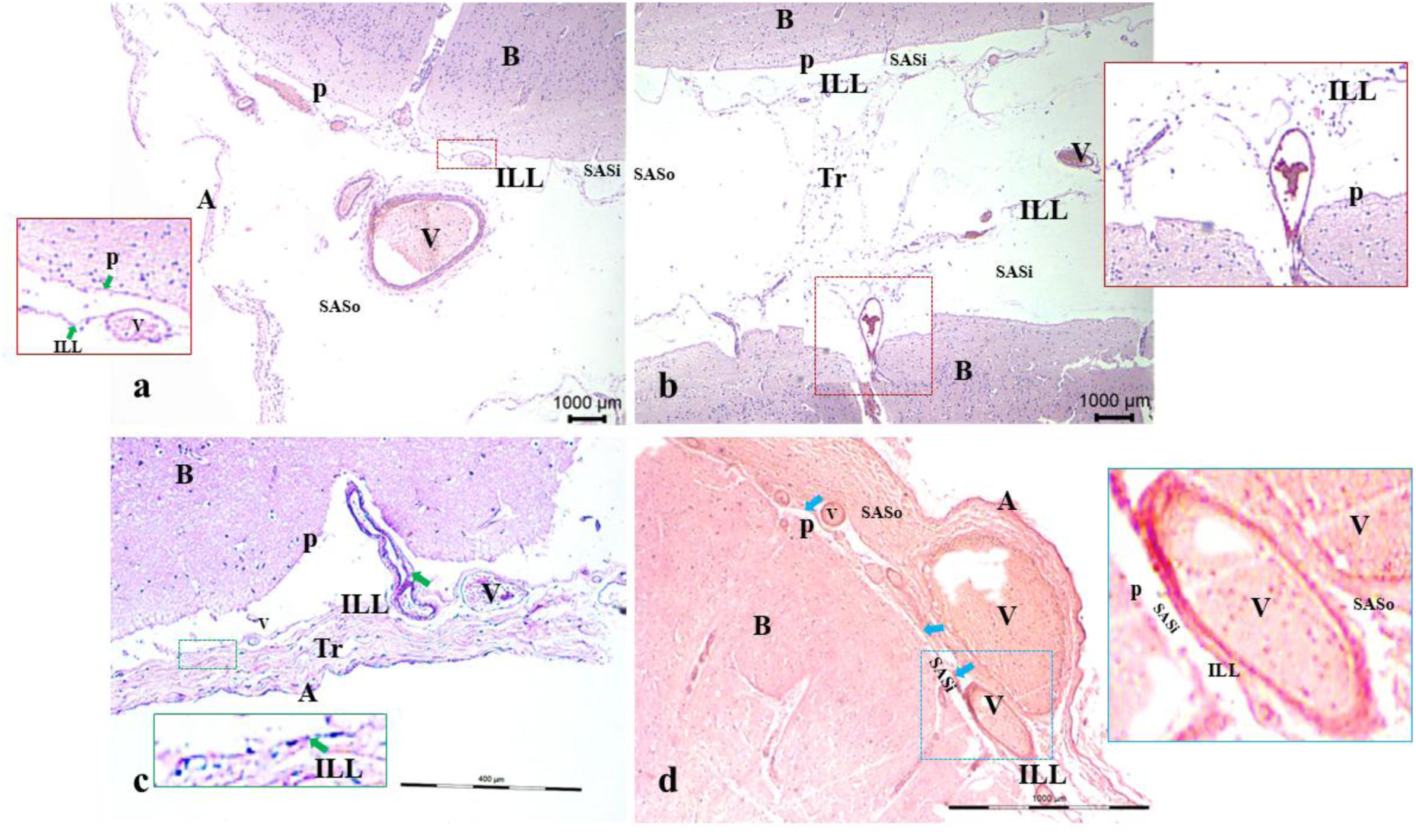
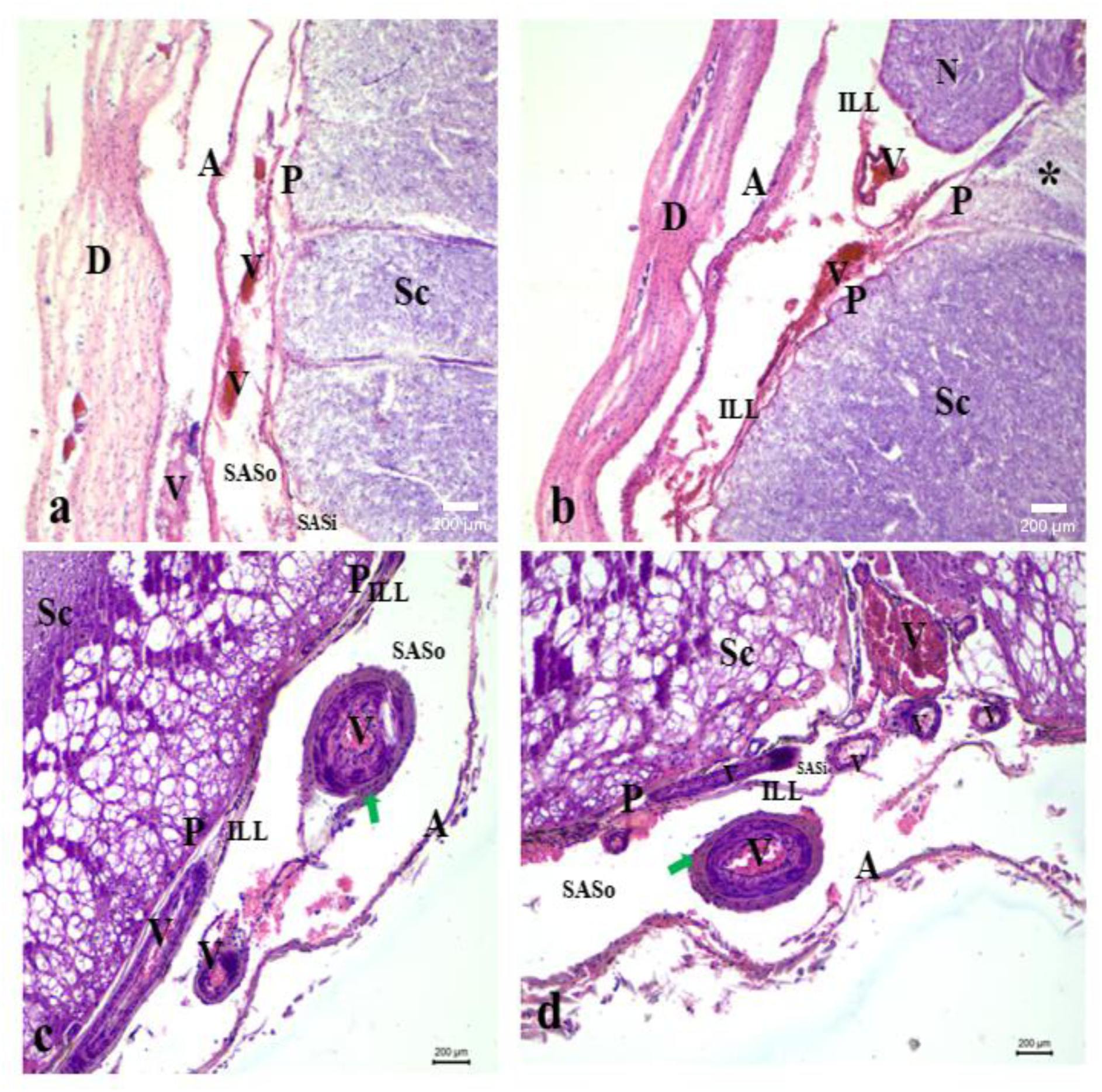
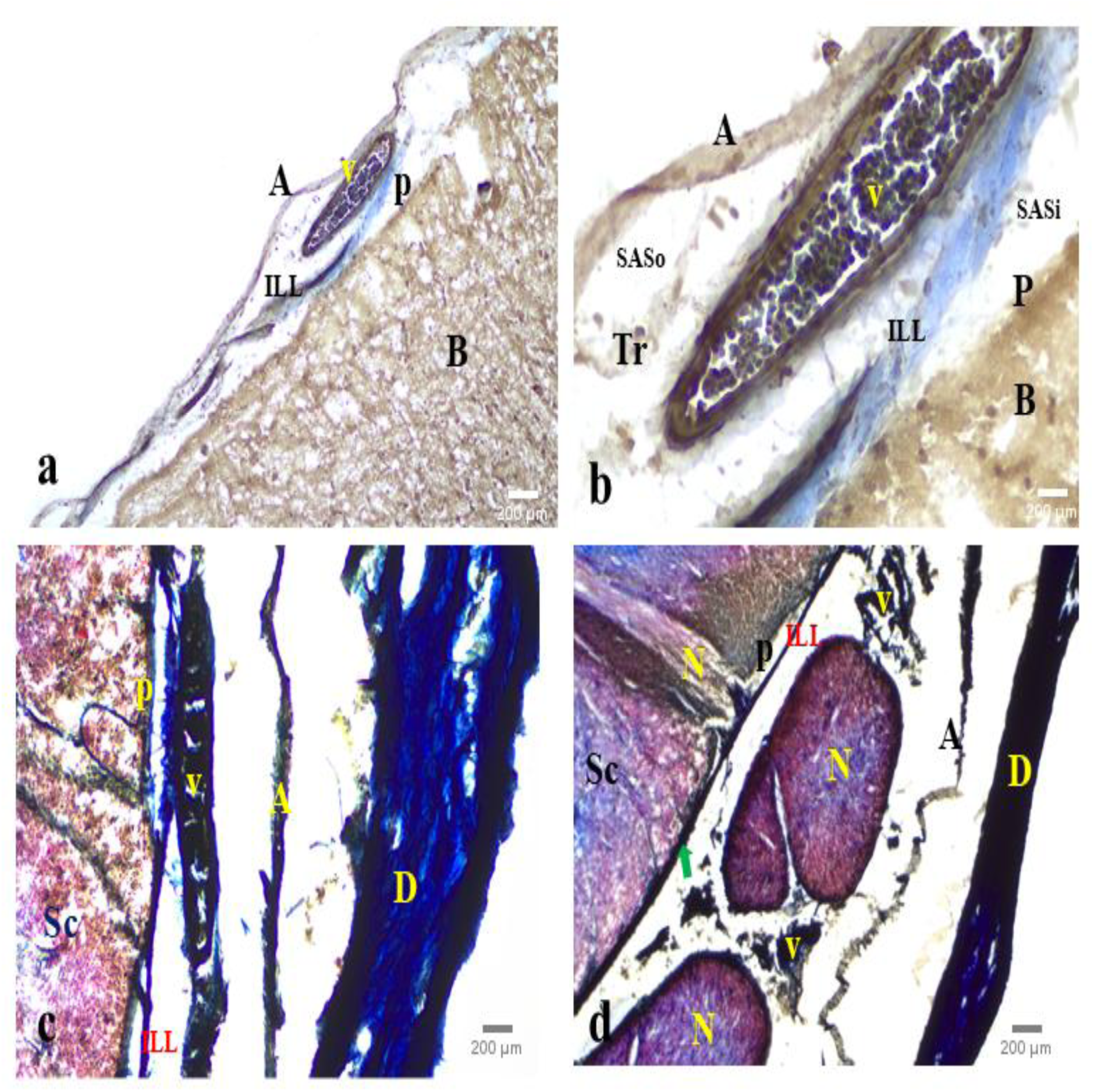
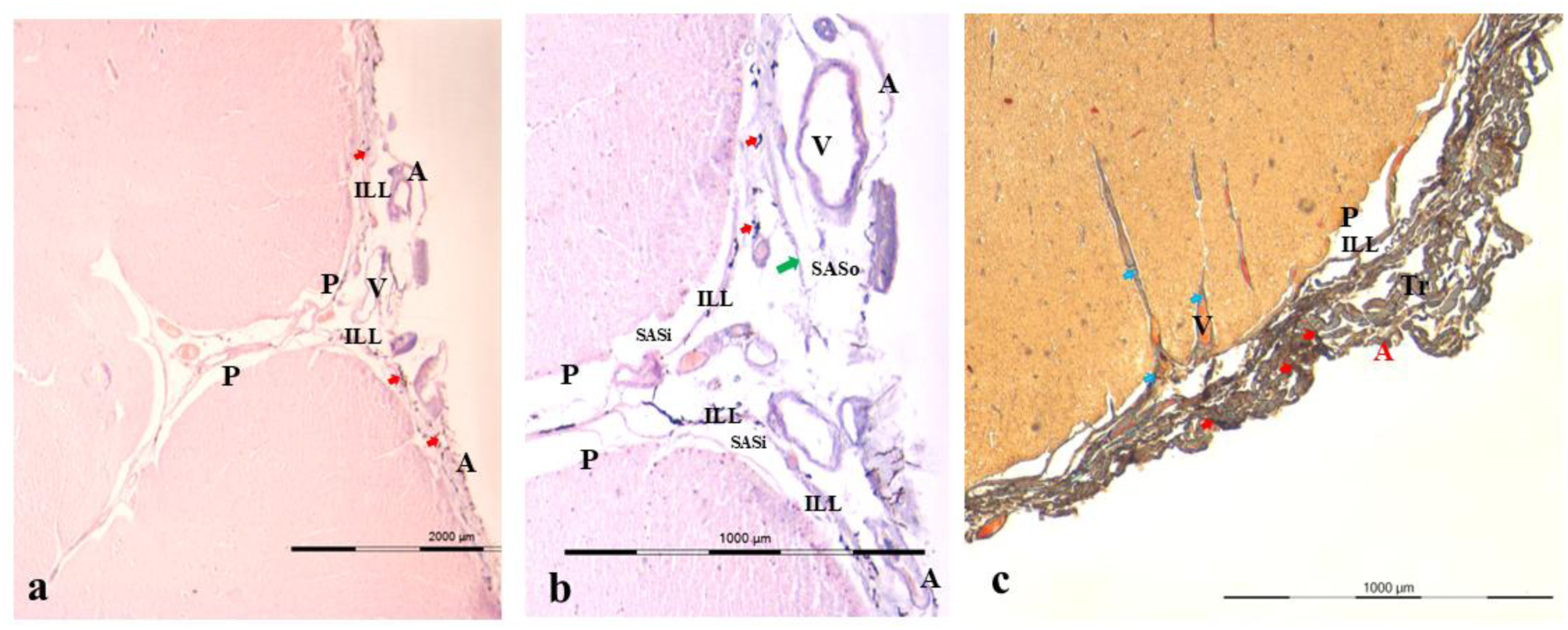
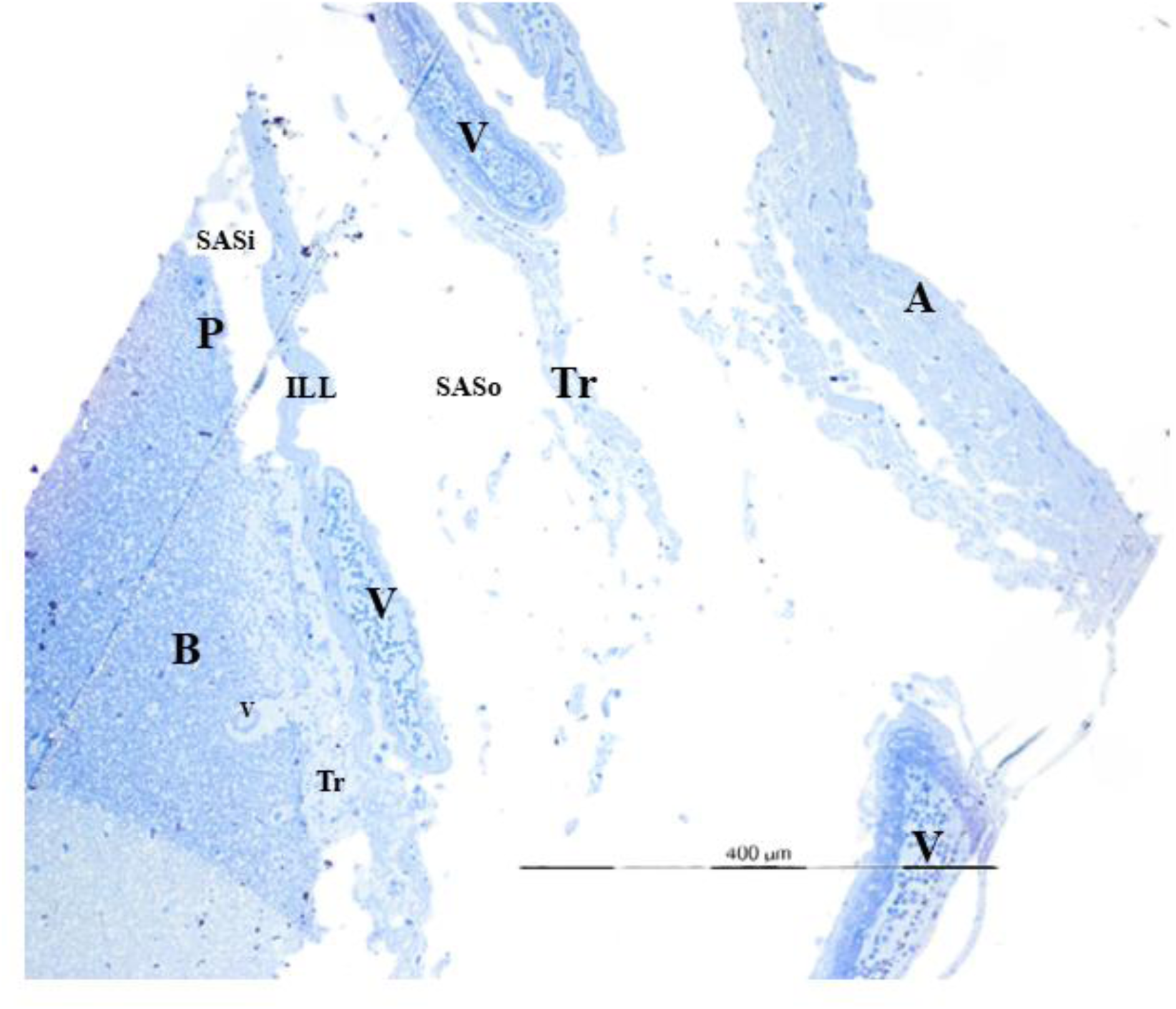
Histological features of the adult human cerebral cortex and spinal cord. **A. Hematoxylin and Eosin staining** **(a–c) Non-injured brain**: On the cortical surface of the brain (B), the intermediate leptomeningeal layer (ILL) is observed between the arachnoid mater (A) and the pia mater (P). The ILL divides the SAS into two compartments: the outer SAS (SASo) and the inner SAS (SASi), which is traversed by trabeculae (Tr). Vascular profiles in the SASo appear larger than those in the SASi, suggesting that major vascular branches are primarily confined to the SASo, while the SASi contains only a fine vascular network. A double fold of the ILL extends into the depths of the sulci, carrying an extension of the SASo (b). The ILL also forms a perivascular sheath around vessels (V) passing through it, predominantly arteries (c, green arrow). Insets showing an enlarged view of the cortical surface. **(d) Traumatic brain injury cases with subarachnoid hemorrhage**: The ILL (blue arrows) restricted the extravasated blood and inflammatory exudate from entering the SASi, suggesting a potential barrier function of the ILL. Inset showing an enlarged view of the cortical surface. **B. Spinal cord** **(a–b) Non-injury cases with dura mater (D) intact:** On the spinal surface (Sc), the ILL is visible between the arachnoid mater (A) and the pia mater (P). The ILL divides the SAS into outer (SASo) and inner (SASi) compartments. Vascular profiles (V) are seen in both compartments. A cross-section of a spinal nerve rootlet (N) is seen in SASo; the origin of the nerve rootlet from the spinal parenchyma (Sc) is marked with an asterisk. **(c–d) Traumatic spinal injury cases showing signs of vascular damage**: The dura mater was removed during autopsy. Vascular profiles are present in both CSF compartments. Vascular bleeding is predominantly confined to the SASo, and hemorrhage is also observed along the sulcal vessels. Arteries within the SASo exhibit a distinct perivascular sheath formed by the ILL (green arrows). **C. Masson’s Trichrome Staining** **(a–b) Non-injured brain:** On the cortical surface of the brain (B), the ILL is visible between the arachnoid mater (A) and the pia mater (P). The ILL divides the SAS into two compartments: the outer SAS (SASo) and the inner SAS (SASi), which are traversed by trabeculae (Tr). A large vascular profile is also visible within the SASo. Collagen bundles within the ILL are stained blue. **(c–d) Spinal Cord – Non-injury cases with dura mater (D) intact**: On the spinal surface (Sc), the ILL is visible between the arachnoid mater (A) and the pia mater (P). The arachnoid and ILL are separated by vessels (c) and spinal nerve (N) rootlets (d). Intense blue staining is observed in the dura, arachnoid, and ILL, indicating the presence of collagen bundles in these layers. The ILL is adherent to the pia mater at multiple loci (indicated by green arrow). **D. Hematoxylin and Eosin and Masson’s Trichrome staining following dye injection in SAS** (a-b) The section was taken through the anterior median sulcus after Trypan Blue dye was injected just beneath the arachnoid mater. On the spinal surface (Sc), the ILL is observed between the arachnoid mater (A) and the pia mater (P). A higher magnification of the anterior median fissure (b) shows that the ILL splits on both sides: one-layer bridges over the sulcus (green arrow, partially broken during histological processing), while the other enters its depth. This results in a double fold of ILL extending into the sulcus, enclosing vascular branches within. The ILL divides the SAS into outer (SASo) and inner (SASi) compartments. Injected dye particles are visible between the arachnoid and ILL and are confined to the SASo (red arrows). (c) Masson’s Trichrome staining of a section taken from the spinal surface reveals the fibrocellular structure of the ILL, showing morphological similarities to the arachnoid layer. Dense collagenous trabeculae (Tr) are observed between the two layers. Injected dye particles remain confined between the arachnoid and ILL within the SASo (red arrows). Double folds of the ILL enclosing vessels are seen entering small sulci (blue arrows). **E. Toluidine blue staining and light microscopic imaging of semithin sections of the subarachnoid space in the human brain** Semithin sections (1 µm) of SAS specimens from adult human brains were stained with Toluidine blue and examined under a bright-field light microscope. Between the outermost arachnoid (A) and the innermost pia mater (P), the ILL was observed. All three leptomeningeal layers were separated by trabeculae (Tr) and blood vessels (V). Abbreviations: D-dura, A-arachnoid, P-pia, ILL-intermediate leptomeningeal layer, Tr-trabeculae, N-nerve, V-vessel, SAS-subarachnoid space, i-internal, o-outer. B-brain, Sc-spinal cord.

Specialized staining further supported these observations: Masson’s trichrome demonstrated the fibrocellular composition of the ILL (Fig. 2C: a–d), and Trypan blue injected into the outer SAS remained confined between the arachnoid and ILL, indicating barrier function (Fig. 2D: a–c). This compartmental organization was also evident in semithin (1 μm) sections (Fig. 2E).

### A Podoplanin⁺ Prox1⁺ Lyve-1⁻ Vegfr-3⁻ layer divides the SAS into two CSF compartments

Immunohistological analyses using DAB and fluorescent markers identified a Podoplanin⁺ Prox1⁺ Lyve-1⁻ Vegfr-3⁻ layer (ILL) in the cortex and spinal cord that partitions the SAS into two CSF compartments (Fig. 3–4). Podoplanin was expressed in the ILL, pia, and inner trabeculae, but not in outer trabeculae (P < 0.001), thereby distinguishing the two compartments (Fig. 3A, S2:a-b). Prox-1 expression was specific to the ILL (F = 140, P < 0.001) and absent in the arachnoid, pia, and trabeculae (P < 0.001) (Figs. 3B–C, S2:c-e), with significant nuclear localization (Fig. 3D, Costes P = 1). Lyve-1 and Vegfr-3 were expressed by none of the leptomeningeal layers or trabeculae (Fig. 3: D-E), except by a unique cell population in ILL, predominantly in the spinal cord, which also expressed typical macrophage markers, and were indicated to be resident or infiltrating macrophages (See further results in section “Macrophage-like cells in the ILL”).

**Figure 3.**
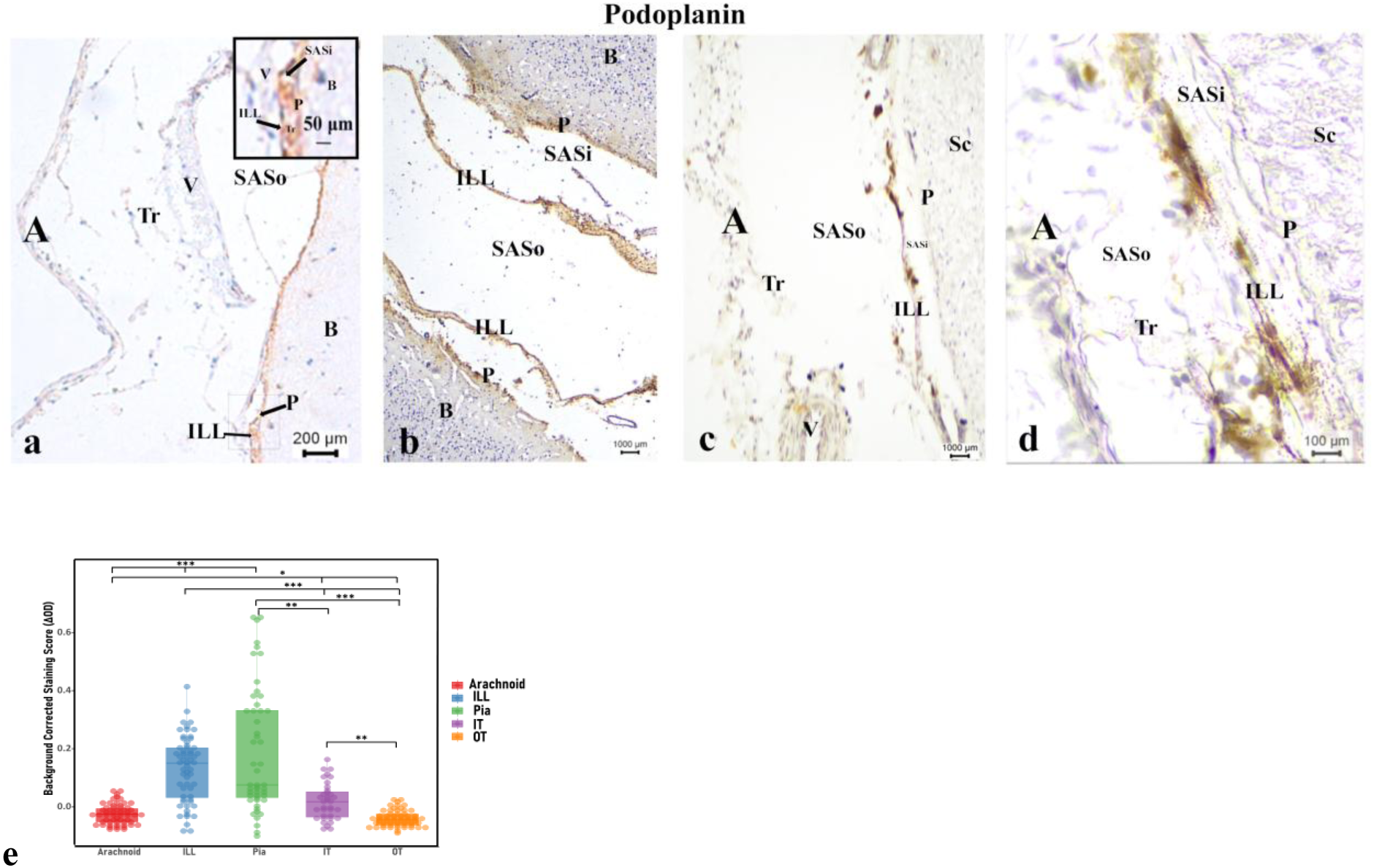
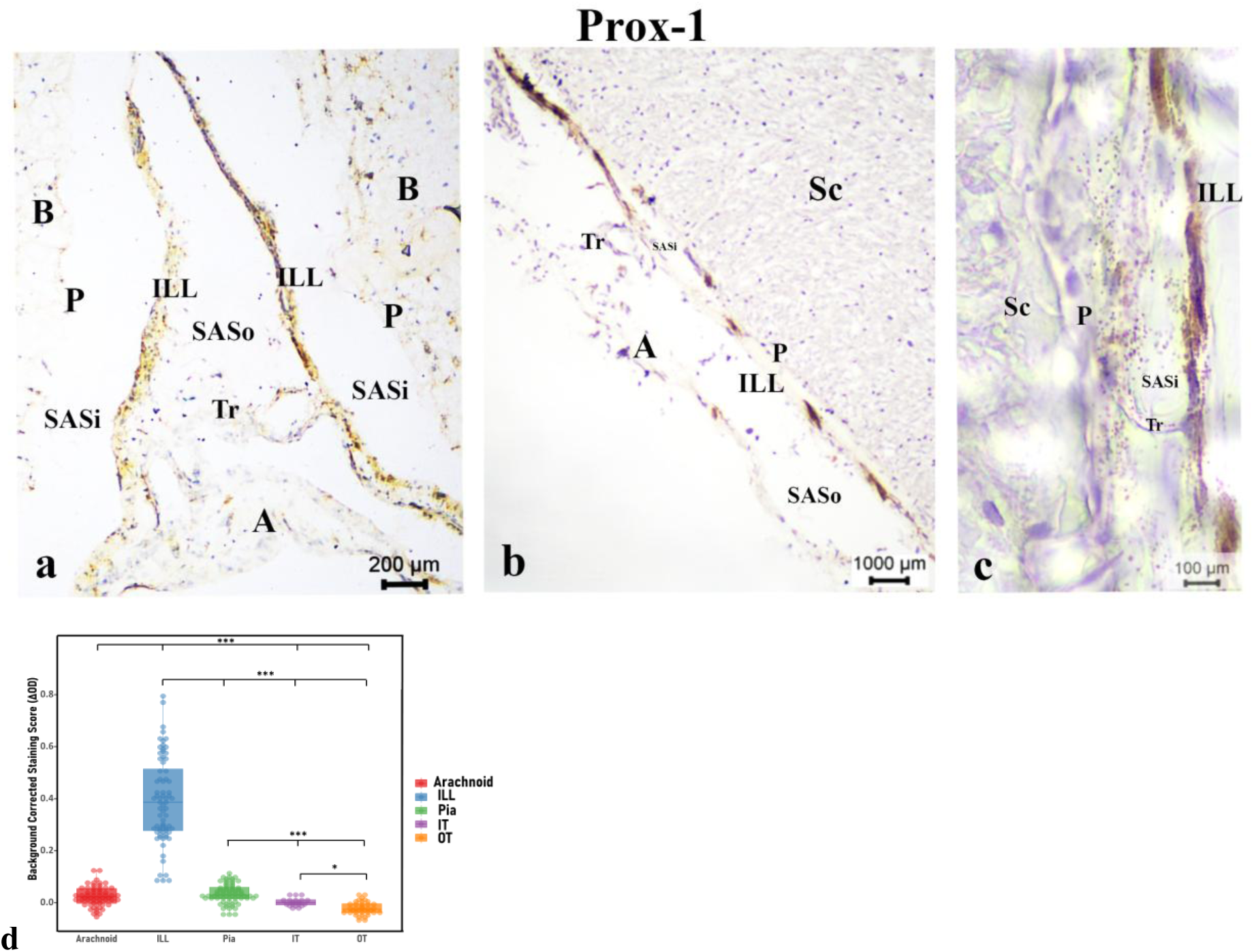
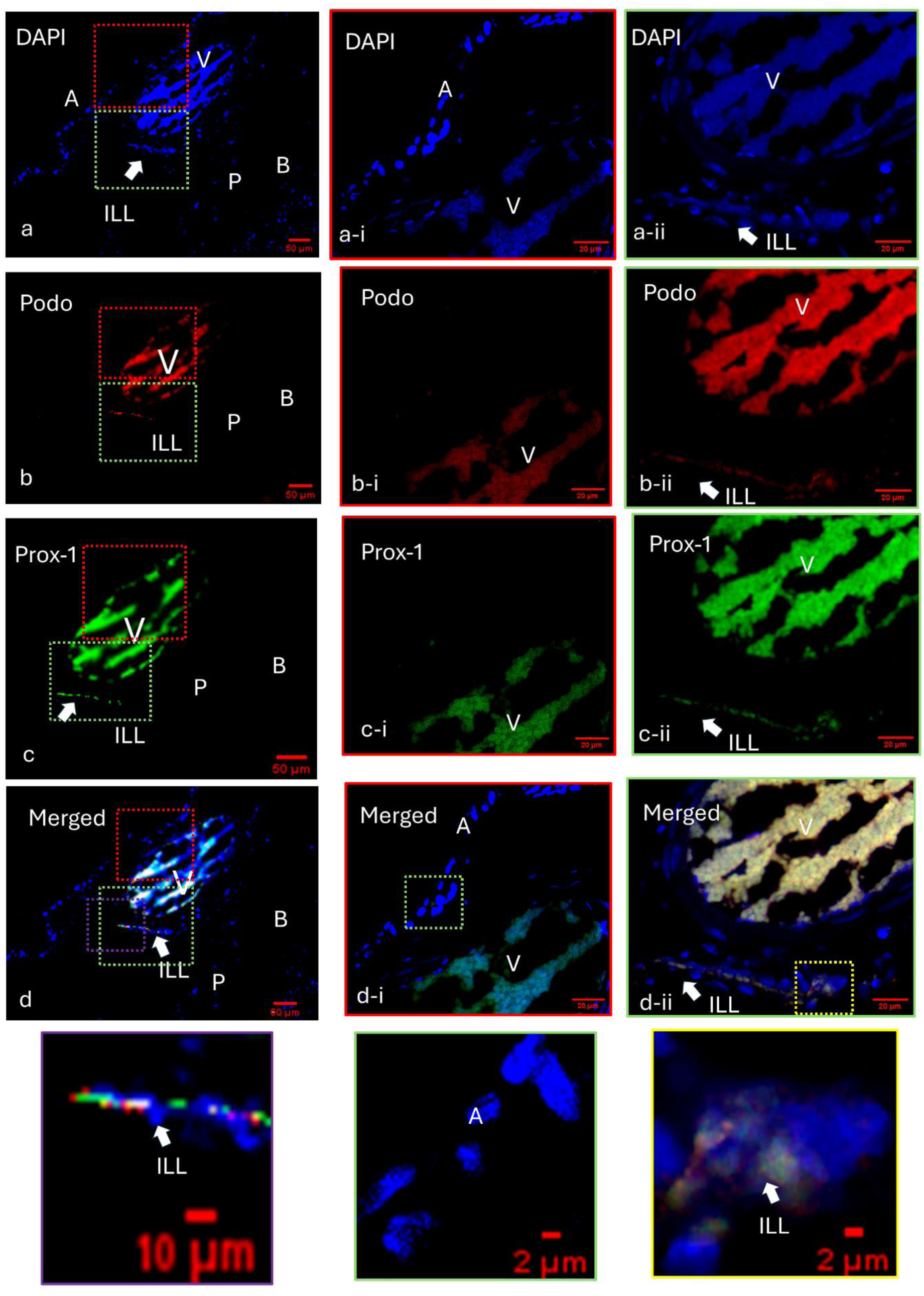
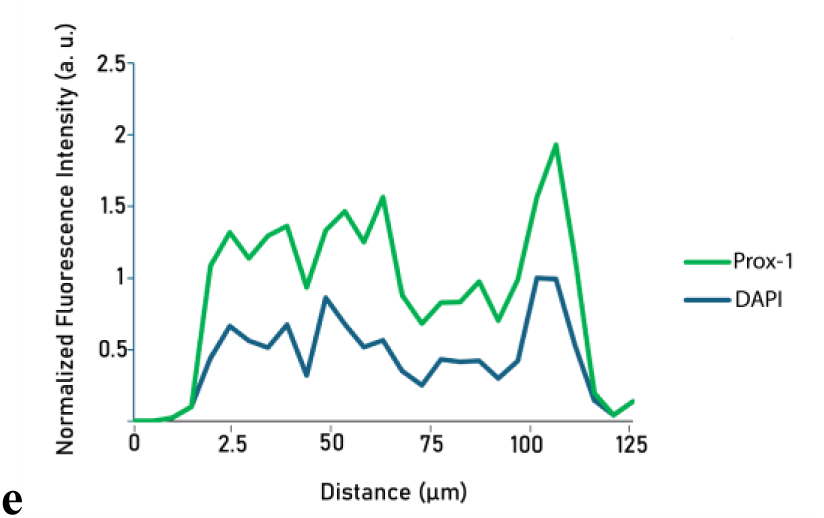
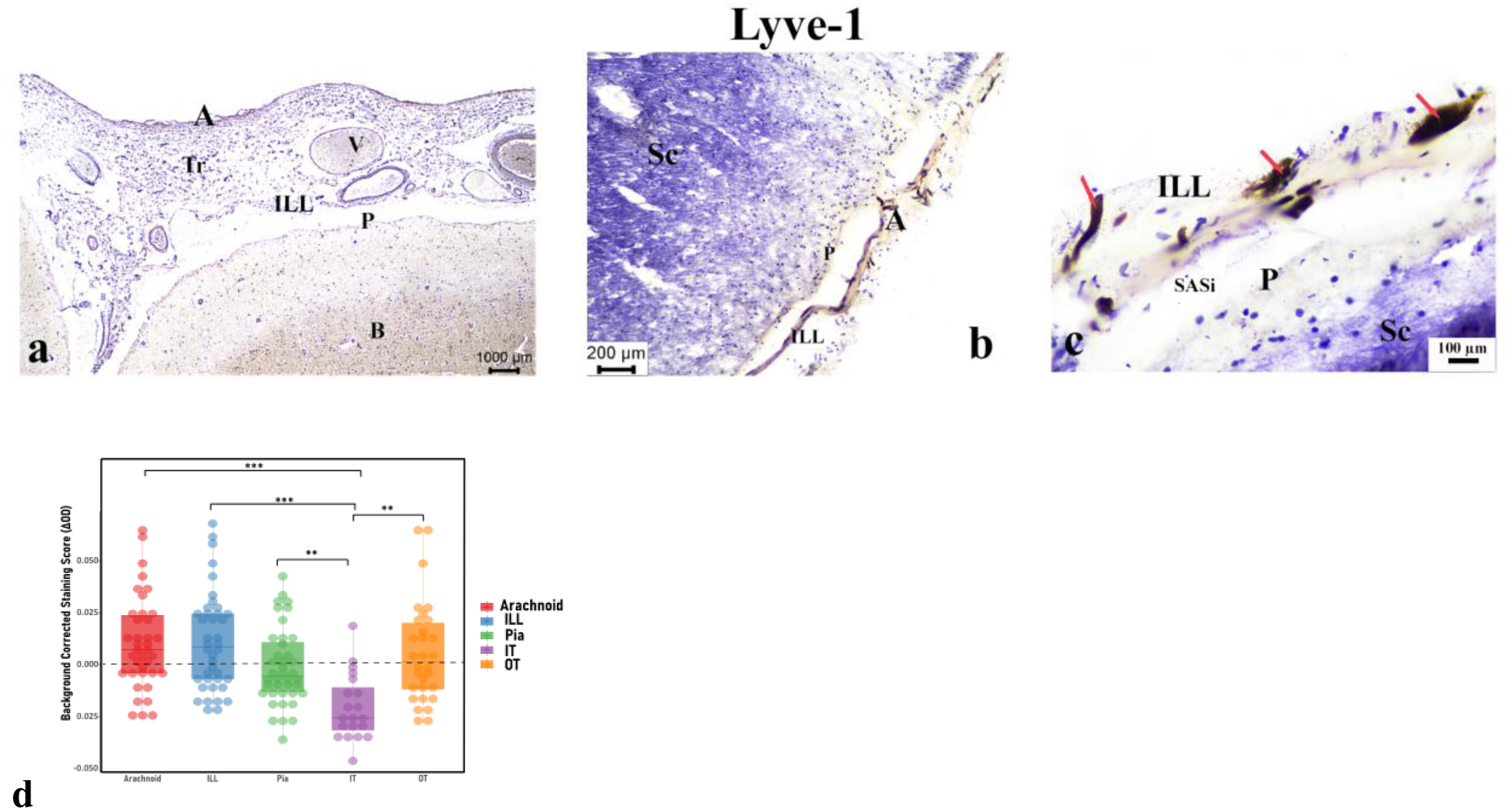
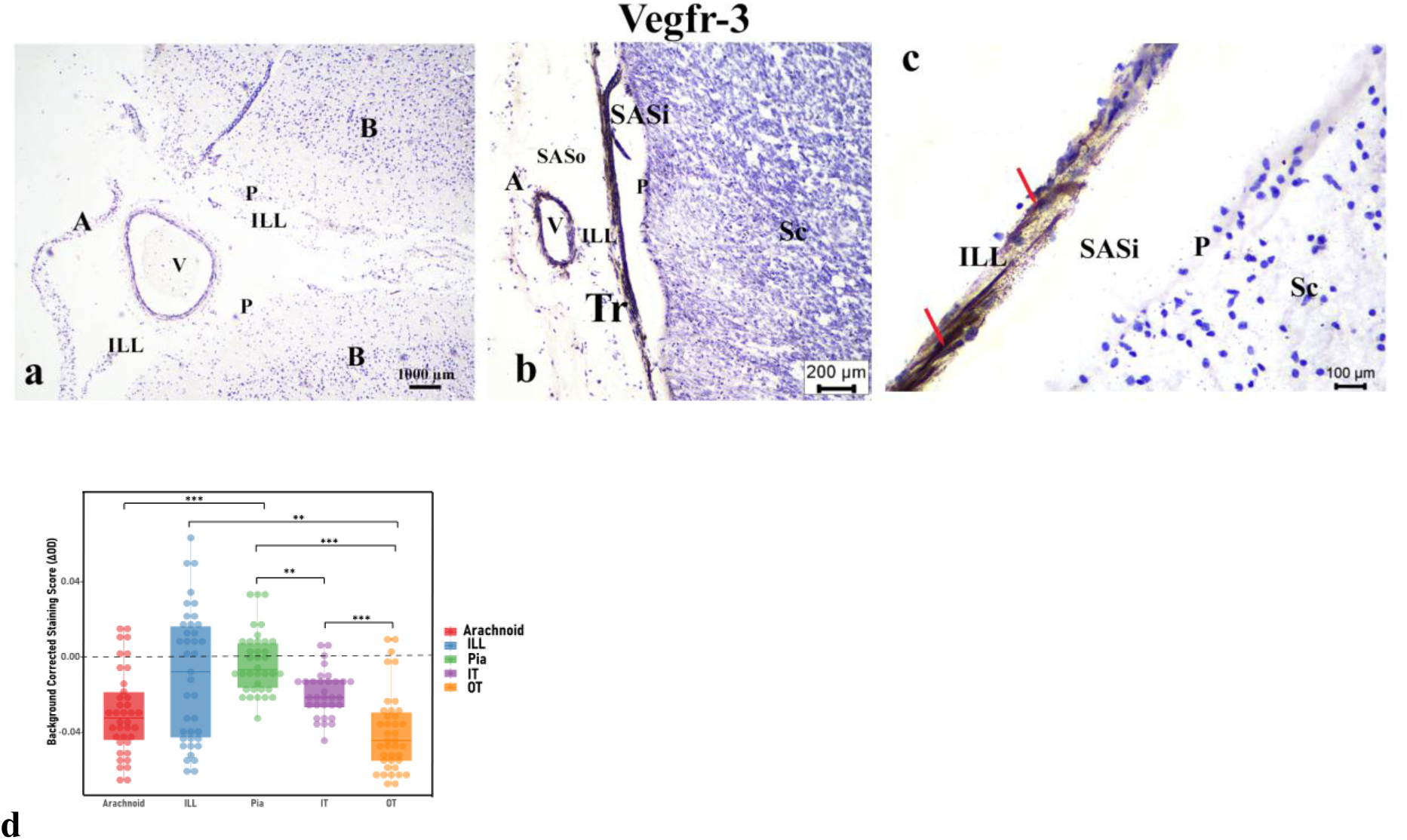
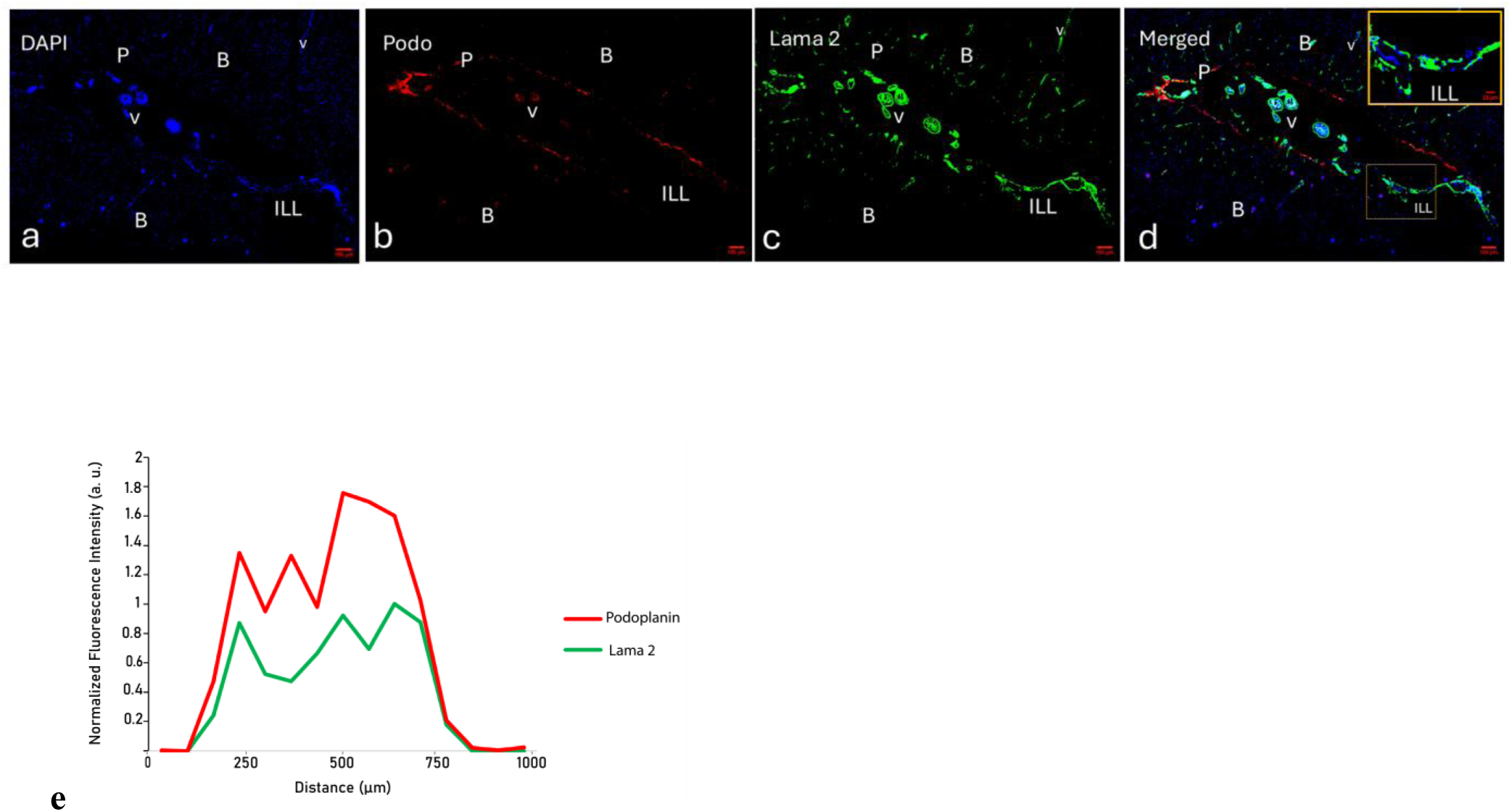
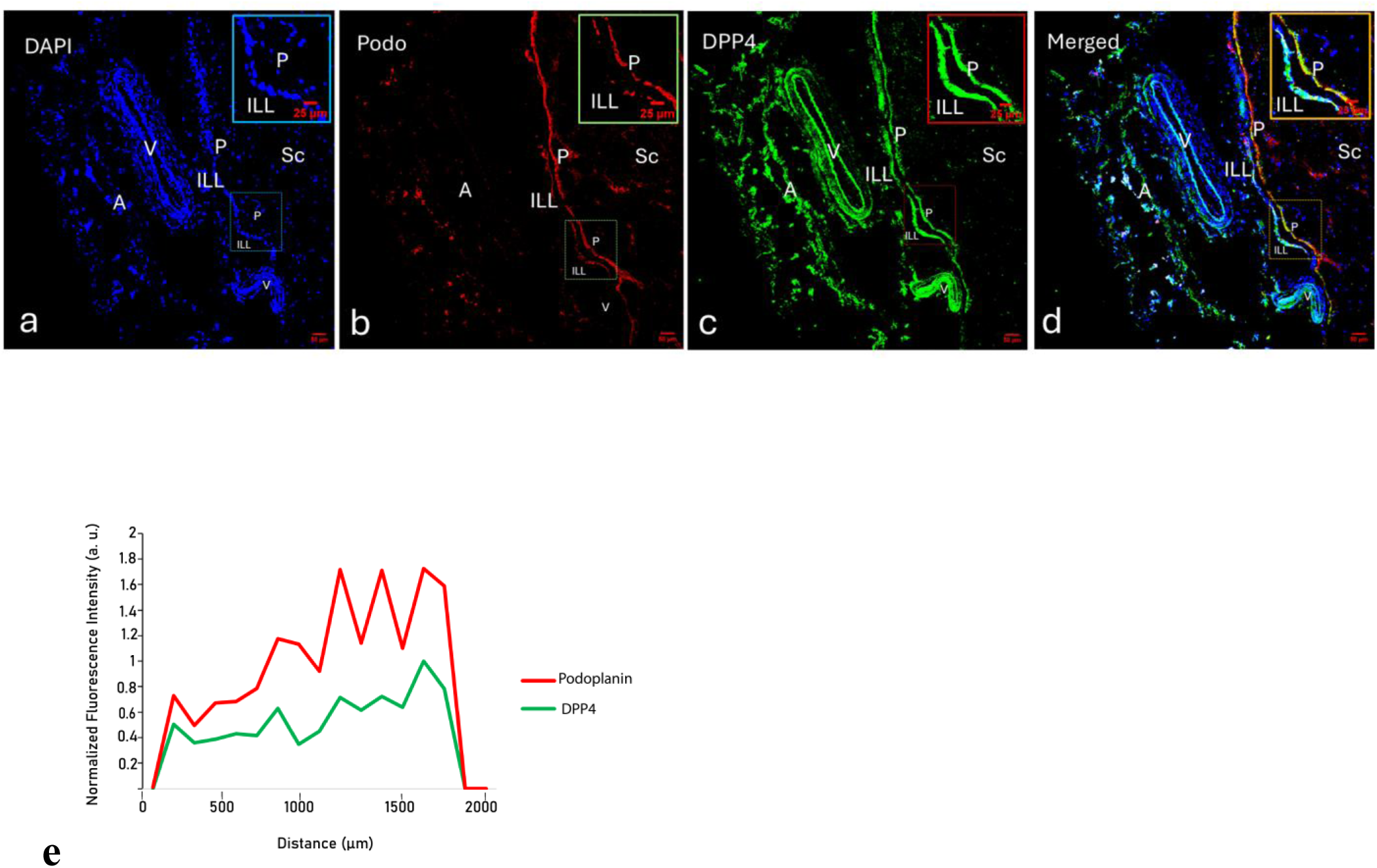
Immunohistological expression of meningeal markers in adult human brain and spinal cord. A. Podoplanin (Staining with diaminobenzidine (DAB) and counterstaining with hematoxylin) **(a-b)** On the cortical surface of the brain (B), the arachnoid mater (A) showed no Podoplanin expression, while the ILL and pia mater (P) exhibited positive staining. The ILL divided the SAS into outer (SASo) and inner (SASi) compartments. Trabeculae (Tr) in the SASo were negative for Podoplanin, whereas those in the SASi were positive. Inset showing an enlarged view of the SASi (a). **(c-d)** On the spinal surface, the arachnoid mater (A) exhibited no Podoplanin expression, whereas the ILL displayed positive staining. The pia mater (P) showed variable staining. The ILL demarcated the SAS into outer (SASo) and inner (SASi) compartments. **(e)** Raincloud plot showing regional differences in Podoplanin expression in SAS components. Robust one-way ANOVA revealed a significant regional effect (F₄,₆₇ = 32.7, p < 0.001; ES = 0.779, 95% CI 0.655–0.951). Podoplanin expression was highest in the pia and ILL, intermediate in the IT, and lowest in the arachnoid and OT. ILL and pia did not differ significantly (p = 0.655, post hoc pairwise comparisons). *p≤0.05, **p≤0.01, ***p≤0.001, n.s. p> 0.05. **B. Prox-1** (Staining by diaminobenzidine (DAB) and counterstaining by hematoxylin) **(a)** On the cortical surface of the brain (B), the arachnoid mater (A) and pia mater (P) showed no Prox-1 expression, whereas the ILL exhibited positive staining. The ILL divided the SAS into outer (SASo) and inner (SASi) compartments. A double fold of the ILL extended into the depths of the sulcus, carrying an extension of the SASo. **(b-c)** On the spinal surface, the arachnoid mater (A) and pia mater (P) showed no Prox-1 expression, whereas the ILL exhibited positive staining. The ILL divided the SAS into outer (SASo) and inner (SASi) compartments. **(d)** Raincloud plot showing regional differences in Prox-1 expression across SAS components. Robust one-way ANOVA revealed a significant regional effect (F₄,₇₁ = 83.7, p < 0.001; ES = 1.72, 95% CI 1.15–2.51). Prox-1 expression was highest in the ILL, significantly exceeding all other regions (p < 0.001, post hoc pairwise comparisons), indicating its preferential enrichment. *p≤0.05, **p≤0.01, ***p≤0.001, n.s. p> 0.05. **C. Podoplanin and Prox-1 co-expression on the cortical surface** (Stained with Alexa Fluorophores 488, 594, and DAPI) (a–d) On the cortical surface of the brain (B), the leptomeningeal arrangement was observed in relation to a subarachnoid vessel. Regions superior (a–d-i) and inferior (a–d-ii) to the vessel were further focused. a. DAPI only; b. Podoplanin; c. Prox-1; d. Merged expressions; e. Nuclear localization of Prox-1 a. The linear nuclear arrangement superior and inferior to the vessel corresponded to the arachnoid and intermediate leptomeningeal layer (ILL), respectively. b–c. The arachnoid (A), which passed over the vessel, showed no Podoplanin or Prox-1 expression. The ILL, located beneath the vessel, exhibited positive staining for both markers. Podoplanin expression was also observed in the pia mater (P). e. The line graph of normalised fluorescence intensity demonstrated prominent nuclear localisation of Prox-1 expression, supported by significant spatial overlap with DAPI (Costes P = 1). **D. Lyve-1 Expression in the Human Brain and Spinal Cord** (Staining with diaminobenzidine (DAB) and counterstaining by hematoxylin) **a. Cortex:** DAB staining for Lyve-1 on the cortical surface of the brain (B) showed no expression in any leptomeningeal layer. **b–c. Spinal cord:** DAB staining for Lyve-1 in the spinal cord, counterstained with hematoxylin, revealed no expression in the arachnoid (A) and pia mater (P) (b). However, the ILL exhibited positive staining in certain cells (b-c). The Lyve-1-positive cells displayed macrophage-like characteristics, including enlarged nuclei and cytological debris, indicating the presence of infiltrating or resident macrophages in ILL (c). **d.** Raincloud plot showing regional differences in Lyve-1 expression across SAS components. Inter-regional differences in corrected optical density values were observed. However, no significant enrichment above background (indicated by dotted black line) was observed in any region. *p≤0.05, **p≤0.01, ***p≤0.001, n.s. p> 0.05. **E. Vegfr-3 Expression in the Human Brain and Spinal Cord** **a. Cortex:** DAB staining for Vegfr-3 on the cortical surface (B) showed no expression in any leptomeningeal layer. **b-c. Spinal cord:** DAB staining for Vegfr-3 in the spinal cord, counterstained with hematoxylin, also showed no expression in the arachnoid (A) or pia mater (P) (b). However, the ILL exhibited positive staining in certain cells (b-c). The Vegfr-3-positive cells displayed macrophage-like characteristics, including enlarged nuclei and cytological debris, indicating the presence of infiltrating or resident macrophages in ILL (c). **d.** Raincloud plot showing regional differences in Vegfr-3 expression across SAS components. Inter-regional differences in corrected optical density values were observed. However, no significant enrichment above background (indicated by dotted black line) was observed in any region. *p≤0.05, **p≤0.01, ***p≤0.001, n.s. p> 0.05. **F. Podoplanin and Laminin-alpha-2 (Lama 2) co-expression in the Adult Human Brain** (Stained with Alexa Fluorophores 488 and 594, and DAPI) **a.** DAPI only; **b.** Podoplanin; **c.** Laminin-alpha-2 (Lama 2); **d.** Merged expressions; **e.** Line graph demonstrating co-expressions of Podoplanin and Lama 2 in ILL. **(a–e)** The leptomeningeal arrangement within the cortical sulcus of the brain (B) is shown. a. The focus was on a midline structure traversing the sulcus. The linear nuclear arrangement corresponded to a double fold of the leptomeningeal layer, separated from the cortical margins and enclosing blood vessels (V) between its layers. b–d. Both the ILL and the pia mater (P) stained positively for Podoplanin and Lama 2, suggesting mesodermal origin and the presence of a basement membrane, respectively. An inset from panel d is enlarged to highlight the co-expression of Podoplanin and Lama 2 within the ILL. e. The line graph of normalized fluorescence intensity profiles for Podoplanin and Lama 2, demonstrating their co-expression in ILL. **G. Podoplanin and Dipeptidyl peptidase 4 (DPP4) co-expression in the Human Spinal Cord** (Stained with Alexa Fluorophores 488 and 594, and DAPI) **a.** DAPI only; **b.** Podoplanin; **c.** Dipeptidyl peptidase 4 (DPP4); **d.** Merged expressions; **e.** Line graph demonstrating normalized fluorescence intensity profiles for Podoplanin and DPP4 within the ILL. **(a–e)** The leptomeningeal arrangement at the surface of the spinal cord (Sc) is shown. Insets from panels a–d are enlarged to highlight the distinct expression patterns of the markers. A line graph of normalized fluorescence intensity profiles for Podoplanin and DPP4 within the ILL is presented to assess their co-expression (e). **a.** The focus was on the surface of the spinal cord (Sc). The nuclear arrangements delineated the three leptomeningeal layers from outer to inner: arachnoid mater (A), ILL, and pia mater (P). Multiple blood vessel profiles (V) are visible between these meningeal layers. **b–d.** The ILL and pia mater (P) exhibited positive staining for Podoplanin; however, the arachnoid mater and perivascular sheath were negative. DPP4 expression was detected in all leptomeningeal layers, as well as in the trabeculae and blood vessels, including the perivascular sheath. A magnified view of the inset from b–d is presented in the upper right corner of each image. **e.** A line graph of normalized fluorescence intensity profiles for Podoplanin and DPP4 within the ILL is demonstrating their co-expression. Abbreviations: A-arachnoid, P-pia, ILL-intermediate leptomeningeal layer, Tr-trabeculae, N-nerve, V-vessel, SAS-subarachnoid space, i-internal, o-outer. B-brain, Sc-spinal cord.

**Figure 4.**
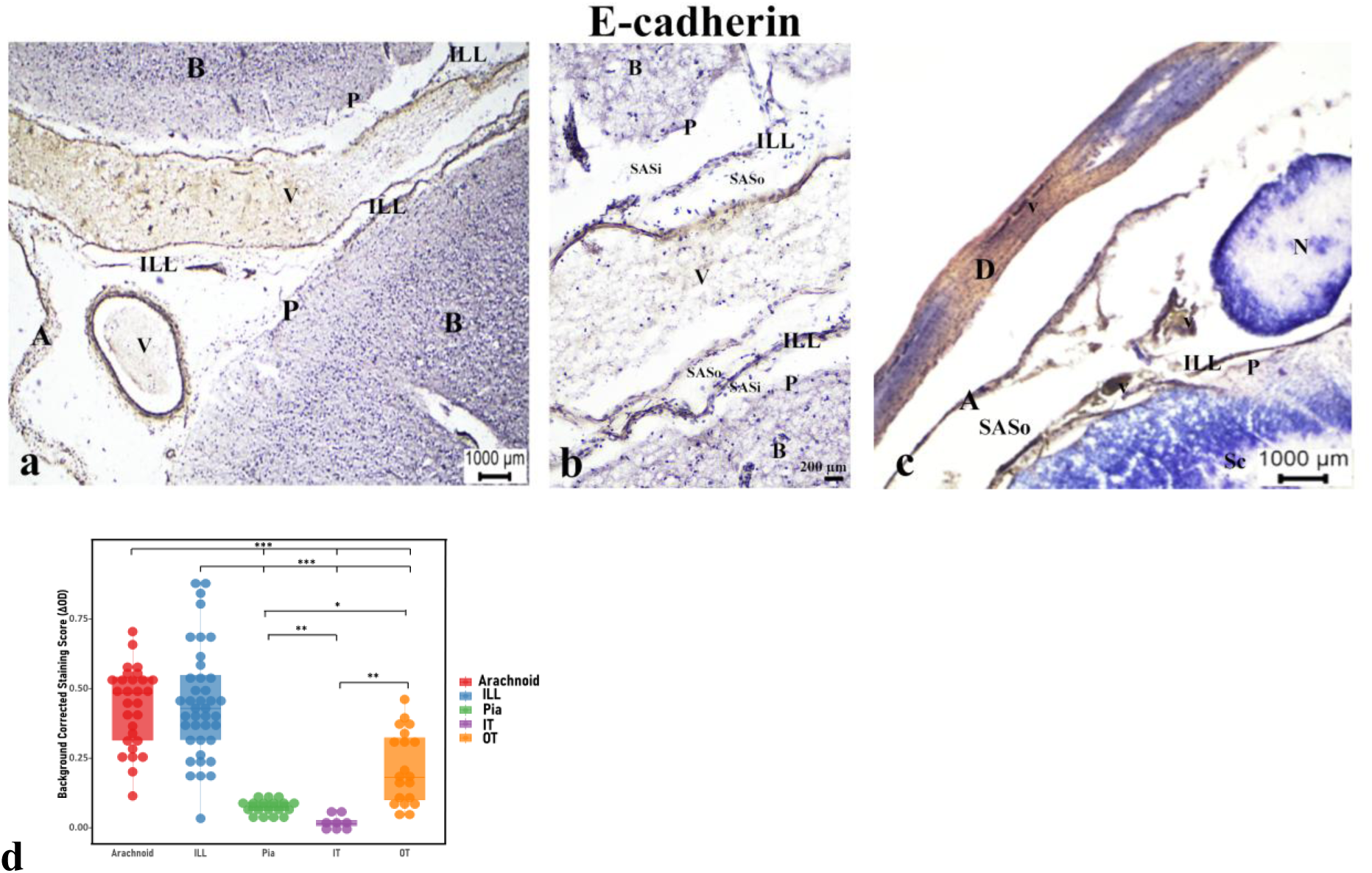
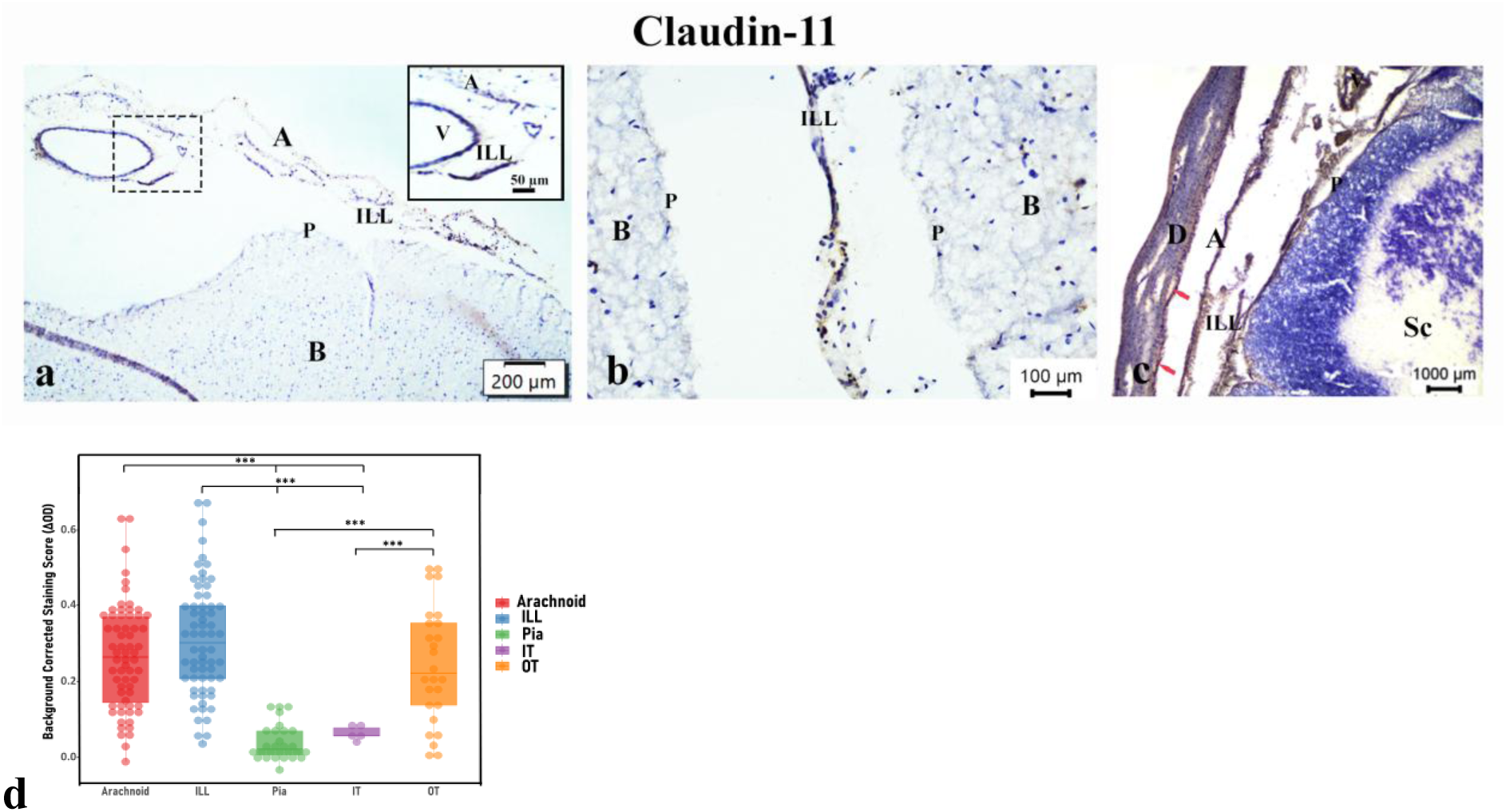
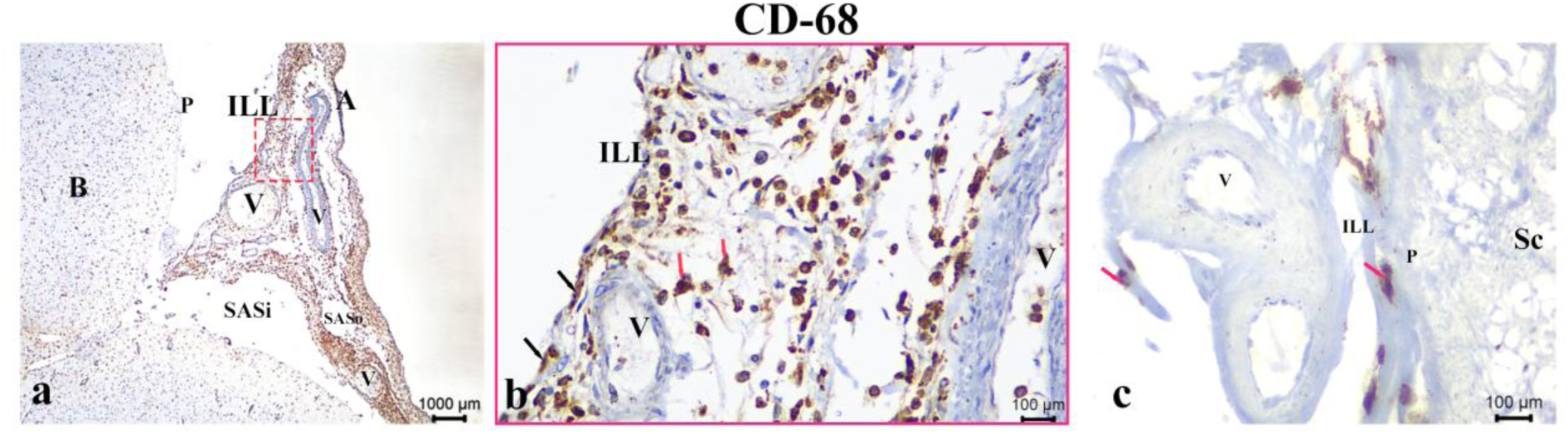
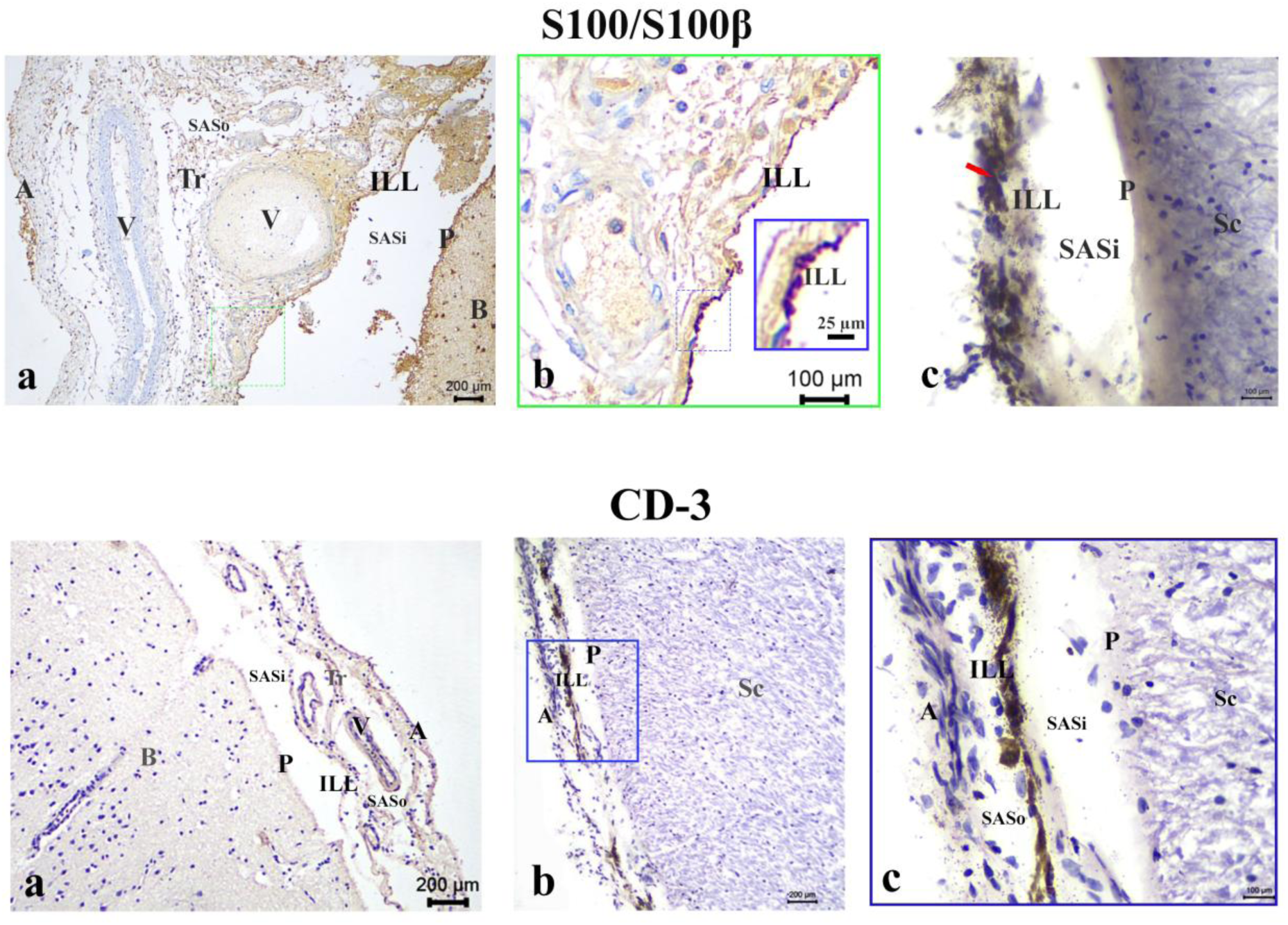
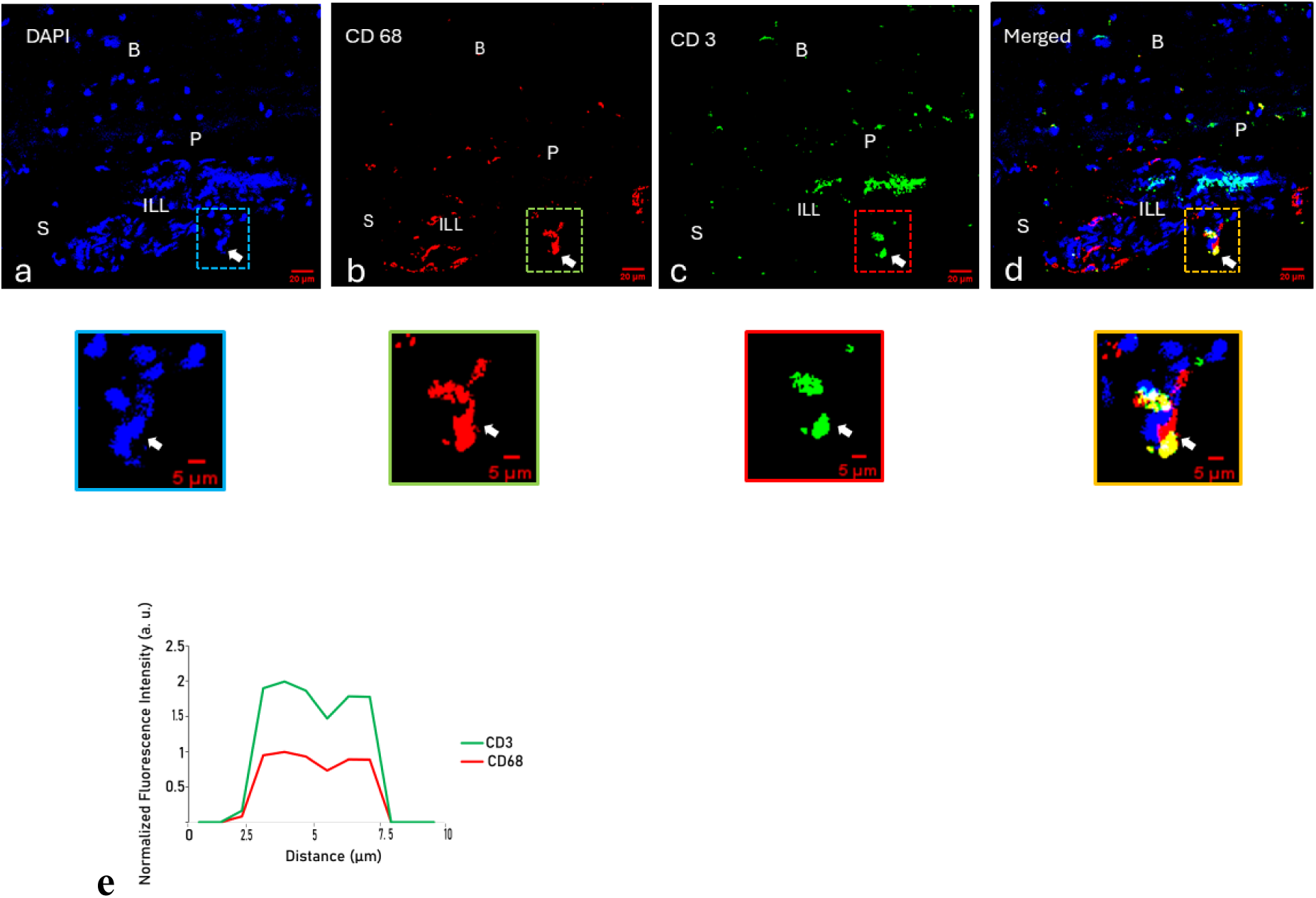
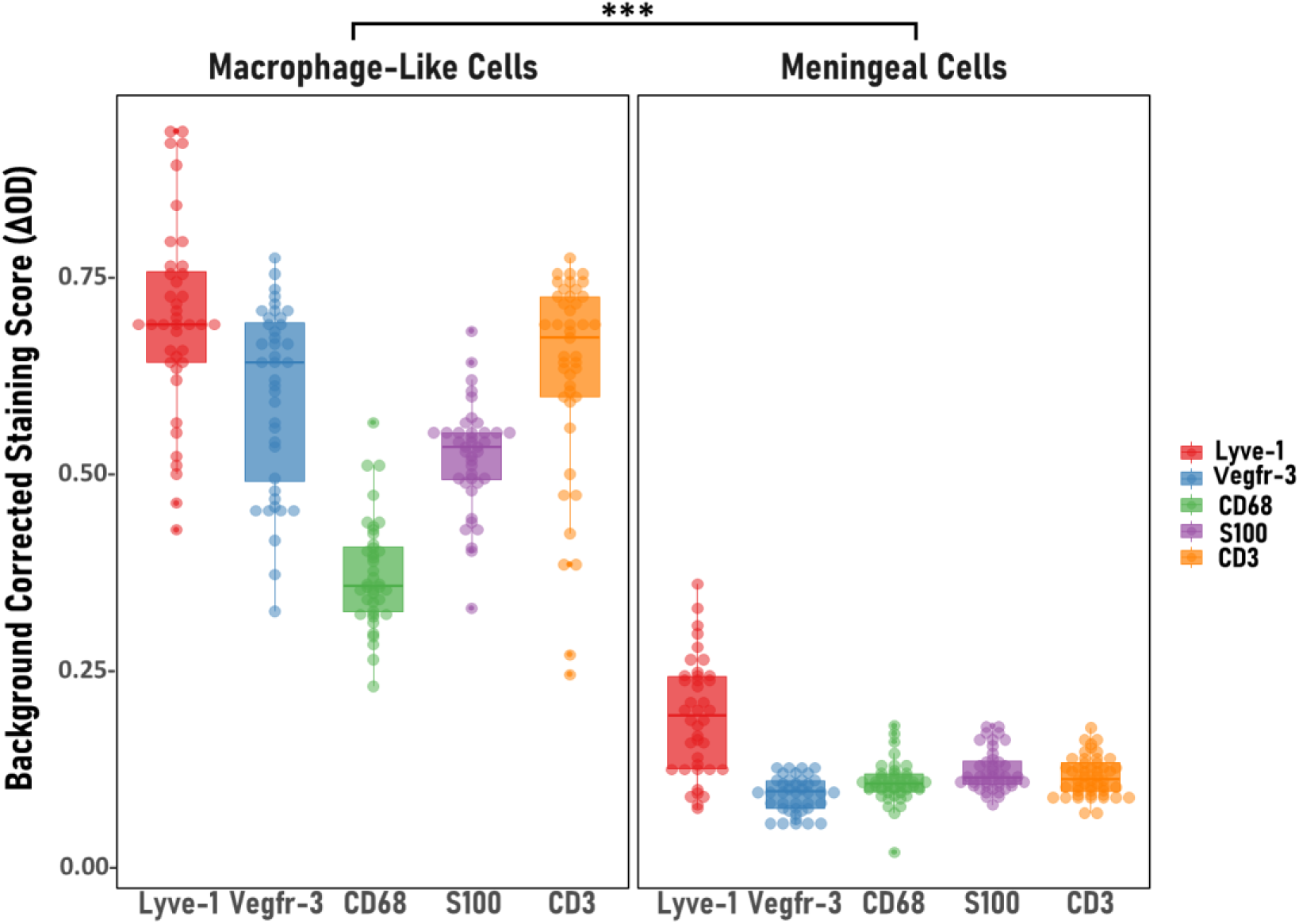
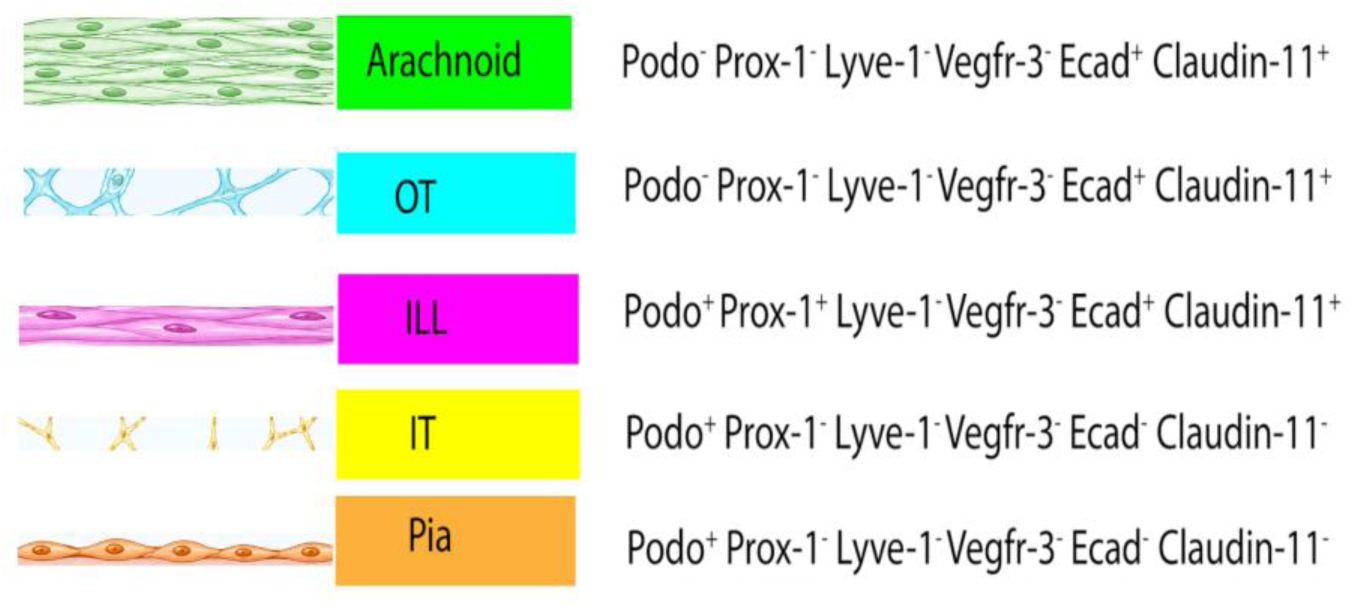
Immunohistological expression of meningeal markers in adult human brain and spinal cord. **A. E-cadherin** (Staining with diaminobenzidine (DAB) and counterstaining with hematoxylin) **(a–b) Brain**: On the cortical surface and within a sulcus of the brain (B), E-cadherin expression, an adherens junction marker, was observed in the leptomeningeal layers. E-cadherin expression was detected in both the arachnoid (A) and the ILL. A double fold of the ILL extended into the sulcus, enclosing vascular structures (V). An inset (b) shows distinct E-cadherin expression in the ILL. E-cadherin expression was also observed in the vascular endothelium (a–b). **(c) Spinal Cord**: The meningeal arrangement, including the dura mater, was visualized in the adult spinal cord (Sc). E-cadherin expression was evident in the arachnoid (A) and the ILL. The ILL divided the SAS into outer (SASo) and inner (SASi) compartments. Multiple vascular profiles and cross-sections of spinal nerve rootlets were present between the arachnoid and ILL. **(d)** Raincloud plot showing regional differences in E-cadherin expression across SAS components. Robust one-way ANOVA revealed a significant regional effect (F₄,₂₆.₆ = 79.0, p < 0.001; ES = 0.800, 95% CI 0.725–0.938). E-cadherin was enriched in the arachnoid and ILL, with additional expression in the OT. Arachnoid and ILL did not differ significantly (p = 0.543, post hoc pairwise comparison). *p≤0.05, **p≤0.01, ***p≤0.001, n.s. p> 0.05. **B. Claudin-11** (Staining with diaminobenzidine (DAB) and counterstaining by hematoxylin) **(a–b) Brain**: On the cortical surface and within a sulcus of the brain (B), Claudin-11 expression, a tight junction marker, was observed in the leptomeningeal layers. Claudin-11 expression was detected in both the arachnoid (A) and the ILL, as well as in the vascular endothelium. An enlarged view of the inset distinctly shows Claudin-11 expression in the ILL. The pia mater (P) stained negatively for Claudin-11. A double fold of the ILL that extended into the sulcus showed a distinct Claudin-11 expression. **(c) Spinal Cord**: The meningeal arrangement, including the dura mater, was visualized in the adult spinal cord (Sc). Claudin-11 expression was evident in the arachnoid (A) and the ILL. In the dura mater, Claudin-11 expression was noted in the neurothelial border (indicated by red arrow). The pia mater (P) stained negatively for Claudin-11. **(d)** Raincloud plot showing regional differences in Claudin-11 expression across SAS components. Robust one-way ANOVA revealed a significant regional effect (F₄,₂₉.₈ = 55.6, p < 0.001; ES = 0.723, 95% bootstrap CI 0.586–0.921). Claudin-11 was enriched in the arachnoid and ILL, with additional expression in the OT. Arachnoid and ILL did not differ significantly (p = 0.350, post hoc pairwise comparison). *p≤0.05, **p≤0.01, ***p≤0.001, n.s. p> 0.05. **C. CD68 Expression in the Adult Human Brain** (Staining with diaminobenzidine (DAB) and counterstaining with hematoxylin) **(a-b) Injured brain**: a. In a case of traumatic brain injury where death occurred a few days after the incident, subarachnoid hemorrhage and dilated blood vessels were noted. The extravasated blood and infiltrating immune cells (mononuclear phagocytes) were confined between the arachnoid mater (A) and the ILL, i.e., in the SASo. Extensive CD68 expression was evident in this region. In addition to the CSF spaces, CD68-positive cells also appeared to be present in ILL. b. Magnified inset images from (a), focused toward the ILL, showed intense CD68 expression within the ILL, indicating the presence of resident or infiltrating macrophages (blue arrows). The CD68 expression was also observed in the trabeculae and perivascular sheath. **(c) Non-injured spinal cord**: High-resolution views of the ILL show that the CD68 expression is intermittent, with unstained meningeal cells interspersed. CD68 expression was also detected in the perivascular sheath surrounding vascular profiles between the arachnoid and ILL (blue arrows). **D. S100/S100β Expression in the Adult Human Brain and Spinal Cord** (Staining with diaminobenzidine (DAB) and counterstaining with hematoxylin) **a–b. Injured brain**: a. In a case of traumatic brain injury where death occurred a few days after the incident, subarachnoid hemorrhage and dilated blood vessels were noted. S100 expression was observed in each leptomeningeal layer, from outer to inner: arachnoid (A), ILL, and pia (P). The extravasated blood and infiltrating immune cells (mononuclear phagocytes) were confined between the arachnoid mater and the ILL, i.e., within the outer subarachnoid space (SASo). b. High-resolution views of the outer SAS (SASo) revealed intense S100 expression in the meningeal cells within the ILL (also shown in a further magnified inset view). **c. Non-injured spinal cord**: S100β was expressed in the ILL. It was intermittent, with unstained meningeal cells interspersed. **E. CD3 expression in the Human Brain and Spinal Cord** (Staining with diaminobenzidine (DAB) and counterstaining with hematoxylin) **(a) Non-injured brain**: On the cortical surface of the non-injured brain (B), CD3 expression—a common T cell marker—was not detected in any component of the SAS. **(b–c) Non-injured spinal cord**: Views were taken at the spinal surface. **b.** Expression of CD3 was observed in the ILL. **c.** High-resolution views of the ILL showed that CD3 expression is intermittent, with unstained meningeal cells interspersed. **F. CD68 and CD3 Co-expression in the Adult Human Brain** (Stained with Alexa Fluorophores 488 and 594, and DAPI) (a–d) Within the cortical sulcus of the brain (B), the leptomeningeal arrangement was observed. Insets from panels a–d are enlarged to highlight the distinct expression patterns of the markers. (e) The line graph of normalised fluorescence intensity data for CD3 and CD68 demonstrated significant non-random spatial association (Pearson’s R = 0.40; thresholded R = –0.23; tM1 ≈ 0.48, tM2 ≈ 0.58; ICQ = 0.40; Costes P = 1.00). a. DAPI only; b. CD68; c. CD3; d. Merged expressions; e. Line graph demonstrating CD 68 and CD3 colocalization a. The focus was on a midline structure traversing the sulcus. The linear nuclear arrangement corresponded to a double fold of the leptomeningeal layer, separated from the cortical margins. b–d. CD68 and CD3-positive cells were observed in ILL. Both markers showed cytoplasmic expressions and embraced a large, lobulated nucleus, indicating macrophage-like cells. A subset of CD68-positive cells co-expressed CD3, indicating a unique population of border-associated macrophages (BAMs). A magnified view of the insets from b–d is presented below each image. **G. A comparative immunophenotypic analysis of macrophage-like and meningeal cells in ILL in the Adult Human Spinal Cord**. Raincloud plot showing differential expression of the immune markers: Lyve-1, Vegfr-3, CD68, S100, and CD3 between macrophage-like cells and meningeal cells within the ILL in the adult human spinal cord. Across all markers, macrophage-like cells demonstrated significantly higher expression levels than meningeal cells (all p < 0.001, post hoc pairwise comparisons). *p≤0.05, **p≤0.01, ***p≤0.001, n.s. p> 0.05 **H. Immunophenotypic landscape of SAS components in Adult Human**

ILL expressed Lama 2, which co-expressed with Podoplanin, indicating the presence of a basement membrane for this layer (Fig. 3-F). DPP4 (CD26), previously considered arachnoid-specific, was also expressed in the ILL and pia, indicating limited layer specificity (Fig. 3-G, S2-o).

### Perivascular and perineural organization of the ILL

The ILL formed a perivascular sheath around arterial profiles, localized predominantly to the inner compartment in the cortex and the outer compartment in the spinal cord (Fig. S2: a-b). ILL contributing to the vascular sheath formation was further confirmed in SEM and TEM (see Results: Ultrastructural observations, Fig. 5, 6). It also formed perineural sheaths around spinal nerve rootlets (Fig. S2: d).

**Figure 5.**
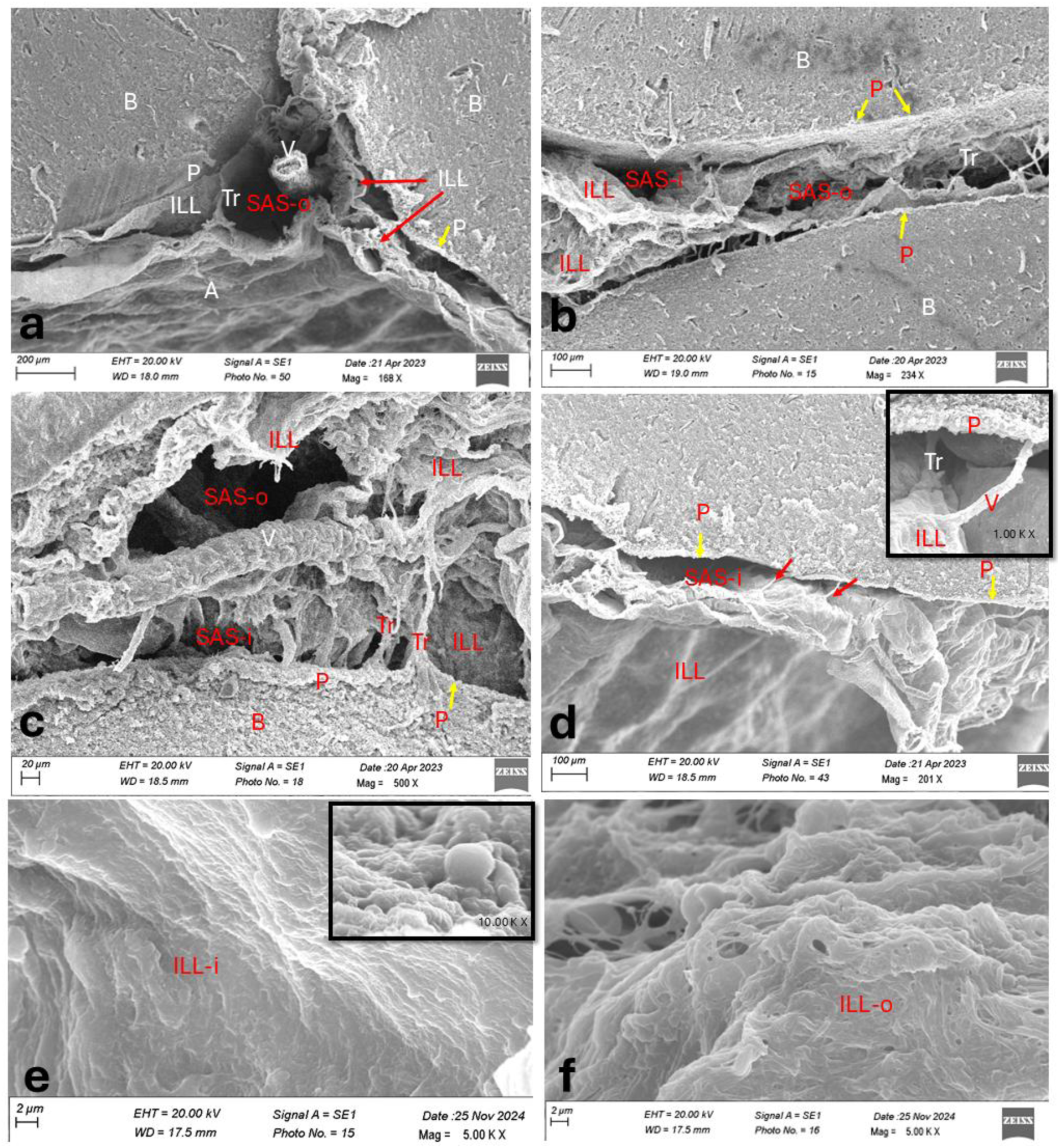
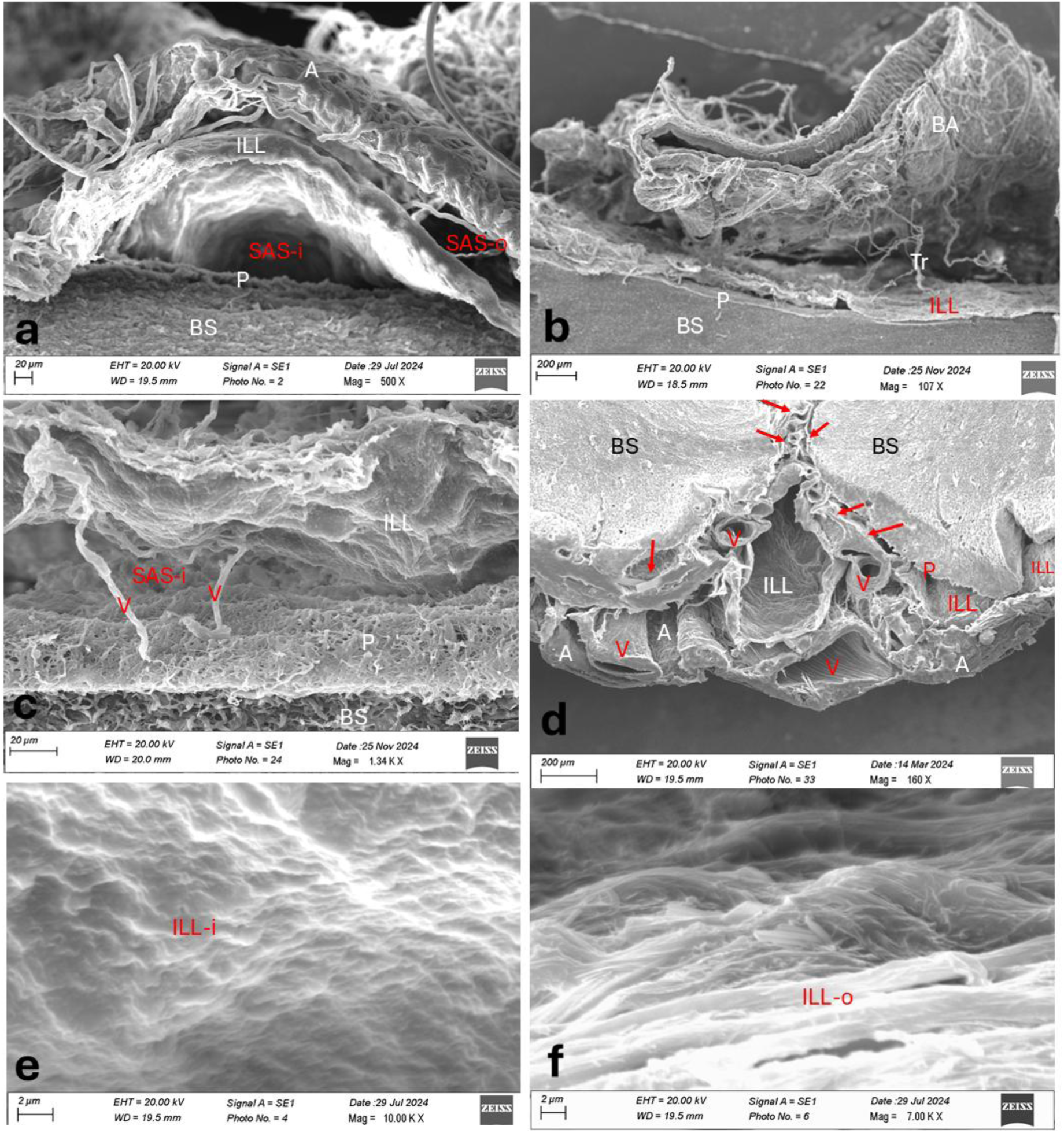
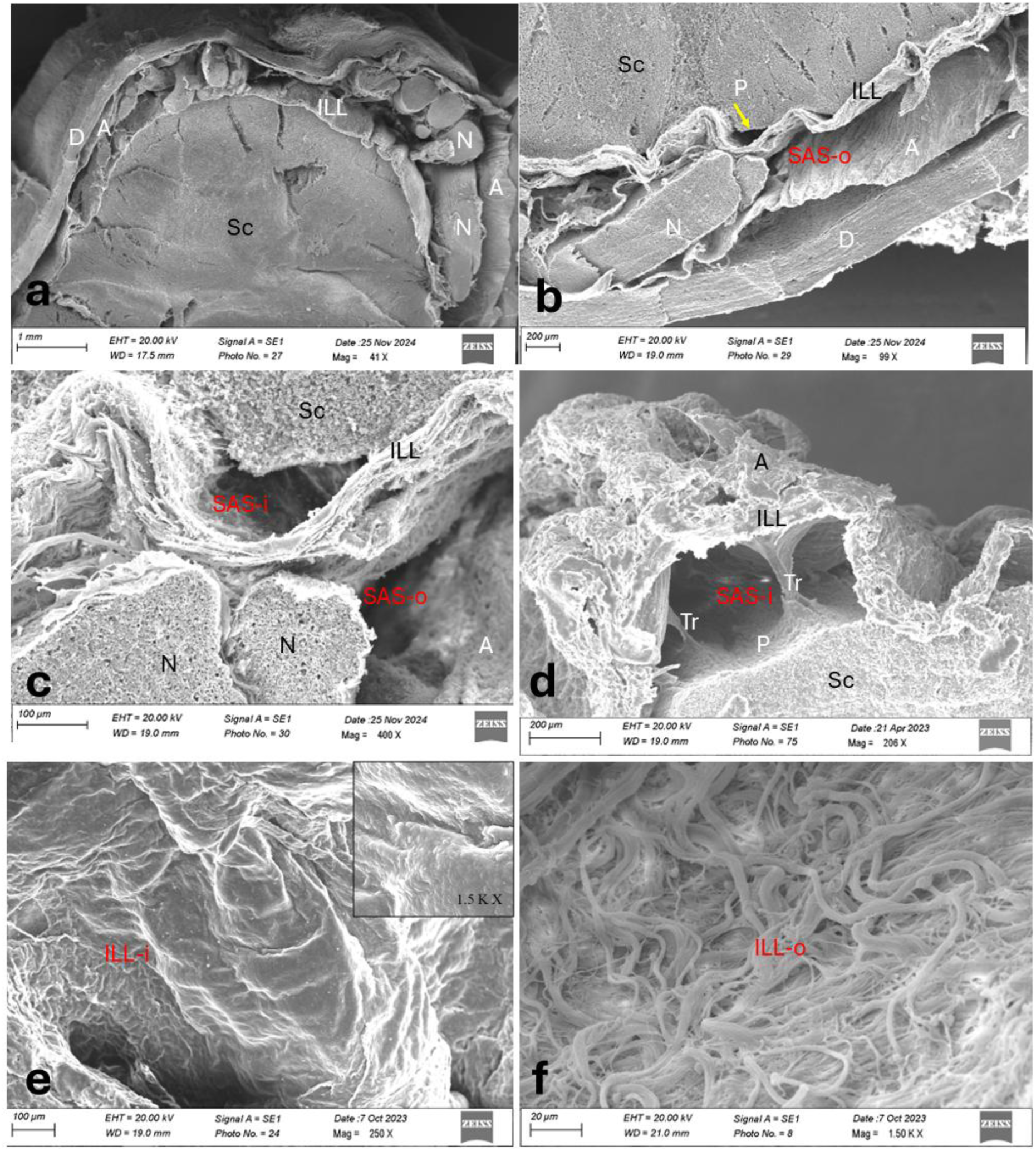
Scanning Electron Microscopy of meningeal arrangement around Adult Human Brain and Spinal Cord. **A. Cortex** **(a–f):** Scanning electron microscopy (SEM) of the postmortem adult human brain (B) specimens was performed. The dura mater was removed during autopsy as per standard protocol. The cortical surface and sulci were examined to observe the arrangement of the leptomeningeal layers. **(a)** At the cortical surface, three layers can be identified from outer to inner: Arachnoid (A), ILL, and Pia (P). The pia is the thinnest layer and is adherent to the cortical surface (yellow arrow). In contrast, the arachnoid and ILL are comparatively thicker and resemble each other. The ILL divides the SAS into the outer SAS (SASo) and inner SAS (SASi). Blood vessels (V) and trabeculae (Tr) are observed between the leptomeningeal layers, being more prominent in the SASo. **(b–c)** A double fold of the ILL enters the sulcus, enclosing the blood vessels (V). Within the sulcus, only the pia (P, marked by the yellow arrow) and ILL are visible, as the arachnoid (A) bridges over the sulcus and does not enter it. The CSF space that enters between the double folds of the ILL is a continuation of the SASo. Interlayer trabeculae (Tr) connecting the ILL folds are also observable. **(d)** Fine blood vessels and trabeculae (Tr) can be seen between the ILL and pia. In the SASi, the blood vessels carry a sheath derived from the ILL. The inset shows a magnified view of a trabecula (Tr) and a blood vessel (V, red arrows) connecting the ILL and pia (yellow arrows). **(e)** The surface of the ILL facing the pia is very smooth and non-sieved. The inset presents a highly magnified view of this ILL surface, confirming its non-sieved nature. Blebs or podia are visible, indicating the epithelial character of the ILL. **(f)** The surface of the ILL facing the arachnoid is rough and fibrous. Interlacing fiber bundles and perforations are visible. **B. Brainstem** **(a–f):** Scanning electron microscopy (SEM) of the postmortem adult human brainstem (BS) specimens. The dura mater was removed during autopsy as per standard protocol. The neural surface and sulci were examined to observe the arrangement of the leptomeningeal layers. **(a)** At the meningeal surface, three layers can be identified from outer to inner: Arachnoid (A), ILL, and Pia (P). The pia is the thinnest layer and is adherent to the neural surface. In contrast, the arachnoid and ILL are comparatively thicker. The ILL divides the SAS into the outer SAS (SASo) and inner SAS (SASi). **(b–c)** The shallow basilar sulcus over the pons was examined. The basilar artery was located in the SASo between the arachnoid (not shown here) and the ILL. Fine blood vessels (V) are seen running between the ILL and the pia (P). **(d)** The leptomeningeal arrangement was examined over the anterior median sulcus at the lower end of the medulla oblongata. A double fold of the ILL enclosing blood vessels (V) is seen entering the sulcus (marked by red arrows). Vascular profiles are present between the arachnoid (A) and ILL, as well as between the ILL and pia (P), helping distinguish them as independent layers. The vascular profiles between the arachnoid and ILL are comparatively larger. Where blood vessels are absent, the ILL may adhere to either the arachnoid or pia, making its identification more difficult. **(e)** The surface of the ILL facing the pia is very smooth and non-sieved. **(f)** The surface of the ILL facing the arachnoid is rough and fibrous. Interlacing fiber bundles are visible. **C Spinal Cord** **(a–f)** Scanning electron microscopy (SEM) of the postmortem adult human spinal cord (Sc) specimens was performed. The dura mater was left intact. The spinal surface was examined to observe the arrangement of the meningeal layers. **(a-c)** At the meningeal surface, four layers can be identified from outer to inner: Dura (D), Arachnoid (A), ILL, and Pia (P). The dura is the thickest, and its neurothelial surface towards the arachnoid is seen intact. Among the leptomeningeal layers, the pia is the thinnest layer and is adherent to the neural surface. In contrast, the arachnoid and ILL are comparatively thicker and resemble each other. The ILL divides the SAS into the outer SAS (SASo) and inner SAS (SASi). The cross-sections of spinal nerve rootlets (N) are also seen between the arachnoid and ILL. A higher resolution view shows the spinal ILL is fibrocellular and multilayered (c). **(d)** A high-resolution view of SASi reveals the trabeculae (Tr) extending between ILL and pia. The meningeal ends of the trabeculae widen to give them the appearance of an inverted pyramid. The SASo is seen obliterated, and ILL adhered to the arachnoid. **(e)** The surface of the ILL facing the pia is very smooth and non-sieved. The inset presents a highly magnified view of this ILL surface, confirming its non-sieved nature. Blebs or podia are visible, indicating the epithelial character of the ILL. **(f)** The surface of the ILL facing the arachnoid is rough and fibrous. Interlacing fiber bundles and perforations are visible. Abbreviations: D-dura, A-arachnoid, P-pia, ILL-intermediate leptomeningeal layer, Tr-trabeculae, N-nerve, V-vessel, BA-basilar artery, SAS-subarachnoid space, i-inner, o-outer. B-brain, Sc-spinal cord.

### Barrier properties of the ILL

The ILL expressed E-cadherin and Claudin-11 (Fig. 4A–B, S2f–i), with levels comparable to the arachnoid (p = 1), but displayed a more discontinuous pattern (Fig. 4A–B, S2f–i). The E-cadherin and Claudin-11 were also expressed in the outer but not in the inner trabeculae. Histological and immunostaining analyses of traumatic brain injury specimens (CD68, S100/S100β) demonstrated that the ILL restricts the spread of extravasated blood and infiltrating immune cells (Fig. 4C–D; see Results: Morphometric and quantitative estimations). Ultrastructural analyses further supported its barrier role, showing intact, non-perforated surfaces and junctional complexes on SEM and TEM (see Results: Ultrastructural observations, Fig. 5, 6).

### Macrophage-like cells in the ILL

Macrophage-like cells were identified within the ILL, predominantly in the spinal cord. These cells were larger, with irregular nuclei and cytoplasm rich in granular phagocytic debris on routine and specialized staining (Figs. 3–4, S2). They expressed Podoplanin and Prox-1 (Figs. 3A–B, S2a–e), along with Lyve-1 and Vegfr-3 (Fig. 3D-E), and showed CD68 and S100/S100β positivity, confirming their macrophage-like phenotype (Fig. 4C-D, S2j-n). Unlike circulating macrophages, they were localized exclusively to the ILL (Fig. 4C-c, D:a-c).

A subset of these cells also expressed CD3 (Fig. 4E: a–c, F). Colocalization analysis of CD68 and CD3 expressions demonstrated significant non-random spatial association between markers (Fig. 4G, Pearson’s R = 0.40; thresholded R = –0.23; tM1 ≈ 0.48, tM2 ≈ 0.58; ICQ = 0.40; Costes P = 1.00), consistent with a specialized population of border-associated macrophages (BAMs). Their presence was further supported by TEM imaging (see Results: Ultrastructural observations, Fig. 6 m–n).

**Figure 6.**
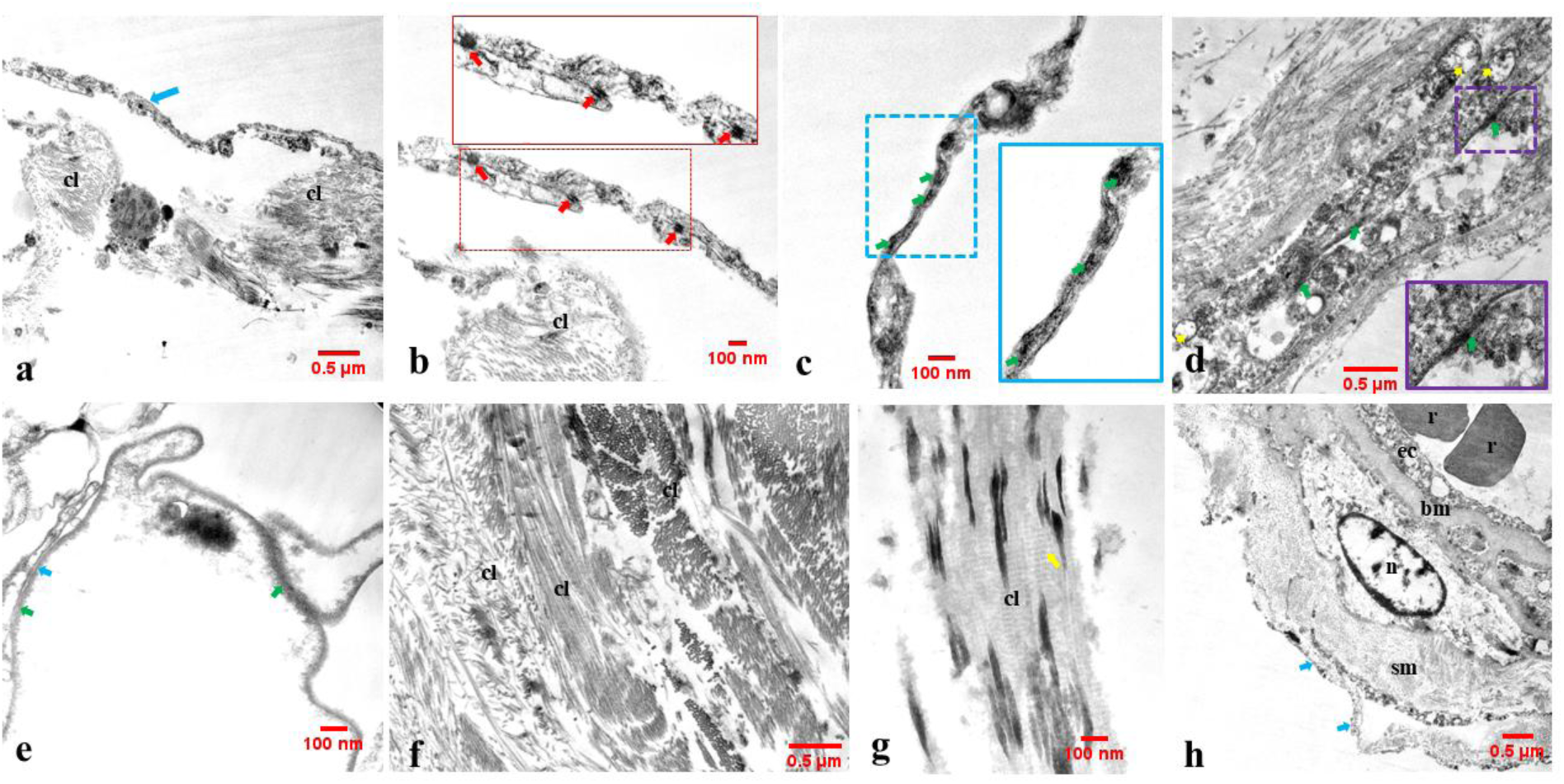
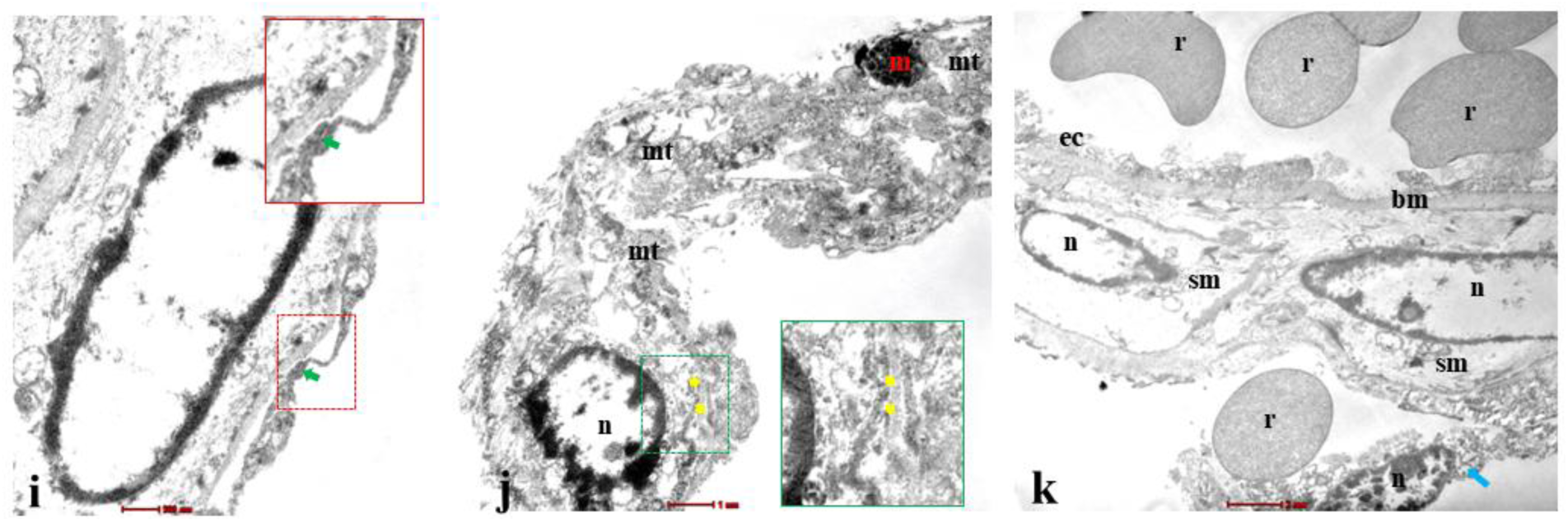
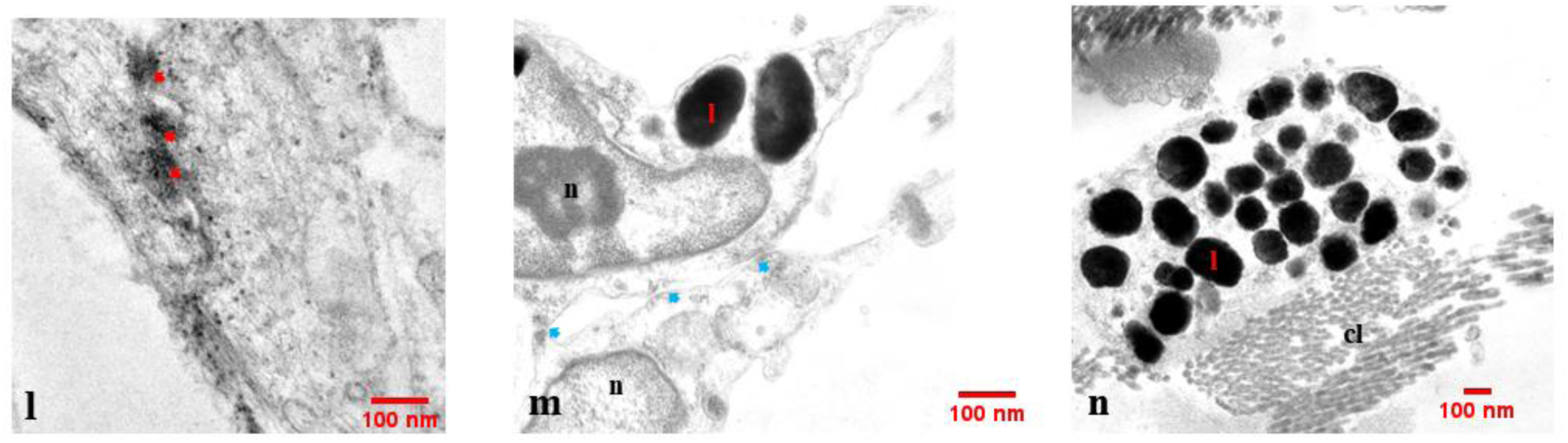
Transmission Electron Microscopy of Adult Human Brain and Spinal Cord. **(a–k)** Transmission Electron Microscopy (TEM) of dissected specimens of the ILL from postmortem adult human brain and spinal cord. **(a–d)** TEM revealed the fibrocellular nature of the ILL. The cellular component is indicated by blue arrows, while the fibrous element is labeled as collagen (cl). In the cortex, the ILL consisted of one to two cell layers. **(b–d)** Adherence and tight junctions were observed in the cortical ILL. Magnified insets show a desmosome (red arrows) and multiple tight junctions (green arrows). Mitochondria are visible within the cells (yellow arrows). **(e)** Pseudopodia-like projections were observed extending from the cells, more prominently in the spinal ILL, contributing to intercellular junctions. The spinal ILL exhibited multiple cell layers, ranging from two to five. Tight and adherens junctions were present between these parallel cell layers (green and blue arrows, respectively). **(f–g)** A magnified view of the fibrous component revealed collagen bundles (cl) oriented in multiple directions (**f**). A characteristic collagen-specific banding pattern was evident (**g**). **(h)** A magnified view of a cortical blood vessel is shown. The lumen of the vessel is identifiable by the presence of red blood cells (r) and an endothelial cell (ec) lining. A large smooth muscle (sm) cell is visible in the vessel wall, distinguished by its large nucleus (n) and numerous actin filaments in the cytoplasm. The muscle layer is separated from the endothelial layer by a thick basement membrane (bm). The ILL forms a perivascular sheath around vessels that traverse this layer (marked by blue arrows). **(i)** An intercellular tight junction is visible (marked by a green arrow and shown in the inset) within the ILL-derived perivascular sheath (green arrow). **(j)** An ILL cell is shown, exhibiting characteristics of a typical meningeal cell. The nucleus (n) is visible. Multiple mitochondrial profiles (mt) are also present, though faint due to postmortem autolysis. The basement membrane, marking the cell boundary, is visible (highlighted with yellow arrows and shown in the inset). Toward the upper end, a mast cell (mc) is seen indenting the cytoplasmic membrane. **(k)** Evidence of blood cell extravasation from an injured vessel is shown. A red blood cell (r) is seen lodged in the perivascular space bounded by the ILL (blue arrow). Signs of injury are present in both the vessel wall and the ILL-derived perivascular sheath. **(l-n)** The desmosomes (red arrows) (l) and adherens junctions (m, indicated by blue arrows) are visible between the cells of the ILL in sections taken from the spinal cord. Lysosomes (ly) are visible in the cell cytoplasm, forming clusters at cellular processes (n), indicating the macrophage-like properties of ILL cells in the spinal cord. **Abbreviations:** cl-collagen, r-rbc, ec-endothelial cell, bm-basement membrane, sm-smooth muscle, n-nucleus, mt-mitochondria, mc-mast cell, ly-lysosome.

### Ultrastructural observations

SEM confirmed the presence of the ILL across the cortex, brainstem, and spinal cord, consistent with gross and histological findings (Fig. 5A–C). It delineated the ILL compartmentalizing the SAS, extending into sulci and fissures, and forming perivascular sheaths, while remaining separated from the arachnoid and pia by intervening trabeculae and neurovascular structures. At higher resolution, the ILL appeared non-sieved, with a smooth inner and relatively rough outer epithelial surface (Fig. 5A, C).

TEM visualization of intact composite specimens was limited by the width of the human SAS; however, imaging of dissected ILL segments from the cortex and spinal cord corroborated the light microscopic findings. These showed a fibrocellular structure with adherens and tight junctions, and a contribution to perivascular sheaths (Fig. 6a–l). Cellular components with macrophage-like features were supported by pseudopodia-like projections and lysosomal clusters (Fig. 6e, m–n).

### Morphometric and quantitative estimations: (Fig. 7: A-C)

#### Comparative thickness of ILL

Based on the morphometric evaluation in SEM images, ILL was found to be thinnest in the cortex (mean ± SE: arachnoid: 25±9 µm; ILL: 15±12 µm, pia: 7.5±5 µm), comparatively thicker in the brainstem (mean ± SE: arachnoid: 65±22.5 µm, ILL: 21±9 µm, pia: 6.5±3 µm), and thickest in the spinal cord (mean ± SE: arachnoid: 48.5±26 µm, ILL: 59±24 µm, pia: 7±3 µm). The thickness of the ILL was significantly less than that of the arachnoid in the cortex and brainstem (p < 0.001) but comparable to that in the spinal cord (p = 0.54). In contrast, the ILL thickness was significantly greater than pia in the cortex, brainstem, and spinal cord (p < 0.001) **(Fig. 7A)**.

### Outer vs. Inner SAS compartment dimensions

As the CSF compartments collapse in fixed histological sections, the circumference of the vessels was used as a proxy to indicate the dimensions of SASo and SASi. The circumference of the vessels located in the SASo was much larger than those in the SASi (mean ± SE: 2094 ± 147.9 μm vs. 165 ± 6.1 μm, fold difference of mean: 12.7, p<0.0001), indicating the outer compartment being much larger than the inner **(Fig. 7B)**.

**Figure 7.**
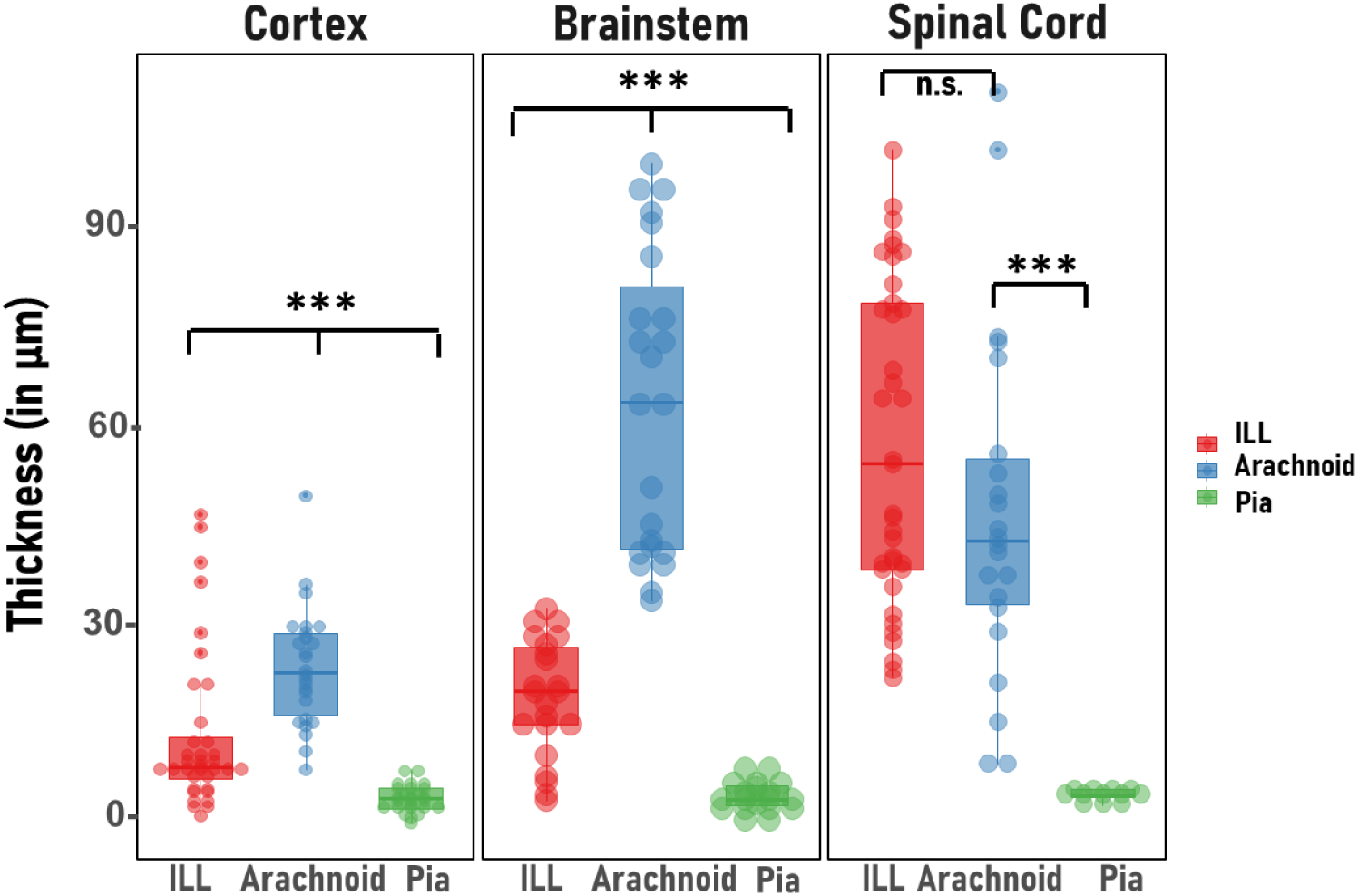
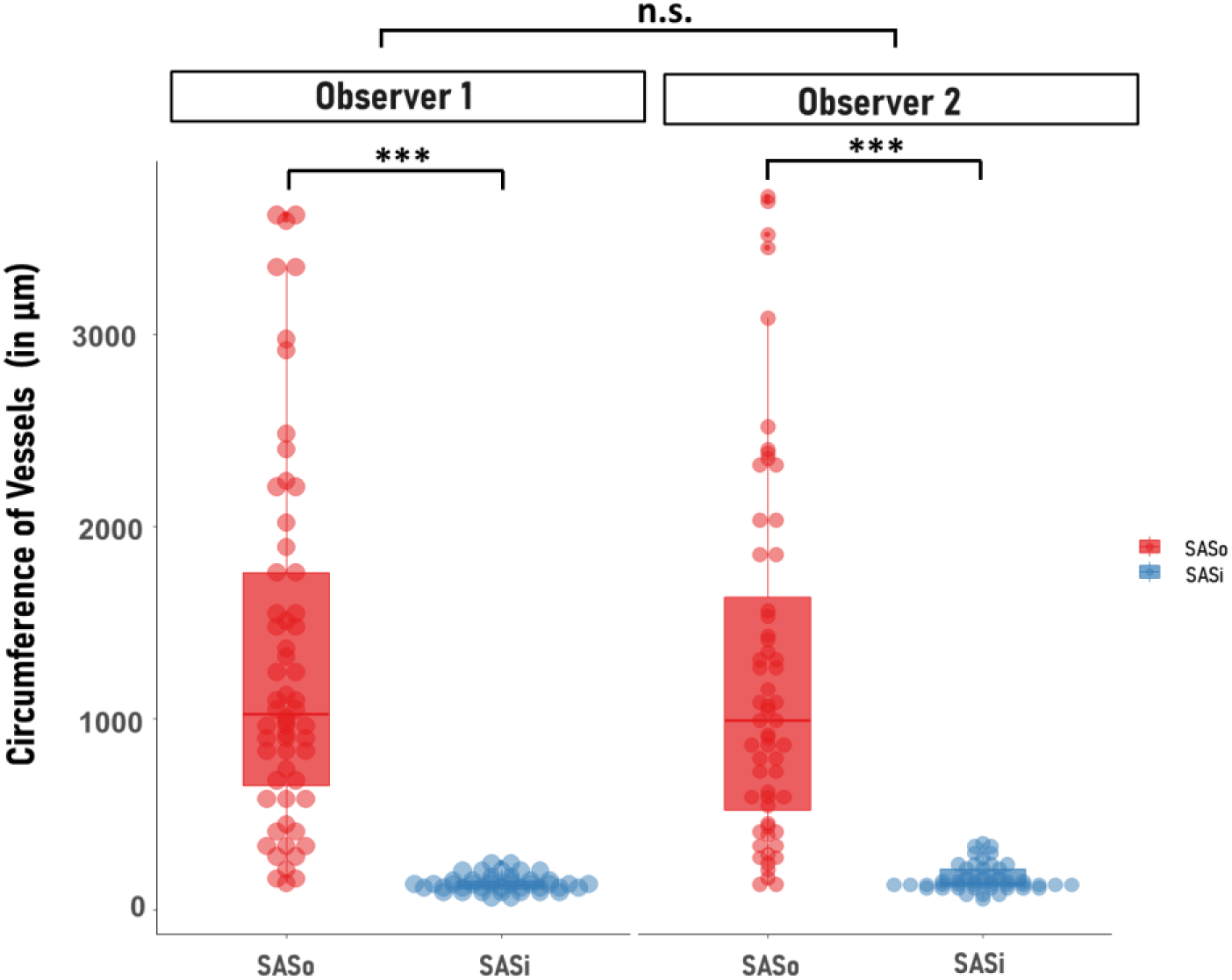
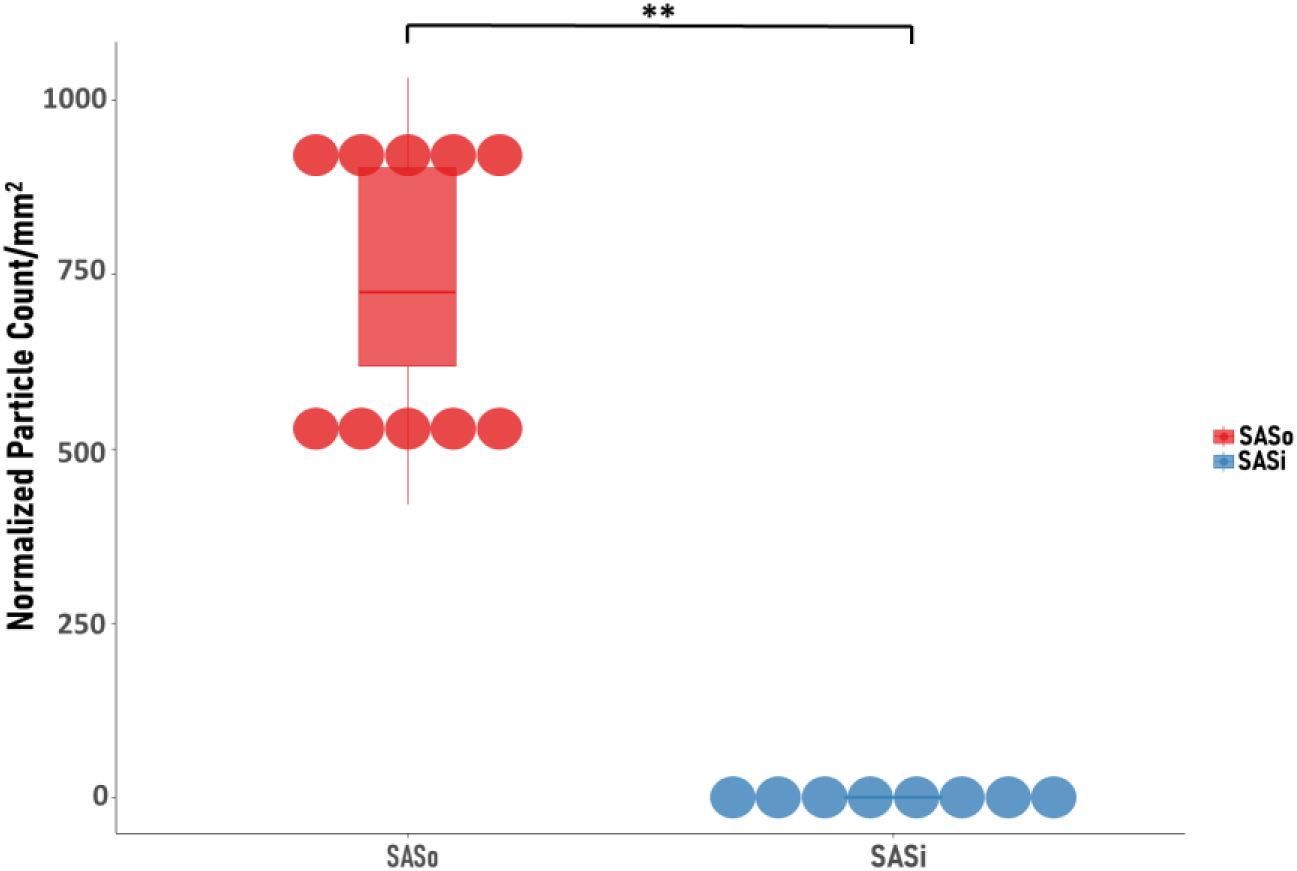
Morphometric and quantitative estimations of ILL. **A. Comparative thickness of leptomeningeal layers across the human CNS.** Raincloud plot showing regional variation in thickness (in µm) across leptomeningeal layers within the cortex, brain stem, and spinal cord. Across the CNS, the arachnoid was the thickest layer, followed by the ILL and pia (p < 0.001). The thickness of the ILL increased progressively from the cortex to the spinal cord (p < 0.001). In the spinal cord, the ILL thickness was observed to be comparable with the arachnoid (p = 0.113, post hoc pairwise comparison). *p≤0.05, **p≤0.01, ***p≤0.001, n.s. p> 0.05. **B. A morphometric assessment of the vascular dimensions in Outer vs. Inner SAS compartments in the adult human CNS**. Raincloud plot showing variation in vessel circumferences between the outer (SASo) and inner (SASi) subarachnoid space compartments in the adult human CNS. A highly significant effect of SAS compartment was observed (Wald Q = 74.496, p < 0.001), whereas no significant observer effect was detected (Wald Q = 0.102, p = 0.750). Vessels within the SASo compartment exhibited significantly larger circumferences than those in the SASi compartment (p < 0.001, post hoc pairwise comparison). These findings indicate marked regional differences in vascular profile dimensions, reflecting similar differences in actual dimensions of the SAS compartments. *p≤0.05, **p≤0.01, ***p≤0.001, n.s. p> 0.05. **B. Distribution of CD68-positive macrophages within outer versus inner SAS compartments following traumatic brain injury**. Raincloud plot showing regional variation in the distribution of CD68-positive macrophages within the outer (SASo) and inner (SASi) subarachnoid space compartments in cases of subarachnoid hemorrhage following traumatic brain injury in human adults. A significant regional disparity in particle concentration was observed (Wilcoxon signed-rank, W = 54.0, p = 0.002), with a very large effect size (rank-biserial correlation = 0.964). The preferential localization of macrophages within specific SAS compartments suggests a potential barrier role of the ILL in limiting macrophage trafficking across the subarachnoid space compartments. *p≤0.05, **p≤0.01, ***p≤0.001, n.s. p> 0.05.

### ILL restricts inter-compartmental immune cell trafficking

We assessed the distribution of CD68-positive immune cells across the outer (SASo) and inner (SASi) compartments in traumatic brain injury and subarachnoid hemorrhage (Fig. 7C). The SASo showed a significantly higher density of CD68-positive particles (737.42 ± 208.13 particles/mm²) compared to the SASi (101.67 ± 218.64 particles/mm²; p = 0.0002). ROI analysis demonstrated consistently high densities in the SASo (420.56–1032.26 particles/mm²), whereas the SASi largely lacked positive staining, with a median density of 0.00 particles/mm² (Fig. 4C: a–b).

These findings indicate that the ILL restricts immune cell distribution between SAS compartments.

## Discussion

The recent claim regarding an ILL in mouse brains raised significant controversy about its true existence (*2, 3, 6*). We investigated the presence of the ILL in human CNS using multimodal imaging ranging from gross examination to ultrastructural analysis. Converging evidence across these scales supports the presence of a continuous ILL along the entire CNS, with region-specific architectural variation. As an independent meningeal layer should be morphologically distinguishable from adjacent layers and extend along the entire CNS, the evidence presented here fulfils these anatomical criteria.

Our macroscopic analyses provide a high-resolution glimpse of the ILL and establish its identity in the human CNS based on its unique morphological characteristics. The ILL divides the SAS into two distinct structural compartments separated by nerves, vessels, and trabeculae (Fig. 1A-D, Supplementary movie: 1-8). The outer SAS is substantially larger, containing coarse trabeculae, large-caliber vessels, and most of the CSF. Whereas, the inner SAS is a narrow slit filled with fine trabeculae, small-caliber vessels, and capillary-layer CSF (Fig. 1B-d-f, Supplementary movie 1-2, 6), except at the cisterns, where a relatively greater amount of CSF may be present (Supplementary movie 7).

The ILL’s unique morphological features can be used to distinguish it macroscopically from the other leptomeningeal layers during the neurosurgical approaches. Although it has a resemblance to the arachnoid, in contrast to the latter, in the cortex, it dips into the sulci and fissures, carrying vessels within its double fold. There, it can be differentiated from the pia by the presence of a CSF-filled space with fine trabeculae between the two (Fig. 1-2, Supplementary movie: 1-8).

Uniquely, a previously under-recognized structure—fine “intra-layer trabeculae”—fills the double fold of the ILL carrying vessels into the depths of the sulci and fissures (Fig. 1B: b, Supplementary movie 1). These intra-layer trabeculae are densest at the sulcal entry points, where they anchor the fold and form a seal that may mechanically preserve sulcal and gyral integrity, and thus overall brain shape.

Compared with the cortex and brainstem in the spinal cord, ILL is significantly thicker, suggesting a possible role in providing mechanical advantages for spinal mobility.

As the spinal surface is smooth, it may be difficult to distinguish ILL from the arachnoid in the cervical and thoracic regions in the fixed cadaveric specimens. However, in the lumbar and sacral region, where spinal nerve roots aggregate and CSF-filled large cisterns are present in between the layers, the two layers can be easily distinguished. A thicker ILL in the spinal cord also helps to differentiate it from the much thinner pia (Fig. 1A: d, C:d-f, Supplementary movie 4).

Leptomeninges are known to develop by the second trimester in human fetuses (*11*). Our observations in examined fetal samples indicated that the ILL develops in parallel with the arachnoid and pia. In second-trimester fetuses, where the cortical sulci were not yet developed, the arachnoid–ILL complex sandwiching the subarachnoid vessels could be easily peeled off, leaving behind a smooth, pia-covered surface (Fig. 1D: a-b). However, in a full-term fetus, where cortical sulci and fissures were well developed, a double fold of the ILL carrying the vessels dipped inward, while the arachnoid bridged over, a pattern also observed in the adult samples (Fig. 1D: c-e). Comprehensive developmental studies are needed to elucidate the early developmental landscape of the ILL.

Our microscopic observations in human CNS validate the lymphatic-like characteristics of ILL, consistent with recent studies in mouse brains (*2*, *3*). We showed a distinct Podoplanin⁺ Prox-1⁺ Lyve-1⁻ Vegfr-3⁻ ILL throughout the human CNS (Fig. 3 A-E). Prox-1 was found to be exclusively expressed by the ILL, thereby representing a key immunophenotypic marker of this layer. Notably, Podoplanin and Prox-1 are characteristically expressed by lymphatic vessel endothelium; therefore, the positivity of the ILL for these markers strongly suggests lymphatic-like properties of this layer. The Podoplanin was also expressed in the pia mater and trabeculae in the inner SAS compartment, indicating their possible mesothelial derivation and developmental proximity.

The ILL seemed to have a basement membrane as it expressed laminin alpha-2 (Lama 2) (Fig. 3F). DPP4 (CD26), recently proposed as an arachnoid-exclusive marker (*12*), was also expressed in the ILL and pia mater (Fig. 3G, S2-o), consistent with evidence demonstrated by Pla et al., 2024 (*3*).

We observed the expression of E-cadherin and Claudin-11, key markers of adherence and tight junctions, respectively, in ILL on repeated examinations (Fig. 4 A-B, S2: f-g, h-i). Of note, Mollgård et al (*2*) described ILL in mice to be a relatively impermeable membrane that did not allow the passage of moieties greater than 1μm in diameter and 3 kDa in mass. However, they showed this layer to be negative for both E-cadherin and Claudin-11. The absence of adherens and tight junctions in their light-microscopic observations made it difficult to determine what caused the impermeability of ILL. Here, we confirmed the presence of an imperforate ILL in SEM-based examinations (Fig. 5A-C: e). Moreover, we showed the presence of these junctions in TEM imaging (a gold standard technique for showing junctional complexes) of dissected ILL specimens from the human cortex and spinal cord (Fig. 6: a-e, i, l).

Interestingly, examining cortical sections from traumatic brain injury cases with subarachnoid hemorrhage in our study revealed that ILL may truly function as a barrier, thereby localizing the extravasation of blood and infiltrating immune cells to the respective space. The compartment-restricted localization of the extravasated blood in the cases of subarachnoid hemorrhage was also indicated in a recent CT-scan-based study (*13*).

The historical TEM-based studies by Krisch et al. on rat brains indicated the compartmentalization of SAS (*14*, *15*). However, the human SAS is too wide to study using TEM. A combined SEM–TEM approach helped overcome this limitation. SEM confirmed the ILL as a non-sieved fibrocellular layer that compartmentalizes the SAS and forms a perivascular sheath, while TEM delineated its detailed fibrocellular architecture and junctional complexes.

The ILL, predominantly in the spinal cord, harboured cells with macrophage-like features, and expressed typical macrophage markers, CD68 and S100 (Fig. 4C-D, S2: l-p). These cells also expressed Lyve-1 and Vegfr-3, which are known to be expressed in macrophages (*16–19*). TEM imaging of the spinal ILL revealed these cells contained pseudopodia-like processes (Fig. 6e) and prominent lysosomal clusters (Fig. 6m–n), consistent with phagocytic activity (Fig. 6m–n). These cells also formed adherent junctions with adjacent cells (Fig. 6m), raising the possibility that they may represent meningeal cells undergoing macrophage-like transformation. However, their precise cellular lineage could not be definitively established at this stage.

The immune function of leptomeninges is an emerging concept, and recent single-cell studies in animals and humans have repeatedly demonstrated border-associated macrophages (BAMs) within the leptomeninges (*20–22*). Our findings indicate a specific role of ILL in this regard. The presence of macrophages in ILL from apparently healthy adults in our study indicates their possible role in physiological immune surveillance. Any increase in their population with aging and in specific clinical conditions requires further elucidation.

Surprisingly, along with CD68, some of these cells co-expressed the common T cell receptor antigen CD3. CD3 is known to be expressed by a subset of macrophages (*23–24*), and CD3^+^CD68^+^ resident macrophages have been previously reported in the mouse liver (*25*); however, we didn’t find any earlier reports of their presence in the meninges. The biological relevance of these cells in ILL is beyond the scope of this study, but this opens several intriguing questions for future research.

Recent studies in mouse brains (*2, 3*) reported that vessels are primarily located within the inner SAS compartment; our observations in human specimens suggest a somewhat different vascular arrangement. In our study, large caliber vessels were more frequently observed in the outer compartment. As vessels traverse from the outer to the inner compartment through the meningeal layers and progressively branch, this distribution appears consistent with the established vascular anatomy of the human head and neck. The relatively limited number of large-calibre vessels in the outer SAS reported in mouse studies may reflect species-specific differences, including the possibility that major cerebral vessels enter the inner SAS earlier in mice.

Any claim of an additional meningeal layer in humans is largely absent from the published literature, except for a sporadic report by Nicholas and Weller (1988) (*5*), who described a sieved ILL in the spinal cord using SEM and light microscopy in cadaveric specimens. In contrast, our high-resolution SEM imaging of fresh postmortem tissue demonstrates a non-sieved ILL, suggesting that the perforations reported in their study may reflect postmortem epithelial degradation.

In contrast to the proponent studies (*2, 3*), two subsequent studies in mice raised strong reservations about the very existence of the ILL (*26*, *27*). These authors, using fluorescence microscopy and TEM demonstrated that the Prox-1^+^ cell layer exists in proximity with the E-cadherin^+^ arachnoid barrier cell (ABC) layer on the outer aspect of the arachnoid; hence, they preferred to call it the inner arachnoid rather than considering it a distinct meningeal layer.

The difficulty in distinguishing a distinct ILL in the mouse CNS may, in part, be attributable to post-fixation adherence to the arachnoid mater or loss of this delicate layer during tissue processing. A narrower outer compartment in the mouse, as we discussed above, may also increase the chances of adherence. When adherent, arachnoid, and ILL may appear in continuity, owing to the presence of continuous trabeculae between them, which bear a covering of meningeal cells. Post-fixation shrinkage may also create overlapping meningeal layers and merge signals in fluorescence microscopy.

Critics also raised concerns that in the recent mouse study (*2*), the neurothelial layer of the dura mater may have been misidentified as the arachnoid mater, and in turn, the actual arachnoid mater as the ILL; hence, an outer compartment might have been artifactually created (*28–29*). To dispel these apprehensions, we carefully removed the intact dura mater during dissection and only included the leptomeningeal layers in microscopic examinations of the cortical tissue. In the spinal cord, we showed an intact dura mater in the SEM, revealing its neurothelial lining along with appropriate identification of the leptomeningeal layers, including ILL (Fig. 5C: a-b).

The ILL consistently ensheathed SAS vessels, particularly arteries. The mouse studies proposing ILL also noted this phenomenon (*2*, *3*). Many other studies have also reported a perivascular sheath around SAS vessels, primarily arteries (*7–9*). The *in vivo* studies in rodents (*8*, *30*) and MRI-based studies in living human subjects using CSF tracers demonstrated the presence of CSF inside the perivascular sheath (*7*). The perivascular sheath may arise from the pia or arachnoid; however, CSF inside the sheath and CSF tracer transport through it to the cisterns suggest that an intermediate layer, perhaps the ILL, forms this sheath (*7–10*).

Conventionally used radiological modalities fall short in distinctly visualising the leptomeningeal layers bordering the CSF compartments. However, ex vivo imaging of formalin-fixed human cadaveric brain using Hierarchical Phase-Contrast Tomography (Hip-CT), an ultra-high-resolution modality capable of visualising anatomical structures at near-microscopic levels (31), appears to corroborate the presence of the ILL. The leptomeningeal arrangement observed in HiP-CT images (Supplementary data, HiP CT annotation: https://doi.org/10.6084/m9.figshare.32587335.v2) closely resembled that demonstrated by SEM in our study (Fig. 5A).

The ILL has considerable physiological and clinical significance, given its strategic location within the SAS, immune properties, and restricted permeability (*2*, *3*). Current concepts of CSF circulation dynamics and CNS protective barriers may require revision to incorporate the role of the ILL (*6*). Its potential involvement in immune surveillance could be critical in defending the CNS against invading pathogens (*6*). It would be valuable to investigate structural and functional changes in this layer associated with ageing, infections, and traumatic brain injury (*6*). Exploring whether dysregulated immune surveillance or traumatic/pathological breaches of this layer contribute to the development of neurodegenerative diseases will be an important area of research (*6*, *32*). Furthermore, the ILL’s contribution to the perivascular sheath indicates a potential role of this layer in the glymphatic system, a mechanism believed to facilitate waste clearance from the CNS (*33*, *34*).

The recent mouse studies highlighted the lymphatic-like characteristics of this layer and suggested its presence in the human brain (*2, 3*). The term “SLYM” aptly reflects its immunophenotypic profile observed in mouse brains, which we further validate in human specimens across the entire neural axis. While further studies may enhance knowledge of the functional aspects of this layer, our study provides robust evidence of its presence as a gross, surgically appreciable structure in humans, comparable to other meningeal layers. In the present study, we use the term “ILL” to denote its anatomical position, consistent with the earlier description of an intermediate leptomeningeal layer in the human spinal cord by Nicholas and Weller (1988) (*5, 6*) and broadly corresponding to the “SLYM” described in mouse brains.

Given these parallel terminologies, and following the tradition of evocative anatomical nomenclature, a unified designation aligned with the existing meningeal framework may be beneficial. In this context, we propose the term *Clara mater*, reflecting the lustrous appearance of this layer in fresh postmortem human specimens, while acknowledging that consensus on nomenclature will require further validation and broad acceptance within the field.

Our study has several limitations. Autolysis in postmortem samples may have obscured certain ultrastructural details in TEM imaging and affected signal precision in immunofluorescence analyses. The inherent autofluorescence in postmortem brain tissue, particularly within the core of blood vessels, represents an additional limitation that may have influenced the fluorescence imaging results. Additionally, we did not examine potential differences in CSF components between the outer and inner SAS compartments. Furthermore, our data on the injured CNS was minimal, and we didn’t examine the composition of ILL at single-cell resolution, highlighting the need for more in-depth exploration of these critical aspects. Despite these limitations, our findings provide robust evidence for an additional, immunophenotypically distinct leptomeningeal layer in humans, with potential neurosurgical relevance and implications for further study in neurological health and disease.

In conclusion, converging macroscopic, microscopic, and ultrastructural evidence supports the presence of an independent, immunophenotypically distinct ILL throughout the human CNS. The relative differences in CSF composition between the outer and inner compartments, along with CSF dynamics within the SAS, as well as the developmental ontogeny, immune properties, and pathological involvement of the ILL, warrant further investigation.

## Acknowledgment

The authors sincerely thank the faculty members, residents, and students of the All India Institute of Medical Sciences (AIIMS), Patna, other AIIMS institutions, medical colleges, and research institutes across India for their valuable feedback and insightful discussions, which contributed to the improvement of this study.

## Author (s) contributions

A.K. conceptualized and designed the study. A.K., A.R., A.D., C.K., and A.A. supervised the project. Data collection and analysis were carried out by A.K., S.K., S.S.Y., R.K., A.S., R.K.N., R.K.J., S.S., A.G., V.P., and C.K., T.C.N., and T.S.R. scrutinized the data. Figures and tables were prepared by A.K. Video recordings were performed by A.K., R.K., and R.K.N., while R.K.N. and A.K did video editing and annotations. The first draft was written by A.K. The final draft was reviewed by C.K., A.A., A.R., A.D., P.K., R.K.N., R.K.J., V.P., M.A., S.B., N.K., M.H., S.N.P., S.S., A.G., T.C.N., and T.S.R. All authors approved the final version of the manuscript.

## Ethics statement

Ethical approval for the study was obtained from the institutional ethics committee following the Helsinki Declaration (Ref. n.: AIIMS/Pat/IEC/2024/1216).

## Data availability statement

The data supporting the findings of this study are available upon reasonable request.

## Supplementary Data

Supplementary movies 1-8: https://doi.org/10.6084/m9.figshare.29375426

Annotation of the revised leptomeningeal arrangement in published HiP CT data: https://doi.org/10.6084/m9.figshare.32587335.v2

Materials and Methods

Figs. S1-2

Tables S1-S3

## Supplementary Data

Supplementary movies 1-8: https://doi.org/10.6084/m9.figshare.29375426 Annotation of the revised leptomeningeal arrangement in published HiP CT data: https://doi.org/10.6084/m9.figshare.32587335.v2 Materials and Methods Figs. S1-2 Tables S1-S3

## Materials & Methods

### i. Materials

Fresh postmortem brain and spinal cord specimens (N=50), 10% formalin-fixed cadaveric specimens (N=6), and human brain and spinal cord specimens without known neuropathology were retrieved. Due to ethical constraints, fresh spinal cord specimens were available only from the upper cervical region; however, complete spinal cords were available from formalin-fixed cadaveric specimens. Additionally, fetal brain specimens (N=5) from stillbirths and abortion cases were included. For each subject, age, sex, cause of death, and medical history were recorded (Table S1). Ethical approval for the study was obtained from the institutional ethics committee following the Helsinki Declaration (Ref. n.: AIIMS/Pat/IEC/2024/1216).

### ii. Methods

#### 1. Gross Dissection and Macroscopic Observations

We retrieved brain and spinal cord specimens with meninges from postmortem and cadaveric adults and human fetuses using standard operative procedures. Intact specimens were macroscopically examined for meningeal arrangement around CNS components through in situ dissection. The stepwise dissection was video-recorded using an ultra-high-resolution (8K) digital camera under standard light settings.

The cranium was opened by cutting through the calvaria. The dural covering was carefully incised and removed, exposing the brain parenchyma covered with leptomeninges. The brain was examined for any visible meningeal pathology, traumatic injury, or hemorrhage. Next, incisions were placed to sever the nerves and vessels near their exit/entry foramina at the skull base. A scalpel was inserted through the foramen magnum to retrieve the upper cervical segments of the spinal cord along with the brain specimen. After detaching neurovascular connections, the brain was carefully extracted with its leptomeningeal coverings intact, leaving the dura mater *in situ*. The brain was then gently washed with running tap water or normal saline before being placed on the dissection table for further examination.

A fine needle was used to pierce the arachnoid mater and enter the subarachnoid space (SAS), preferably at a deeper sulcus. Alternatively, a 2-mL syringe with a fine needle was used to inject air or a vital dye (trypan blue) between the layers. The free spread of air or dye indicated the presence of a potential space and facilitated layer separation during dissection. This method was subsequently applied to semi-fixed postmortem specimens (preserved in 4% paraformaldehyde for 1–2 hours) and stored 10% formalin-fixed cadaveric specimens.

In formalin-fixed cadaveric specimens, the neural axis was incised at the level of the foramen magnum. The cranium and vertebral column were opened, and brain and spinal cord specimens were extracted along with the dural covering using standard dissection protocols. A 2-ml air-filled syringe was used to inject air jets beneath the arachnoid mater bridging a sulcus, which facilitated its separation from the underlying layer. Repeated air injections were applied to further separate the meningeal layers over a larger region.

Additionally, neural tissue samples with leptomeninges were harvested from representative brain and spinal cord regions of postmortem adult subjects for further microscopic analysis. Tissue blocks (2×2 cm) were collected along the entire neural axis, from all lobes of the cerebral cortex, cerebellum, brainstem, and spinal cord.

#### 2. Histological Staining

Hematoxylin and eosin (H&E), Masson’s trichrome, and toluidine blue staining were performed following standard protocols. Stained sections were examined and imaged using a bright-field microscope.

#### 3. Immunohistochemistry (IHC) and Immunofluorescence (IF)

IHC staining was performed using meningeal layer-specific markers (Table S2) following standard protocols. Tissue blocks (2×2 cm) were placed in Falcon tubes containing 4% freshly prepared PFA and fixed at 4°C for 6 hours. After fixation, tissue blocks were washed three times in 0.1 M phosphate buffer (PB) to remove residual fixative and then immersed in 30% sucrose (diluted in 0.1 M PB) for cryopreservation before sectioning.

Sections (10–20 µm thick) were cut using a cryotome. The PFA-fixed tissue samples were embedded in paraffin, and 3–5 µm-thick sections were obtained using a rotary microtome. For paraffin sections, antigen retrieval was performed following standard protocols. Sections were transferred onto Poly-L-lysine-coated glass slides and immunostained. Counterstaining with hematoxylin or eosin was performed to differentiate the structures of interest. Stained sections were imaged using a bright-field microscope.

For IF analysis, Alexa fluorophores (table S2) conjugated to secondary antibodies were used for staining, and slides were examined under a Stellaris 5 confocal laser microscopic platform by Leica.

To minimise technical and imaging-related artefacts, all tissue sections within each experimental set were processed under identical staining conditions, including fixation, antigen retrieval, antibody incubation, washing, and mounting procedures. Fluorescence imaging was performed using identical acquisition parameters for comparative samples, including laser intensity, detector gain, exposure time, pinhole size, and scanning speed, to ensure consistency across experiments. The negative controls, including omission of the primary antibodies, were incorporated into each staining run to assess tissue autofluorescence, background signal, and non-specific binding of the secondary antibody. To reduce spectral overlap and channel cross-talk, spectrally distinct fluorophores were selected and sequential channel acquisition was employed during image capture.

#### 4. Scanning Electron Microscopy (SEM)

Tissue blocks (1×1 cm) were fixed in modified Karnovsky solution (2.5% glutaraldehyde and 2% paraformaldehyde in 0.1 M PB, pH 7.4) at 4°C for 6 hours. Samples were washed three times with PB (15 minutes each) and dehydrated in a graded ethanol series (40%, 50%, 70%, 90%, and 100% for 5 minutes each). After dehydration, samples were critically point-dried, mounted onto aluminium stubs with the structure of interest facing upward, and sputter-coated with colloidal gold. Specimens were analyzed using an EVO18 scanning electron microscope (Carl Zeiss) at an operating voltage of 15 kV.

#### 5. Transmission Electron Microscopy (TEM)

For TEM, tissue samples were cut into 2 × 2 mm pieces and fixed in 2.5 % glutaraldehyde and 2% paraformaldehyde in 0.1 M phosphate buffer (pH 7.4) at 4°C for 6 hours. The samples were then washed and post-fixed in 1% osmium tetroxide (OsO₄) for 1 hour at 4°C.

Following fixation, the tissue was dehydrated in acetone, followed by treatment with dry acetone (absolute acetone containing anhydrous copper sulfate) for 30 minutes. The samples were then cleared with toluene for 15 minutes.

Infiltration was performed under vacuum using a toluene-araldite (CY212) mixture in the following sequence: 3:1 (toluene to araldite) for 2 hours, 1:1 for 2 hours, 1:3 overnight, and finally, pure araldite for 2–4 hours. The samples were then embedded in pure araldite at 50°C overnight, followed by polymerization at 60°C for 72 hours.

Semithin sections (0.5–1 µm) were cut and stained with toluidine blue, while ultrathin sections (60–70 nm) were cut, mounted on copper grids, and stained with aqueous uranyl acetate and alkaline lead citrate for 5 minutes each. The sections were then rinsed in distilled water and examined using a TALOS F200S TEM (Thermo Fisher Scientific, Waltham, MA, USA) at magnifications ranging from 20,000× to 50,000×.

The identification of intercellular junctions in TEM images was based on the established criterion (*32–34*)(Table S3).

#### 6. Morphometry and particle density quantifications

The morphometric measurements and particle density quantifications were performed on microscopic images using the Image J/Fiji(NIH, USA). Measurements were obtained independently by two investigators, with inter-rater agreement predefined as an Intraclass Correlation Coefficient (ICC) > 0.90. Data from both investigators were compiled, and mean ± standard error (SE) values were calculated, and group differences were statistically analyzed (see Data Collection & Statistical Analysis Plan).

##### The comparative thickness of the ILL

The comparative thickness of ILL and other leptomeningeal layers was studied in SEM images of cortex, brainstem, and spinal cord. Measurements were taken at a minimum of eight locations per meningeal layer in each image across multiple images, and mean values were compared statistically.

##### Dimensions: Outer vs. Inner subarachnoid space

As the CSF compartments collapse in the fixed histological sections, the circumference of the vessels was measured, which is a relatively fixed parameter. The circumference of the vessels was used as a proxy to indicate the dimensions of SASo and SASi. We used ImageJ to trace and measure the total circumference of the vessels in the outer and inner compartments in histological images.

##### CD-68 positive immune cell density: Outer vs. Inner subarachnoid space

We assessed the distribution of CD-68 positive immune cells across the outer (SASo) and inner (SASi) compartments in cases of traumatic brain injury and subarachnoid hemorrhage. The mean density of CD-68 positive immune cells (particles/mm^2^) were measured using Image J/Fiji in DAB-stained images, and comparative differences across the SAS compartments were statistically analyzed.

### iii. Data Collection and Analysis

#### a. Analysis of Gross Dissection and Microscopic Data

Three subject experts annotated gross dissection and microscopic findings. The interpretation was based on established anatomical criteria (*1*):

1. The arachnoid mater is the outermost leptomeningeal layer and forms the external boundary of the CSF-filled SAS. It is a thick, multilayered membrane, distinctly separable from the dura mater.
2. The pia mater, the innermost meningeal layer, is a single-cell-thick membrane that adheres to the CNS and follows its contours. Attempting to separate it risks damaging neural tissue.
3. The arachnoid mater bridges over sulci and fissures, while the pia mater follows the neural surface into sulci and fissures.
4. Any intermediate meningeal layer must be anatomically distinct from the pia and arachnoid mater.
5. Trabeculae connect the leptomeningeal layers, enclosing the CSF space.
6. The presence of trabeculae, blood vessels, and intact CSF spaces confirms anatomical distinction and rules out artificial separation during dissection.

#### b. Image Processing and Analysis

Image analysis and annotations were performed using ImageJ/Fiji and Adobe Photoshop/Illustrator (2026). For antibody expression analyses using DAB staining and fluorophores, background-corrected staining scores and normalized fluorescence intensity values, respectively, were used to minimise the influence of non-specific signal variability between sections. Tissue autofluorescence was assessed during control imaging and excluded during analysis wherever necessary. Post-acquisition image processing was restricted to uniform adjustments of brightness and contrast applied equally across all comparative images without selective regional enhancement.

##### Immunohistochemistry (IHC)

**DAB staining scores:** The readings were taken for each histological slide to generate staining scores. The Mean Gray Value’ for each antibody was collected for the six regions (Arachnoid, ILL, Pia, IT-inner trabeculae, OT-outer trabeculae, and Background) across the ROIs in multiple histological sections. DAB staining intensity was calculated by converting ‘Mean Gray Value’ into ‘Optical Density (OD)’ using the formula OD=log 10 (Max Intensity/Mean Gray Value). Where, Max Intensity = 255 (for an 8-bit image) and Mean Gray Value = The average pixel intensity of the stained area. Final Background-Corrected Staining Score (ΔOD) was calculated using the formula ΔOD=OD _target_ −OD _background_.

##### Immunofluorescence (IF)

The Mean Fluorescent Intensity readings were taken for each antibody to generate staining scores. The ‘Normalised Fluorescence Intensity’ was calculated by subtracting the minimum value from the raw value of Mean Fluorescent Intensity measured at selected regions of interest (ROIs), then dividing by the difference between the maximum and minimum values. Coloc 2 (ImageJ/Fiji) was used for colocalization analysis. For visualisation purposes, a Gaussian blur (σ=1) was applied to reduce autofluorescence from secretions and blood cells inside the vessel lumen.

#### c. Statistical Analysis

All quantitative measurements, including DAB staining intensity, CD68-positive particle counts, and meningeal layer thickness, and SAS dimensions, were normalised and grouped according to predefined biological factors (e.g., anatomical compartment, cell type, or region). Data distributions were assessed using the Shapiro–Wilk test, and outliers were identified using the Interquartile Range method.

For DAB staining results in IHC, background-corrected staining intensity scores were compared across compartments or cell types using robust ANOVA (one-way or two-way, as appropriate), with interaction effects evaluated where relevant. The significant effects were followed by post hoc pairwise comparisons using trimmed mean differences (psi-hat).

For colocalization analysis of confocal IF images, signal correlation was quantified using Pearson’s correlation coefficient (R) and thresholded Pearson’s coefficient following Costes automatic thresholding. Signal overlap was assessed using thresholded Manders’ coefficients (tM1, tM2). Statistical significance of colocalization was determined by Costes randomization (≥100 iterations), with P ≥ 0.95 indicating non-random spatial association.

Vessel size measurements were analyzed using the same robust ANOVA framework, with model structure determined by the experimental design. Meningeal thickness was evaluated using robust two-way ANOVA with meningeal layer and anatomical region as factors, including interaction effects, followed by post hoc comparisons.

SAS dimensions based on vessel circumference differences were analyzed using robust two-way ANOVA with compartment and observer as factors to assess regional differences and inter-observer reliability.

In contrast, normalized CD68-positive particle counts across SAS compartments, characterized by small sample size and zero-inflated non-normal distributions, were analyzed using the Wilcoxon signed-rank test, with effect size quantified using rank-biserial correlation.

Effect sizes are reported alongside p-values, including bootstrap-derived 95% confidence intervals where appropriate. All tests were two-sided, with statistical significance defined as p < 0.05, and data distributions with individual variability were visualized using raincloud plots to ensure transparency. All analyses were performed using Jamovi (version 2.7.18) and R (version 4.6.0).

**Figure S1.**
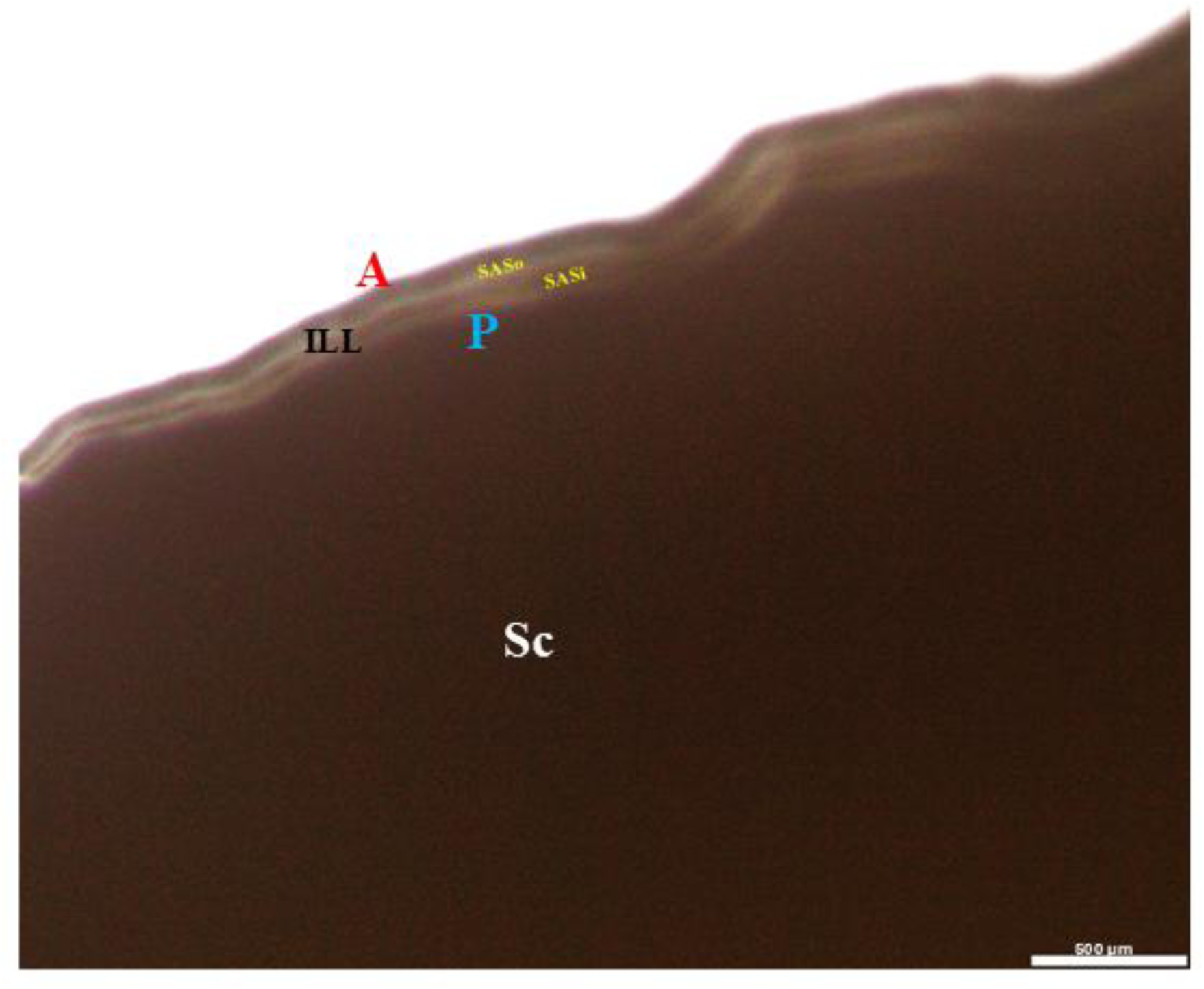
The light microscopic imaging of unstained thick sections of the subarachnoid space in the human spinal cord. Unstained thick sections (20 µm) of subarachnoid space (SAS) specimens from the adult human spinal cord (Sc) were examined under a bright-field light microscope. The transmitted light enabled visualization of the meningeal layers, allowing distinction between the outermost arachnoid mater (A), the ILL, and the innermost pia mater (P). The ILL divided the SAS into outer (SASo) and inner (SASi) compartments.

**Figure S2.**
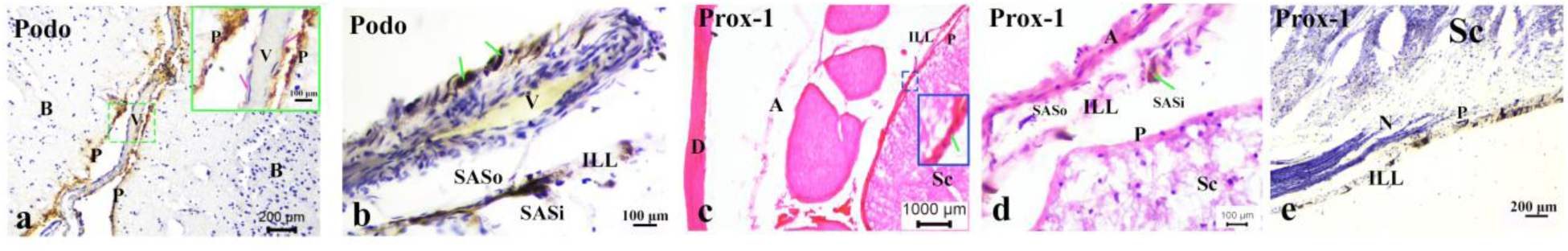
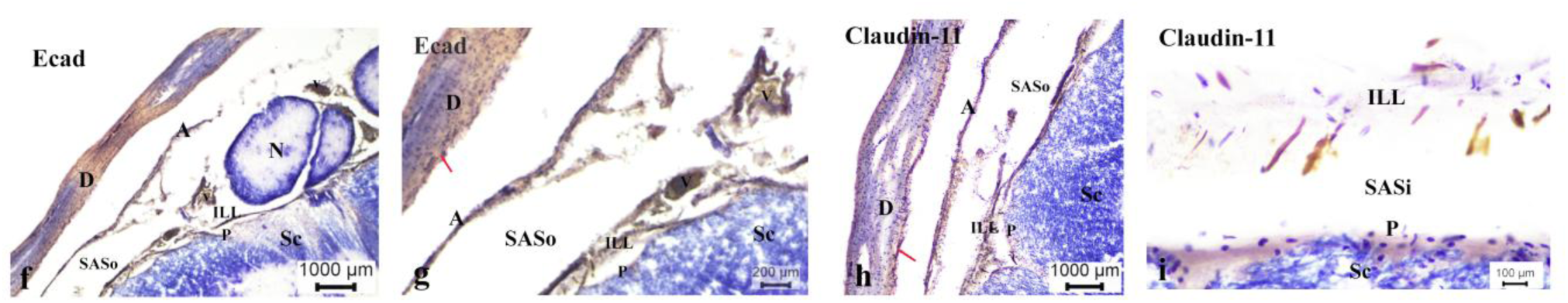
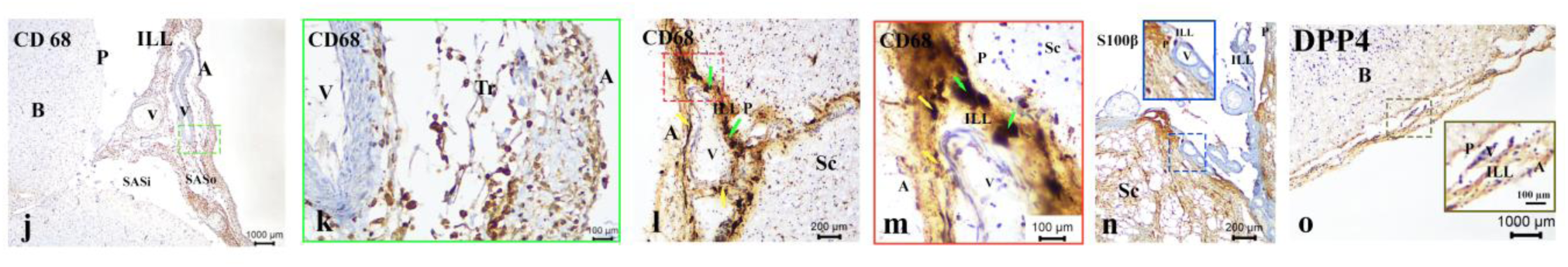
Immunohistological expression of meningeal markers in adult human brain and spinal cord. (Staining with diaminobenzidine (DAB) and counterstaining with haematoxylin/eosin) **(a–b) Podoplanin** In the cortical sulcus of the brain (B), the ILL and pia mater (P) exhibited positive podoplanin staining (a). The ILL created the perivascular sheath (b, green arrows). In the spinal cord, podoplanin-positive perivascular sheaths were observed surrounding vascular profiles in the SASo (b, green arrows). **(c–e): Prox-1** The dura mater (D), arachnoid mater (A), and pia mater (P) showed no Prox-1 expression, whereas the ILL exhibited positive staining (c, counterstained with eosin). The ILL divided the subarachnoid space (SAS) into outer (SASo) and inner (SASi) compartments. In certain regions, the ILL adhered to the underlying pia, completely obliterating the SASi. A magnified view of the inset showing the distinction of ILL from pia (c). Cross-sections of spinal nerve rootlets (N) were observed between the ILL and arachnoid (c). The ILL also formed a perineural sheath around spinal nerve rootlets as they passed through it (e). **(f-g): E-cadherin** The meningeal arrangement, including the dura mater, was visualized in the adult spinal cord (Sc). E-cadherin expression was evident in the arachnoid (A) and the ILL. The ILL divided the SAS into outer (SASo) and inner (SASi) compartments. At multiple places, the ILL adheres to the pia (P). Multiple vascular profiles and cross-sections of spinal nerve rootlets (N) were present between the arachnoid and ILL. Focal linear E-cadherin expression was noted within the dura, suggestive of meningeal lymphatics or intradural vessels (red arrow). **(h-i): Claudin-11** The meningeal arrangement, including the dura mater, was visualized in the adult spinal cord (Sc). Claudin-11 expression was evident in the arachnoid (A) and the ILL. The ILL divided the subarachnoid space (SAS) into outer (SASo) and inner (SASi) compartments. In the dura mater, Claudin-11 expression was noted in the neurothelial border (red arrow). The pia mater (P) stained negatively for Claudin-11 (i). **(j-o): CD68** **(j–k) Injured cortex**: **j.** In a case of traumatic brain injury where death occurred a few days after the incident, subarachnoid hemorrhage and dilated blood vessels were noted. The extravasated blood and infiltrating immune cells (mononuclear phagocytes) were confined between the arachnoid mater (A) and the ILL, i.e., in the outer subarachnoid space (SASo). Extensive CD68 expression was evident in this region. **k.** Magnified inset images from (j), focused towards the arachnoid (A). **(l-o): Injured Spinal Cord** **l.** A case of acute traumatic spinal cord injury (4-6h). The view was taken at the anterior median sulcus. Three distinct leptomeningeal layers are observed. The engorged and dilated vessels are visible between the arachnoid (A) and the ILL. The double fold of ILL is seen entering the depth of the fissure. Intense CD68 expression is observed in the ILL layer (green arrows) and the perivascular sheath (yellow arrows). **m.** An inset from (l) distinctly shows the CD 68 positive macrophage-like cells with large nuclei and cytoplasmic debris in the ILL layer (green arrows). The ILL is distinctly separable from the underlying Pia (P). CD 68 expression is also present in the perivascular sheath around the vessels present between ILL and arachnoid (A) (yellow arrows). **n. S100β** Views were taken at the anterior median fissure of a non-injured spinal cord. S100β, a macrophage marker, was expressed in the ILL, with unstained meningeal cells interspersed. At multiple places, the ILL adhered to; however, a distinct pia (P) can be marked out. **o.** DPP4 expression was positively detected in each leptomeningeal layer on the cortical surface of the brain (B), from outer to inner: arachnoid (A), ILL, and pia (P). Abbreviations: D-dura, A-arachnoid, P-pia, ILL-intermediate leptomeningeal layer, Tr-trabeculae, N-nerve, V-vessel, SAS-subarachnoid space, i-internal, o-outer. B-brain, Sc-spinal cord.

**Table S1.** Phenotypic and clinical details of the samples.

| **Fresh postmortem adult samples** |  |  |  |  |  |
| --- | --- | --- | --- | --- | --- |
| **S. No.** | **Age (In Years)** | **Sex** | **Cause of death** | **PM delay (In Hours)** | **Clinical history** |
| 1. | 52 | Male | Road traffic accident | 16 | No clinical history reported |
| 2. | 68 | Male | Road traffic accident | 17 | No clinical history reported |
| 3. | 17 | Male | Road traffic accident | 17 | No clinical history reported |
| 4. | 70 | Male | Road traffic accident | 9 | No clinical history reported |
| 5. | 54 | Male | Road traffic accident | 6 | No clinical history reported |
| 6. | 33 | Male | Road traffic accident | 16 | No clinical history reported |
| 7. | 40 | Male | Suicide by poisoning | 17 | No clinical history reported |
| 8. | 27 | Male | Electrocution | 17 | No clinical history reported |
| 9. | 50 | Female | Unexplained medical reason | 20 | Malnutrition |
| 10. | 37 | Male | Electrocution | 20 | No clinical history reported |
| 11. | 18 | Male | Suicide by hanging | 12 | No clinical history reported |
| 12. | 32 | Male | Drowning | 24 | Psychiatric illness |
| 13. | 28 | Male | Road traffic accident | 4 | No clinical history reported |
| 14. | 19 | Male | Road traffic accident | 6 | No clinical history reported |
| 15. | 80 | Male | Unknown cause of death | 12 | Malnutrition |
| 16. | 33 | Male | Electrocution | 18 | No clinical history reported |
| 17. | 22 | Female | Suicide by hanging | 10 | No clinical history reported |
| 18. | 21 | Male | Suicide by poisoning | 17 | No clinical history reported |
| 19. | 32 | Male | Road traffic accident | 22 | No clinical history reported |
| 20. | 40 | Male | Myocardial infarction | 22 | Chest pain |
| 21. | 25 | Male | Road traffic accident | 5 | No clinical history reported |
| 22. | 35 | Female | Suicide by hanging | 20 | No clinical history reported |
| 23. | 41 | Male | Suicide by poisoning | 14 | No clinical history reported |
| 24. | 44 | Male | Heat stroke | 15 | No clinical history reported |
| 25. | 40 | Male | Heat stroke | 16 | No clinical history reported |
| 26. | 45 | Female | Suspected heat stroke | 29 | Thyroid medication |
| 27. | 25 | Male | Suicide by hanging | 24 | No clinical history reported |
| 28. | 57 | Male | Heat stroke | 5 | No clinical history reported |
| 29. | 45 | Male | Heat stroke | 24 | No clinical history reported |
| 30. | 45 | Male | Road traffic accident | 12 | No clinical history reported |
| 31. | 35 | Male | Road traffic accident | 18 | Lesions in the liver, under medication |
| 32. | 60 | Male | Suspected heat stroke | 24 | No clinical history reported |
| 33. | 4 | Female | Road traffic accident | 6 | No clinical history reported |
| 34. | 17 | Male | Road traffic accident | 9 | No clinical history reported |
| 35. | 30 | Female | Suicide by hanging | 24 | No clinical history reported |
| 36. | 28 | Female | Suicide by hanging | 24 | Psychiatric Illness |
| 37. | 30 | Female | Suicide by poisoning | 20 | No clinical history reported |
| 38. | 57 | Male | Road traffic accident | 12 | No clinical history reported |
| 39. | 30 | Male | Road traffic accident | 13 | No clinical history reported |
| 40. | 28 | Female | Suicide by hanging | 7 | No clinical history reported |
| 41. | 56 | Female | Road traffic accident | 15 | No clinical history reported |
| 42. | 18 | Male | Suicide by poisoning | 24 | No clinical history reported |
| 43. | 06 | Female | Road traffic accident | 24 | No clinical history reported |
| 44. | 50 | Male | Road traffic accident | 7 | No clinical history reported |
| 45. | 75 | Male | Road traffic accident | 6 | Hypertension |
| 46. | 24 | Female | Suicide by hanging | 15 | No clinical history reported |
| 47. | 2.5 | Male | Road traffic accident | 12 | No clinical history reported |
| 48. | 40 | Male | Road traffic accident | 7 | No clinical history reported |
| 49. | 35 | Male | Firearm injury | 24 | No clinical history reported |
| 50. | 20 | Female | Suicide by hanging | 9 | Suicidal tendency. No other clinical history reported |
| **Cadaveric adult samples\*** |  |  |  |  |  |
| 51. | >18 | Not known | Not known | Not known | No visible developmental anomalies or pathology |
| 52. | >18 | Not known | Not known | Not known | No visible developmental anomalies or pathology |
| 53. | >18 | Not known | Not known | Not known | No visible developmental anomalies or pathology |
| 54. | >18 | Not known | Not known | Not known | No visible developmental anomalies or pathology |
| 55. | >18 | Not known | Not known | Not known | No visible developmental anomalies or pathology |
| 56. | >18 | Not known | Not known | Not known | No visible developmental anomalies or pathology |

\*51-56: Well-preserved, 10% formalin-fixed brain and spinal cord specimens from adult voluntary donors (>18 years) were used for macroscopic observations.

Fetal samples
| S. No. | Age (In Weeks) | Sex | Cause of death | PM delay (In Hours) | Clinical history |
| --- | --- | --- | --- | --- | --- |
| 1. | 24 | Female | Medical termination of pregnancy | 3 | Cardiac anomalies |
| 2. | 40 | Female | Medical termination of pregnancy | 2 | Acute fatty liver of pregnancy, pre-eclampsia to mother |
| 3. | 23 | Male | Intra-uterine death | 3 | Neurological anomalies -Spinal bifida |
| 4. | 16 | Female | Intra-uterine death | 2 | Placental disruption and hemorrhage in the first trimester |
| 5. | 20 | Female | Intra-uterine death | 4 | Umbilical cord strangulation around neck |

**Table S2.** List of Antibodies used.

| S.N. | Name | Source | Identifier | Concentration |
| --- | --- | --- | --- | --- |
| 1. | Anti-Podoplanin, Rabbit monoclonal antibody | Abcam Plc, UK | Catalog no. EPR7072 | 1:500 |
| 2. | Podoplanin (LpMab-12) Mouse monoclonal antibody | Cell signaling technology, CA, USA | Catalog no. 26981S | 1:500 |
| 3. | Prox-1 (FM-79) Mouse monoclonal antibody | Santa Cruz Biotechnology Inc., CA, USA | Catalog no. Sc-81983 | 1:500 |
| 4. | Prox-1 (D2J6J) Rabbit monoclonal antibody | Cell signaling technology, CA, USA | Catalog no. 14963S | 1:500 |
| 5. | E-Cadherin (24E10) Rabbit monoclonal antibody | Cell signaling technology, CA, USA | Catalog no. 3195S | 1:400 |
| 6. | Claudin-11 (D-8) Mouse monoclonal antibody | Santa Cruz Biotechnology Inc., CA, USA | Catalog no. Sc-271232 | 1:250 |
| 7. | Lyve-1 (E9VA4) Rat monoclonal antibody | Santa Cruz Biotechnology Inc., CA, USA | Catalog no. Sc-65647 | 1:500 |
| 8. | Anti-Mo-VEGF Receptor 3 | Thermo Fisher Scientific, MA, USA | Catalog no. 14-5988-82 | 1:250 |
| 9. | CD68 (D4B9C) XP(R) Rabbit mAb | Cell signaling technology, CA, USA | Catalog no. 76437S | 1:500 |
| 10. | S100, Mouse polyclonal antibody | Roche Diagnostics, CA, USA | Catalog no. 790-2914 | 1:100 |
| 11. | S100B Mouse monoclonal antibody (SH-B4) | Thermo Fisher Scientific, MA, USA | Catalog no. MA1-25005 | 1:100 |
| 12. | Anti-Hu CD3, eBioscience, Mouse monoclonal antibody | Thermo Fisher Scientific, MA, USA | Catalog no. 14-0038-82 | 1:250 |
| 13. | CD26 (DPP4) Mouse monoclonal antibody | Thermo Fisher Scientific, MA, USA | Catalog no. MA2607 | 1:100 |
| 14. | Goat anti-Mouse IgG (H+L) Highly Cross-Adsorbed Secondary Antibody, Alexa Fluor™ 488 | Thermo Fisher Scientific, MA, USA | Catalog no. A-11029 | 1:1000 |
| 15. | Goat anti-Rabbit IgG (H+L) Cross-Adsorbed Secondary Antibody, Alexa Fluor™ 594 | Thermo Fisher Scientific, MA, USA | Catalog no. A-11012 | 1:1000 |
| 16. | Peroxidase AffiniPure® Goat Anti-Mouse IgG (H+L) Secondary Antibody Horseradish Peroxidase conjugate | Jackson ImmunoResearch Laboratories, CA, USA | Catalog no. 115-035-003 | 1:1000 |
| 17. | Peroxidase AffiniPure™ Goat Anti-Rabbit IgG (H+L) Secondary Antibody Horseradish Peroxidase conjugate | Jackson ImmunoResearch Laboratories, CA, USA | Catalog no. 111-035-003 | 1:1000 |

**Table S3.** Identifying features of intercellular junctions*.

| Type | Alternative names | Membrane characteristics | Inter-cellular space | Cytoplasmic features |
| --- | --- | --- | --- | --- |
| Tight junction (TJ) | Zonula occludens | Outer membranes of adjacent membranes fuse | Absent at the point of fusion | Distinct junctional complex |
| Adherens Junction (AJ) | Zonula adherens—<br>A belt-like structure/Macula adherens or desmosome—<br>A spot or button-like structure | Plasma membranes remain separated/<br>Electron-dense material fills in the space | Filled with electron-dense material | Electron-dense material extends to the cytoplasmic surface |
| Gap junction | Nexus | Outer leaflets are strictly parallel | Narrow but uniform gaps | Localized cytoplasmic densities |
| *Ref (1-3).<br>References<br>1. L. A. Staehelin, Structure and Function of Intercellular Junctions. *Int. Rev. Cytol.* **39**, 191–283 (1974).<br>2. M. G. FARQUHAR, G. E. PALADE, Junctional complexes in various epithelia. *J. Cell Biol.* **17**, 375–412 (1963).<br>3. M. W. Brightman, T. S. Reese, Junctions between intimately apposed cell membranes in the vertebrate brain. *J. Cell Biol.* **40**, 648–677 (1969). |  |  |  |  |

## References

1. Standring, S. Gray’s anatomy: the anatomical basis of clinical practice 42nd edition, 955–985 (Elsevier, London, 2021).

2. Mollgård, K. et al. A mesothelium divides the subarachnoid space into functional compartments. Science 379, 84–88 (2023).

3. Plá, V. et al. Structural characterization of SLYM—a 4th meningeal membrane. Fluids Barriers CNS 20, 1–18 (2023).

4. Rasmussen, M. K. et al. Trigeminal ganglion neurons are directly activated by influx of CSF solutes in a migraine model. Science 385, 80–86 (2024).

5. Nicholas, D. S. & Weller, R. O. The fine anatomy of the human spinal meninges: a light and scanning electron microscopy study. J. Neurosurg. 69, 276–282 (1988).

6. Kumar, A. et al. Anatomical correlates for the newly discovered meningeal layer in the existing literature: a systematic review. Anat. Rec. 308, 191–210 (2025).

7. Eide, P. K. & Ringstad, G. Functional analysis of the human perivascular subarachnoid space. Nat. Commun. 15, 1–14 (2024).

8. Mestre, H. et al. Periarteriolar spaces modulate cerebrospinal fluid transport into brain and demonstrate altered morphology in aging and Alzheimer’s disease. Nat. Commun. 13, 913 (2022).

9. Hannocks, M. J. et al. Molecular characterization of perivascular drainage pathways in the murine brain. J. Cereb. Blood Flow Metab. 38, 669–686 (2018).

10. Ringstad, G. & Eide, P. K. Glymphatic-lymphatic coupling: assessment of the evidence from magnetic resonance imaging of humans. Cell. Mol. Life Sci. 81, 1–14 (2024).

11. Dasgupta, K. & Jeong, J. Developmental biology of the meninges. Genesis 57, e23288 (2019).

12. Siegenthaler, J. & Betsholtz, C. Commentary on “Structural characterization of SLYM—a 4th meningeal membrane”. Fluids Barriers CNS 21, 1–7 (2024).

13. Almohaimede, K. et al. Assessing the subarachnoid space anatomy on clinical imaging: utilizing normal and pathology to understand compartmentalization of the subarachnoid space. Acta Neurochir. 167, 1–5 (2025).

14. Krisch, B., Leonhardt, H. & Oksche, A. Compartments and perivascular arrangement of the meninges covering the cerebral cortex of the rat. Cell Tissue Res. 238, 459–474 (1984).

15. Krisch, B., Leonhardt, H. & Oksche, A. The meningeal compartments of the median eminence and the cortex: a comparative analysis in the rat. Cell Tissue Res. 228, 597–640 (1983).

16. Brezovakova, V. & Jadhav, S. Identification of Lyve-1 positive macrophages as resident cells in meninges of rats. J. Comp. Neurol. 528, 2021–2032 (2020).

17. Siret, C. et al. Deciphering the heterogeneity of Lyve1+ perivascular macrophages in the mouse brain. Nat. Commun. 13, 7366 (2022).

18. Schoppmann, S. F. et al. Tumor-associated macrophages express lymphatic endothelial growth factors and are related to peritumoral lymphangiogenesis. Am. J. Pathol. 161, 947–956 (2002).

19. Kannan, S. & Rutkowski, J. M. VEGFR-3 signaling in macrophages: friend or foe in disease? Front. Immunol. 15, 1349500 (2024).

20. Sun, R. & Jiang, H. Border-associated macrophages in the central nervous system. J. Neuroinflammation 21, 1–18 (2024).

21. Zhan, T. et al. Border-associated macrophages: from embryogenesis to immune regulation. CNS Neurosci. Ther. 30, e70105 (2024).

22. Schonhoff, A. M. et al. Border-associated macrophages mediate the neuroinflammatory response in an alpha-synuclein model of Parkinson disease. Nat. Commun. 14, 1–16 (2023).

23. Beham, A. W. et al. A TNF-regulated recombinatorial macrophage immune receptor implicated in granuloma formation in tuberculosis. PLoS Pathog. 7, e1002375 (2011).

24. Ocaña-Guzmán, R. et al. Murine RAW macrophages are a suitable model to study CD3 signaling in myeloid cells. Cells 11, 1635 (2022).

25. Wang, C. et al. Potential role of liver-resident CD3+ macrophages in HBV clearance in a mouse hepatitis B model. JHEP Rep. 7, 101323 (2024).

26. Mapunda, J. A. et al. VE-cadherin in arachnoid and pia mater cells serves as a suitable landmark for in vivo imaging of CNS immune surveillance and inflammation. Nat. Commun. 14, 1–23 (2023).

27. Pietilä, R. et al. Molecular anatomy of adult mouse leptomeninges. Neuron 111, 3745–3764.e7 (2023).

28. Neuhuber, W. An “outer subarachnoid space”: fact or artifact? Fluids Barriers CNS 21, 1–2 (2024).

29. Betsholtz, C. et al. Advances and controversies in meningeal biology. Nat. Neurosci. 27, 2056–2072 (2024).

30. Liu, S. et al. Fluid outflow in the rat spinal cord: the role of perivascular and paravascular pathways. Fluids Barriers CNS 15, 1–14 (2018).

31. Walsh C. L., et al. Imaging intact human organs with local resolution of cellular structures using hierarchical phase-contrast tomography. Nat. Methods 18, 1532–1541 (2021). 10.1038/s41592-021-01317-x

32. Does the brain really have a fourth meningeal membrane? APS Division of Fluid Dynamics 77th Annual Meeting https://meetings.aps.org/Meeting/DFD24/Session/L06.5 (2024).

33. Shlobin, N. A. et al. The glymphatic system and subarachnoid lymphatic-like membrane: recent developments in cerebrospinal fluid research. World Neurosurg. 190, 147–156 (2024).

34. Kipnis, J. et al. Resolving the mysteries of brain clearance and immune surveillance. Neuron 113, 3908–3923 (2025).

